# Targeted mutagenesis of specific genomic DNA sequences in animals for the in vivo generation of variant libraries

**DOI:** 10.1101/2024.06.10.598328

**Authors:** Julia Falo-Sanjuan, Yuliana Diaz-Tirado, Ting-Wei Pai, Meghan A. Turner, Alejandro Collins, Julian Davis, Jialing Fang, Claudia Medrano, Olivia Rourke, Peter F. Bailer, Jenna Haines, Joey McKenna, Arman Karshenas, Rahul M. Kohli, Rob Phillips, Michael B. Eisen, Hernan G. Garcia

## Abstract

To uncover the rules by which the number, placement and affinity of transcription factor binding sites dictate gene regulatory programs, it is first necessary to find the binding sites within regulatory regions. Massively parallel reporter assays (MPRAs), in which gene expression driven by tens of thousands of in vitro-generated regulatory sequences is measured, have made this feat possible at high-throughput in single cells in culture. However, because of lack of technologies to incorporate DNA libraries in multicellular organisms, MPRAs have remained limited in whole animals and plants. To overcome this limitation, we developed an in vivo mutagenesis approach in the fruit fly *Drosophila melanogaster* based on targeted base editing. In this system, mutator parents generate populations of offspring, each carrying a distinct variant of a defined genomic region, forming a library of variants we term Flybraries. Through multiple rounds of optimization, we determine that nCas9 fused to a sea lamprey base editor (evoCDA1) guided by arrays of up to 30 gRNAs efficiently introduces mutations across a 200-500 bp sequence in cell culture and in vivo. Applying this approach to the *stripe 2* enhancer of the *even-skipped* gene, we show that induced mutations lead to changes in gene expression consistent with perturbation of known binding sites. Thus, we show that diversifying base editors and arrays of gRNAs is a promising and scalable tool to generate regulatory sequence diversity in vivo, enabling MPRA-like dissection of enhancer function multicellular organisms.

## Introduction

The orchestrated regulation of gene expression—in time, space, or in response to signals—is required for most aspects of life, such as ensuring the correct development of body plans, maintaining homeostasis and regeneration (Howard and Davidson, 2004; Levine, 2010; Rubinstein and de Souza, 2013). Misregulation of gene expression has been increasingly linked to disease, including cancer, where non-coding and regulatory mutations contribute substantially to pathogenesis (Cuykendall et al., 2017; Rheinbay et al., 2020). Among the many layers to gene regulation in eukaryotes, short DNA regulatory sequences termed enhancers play a major role in determining the levels and spatiotemporal pattern of the expression of a gene. Enhancers are bound by transcription factors in 6-10 bp DNA motifs termed transcription factor binding sites. Differences in the transcription factor binding site sequence, number, orientation and spacing, amongst other features, are thought to relate to transcription factor binding strength and dynamics and, in turn, to their interaction with the transcriptional machinery (Barolo and Posakony, 2002; Lelli et al., 2012; Spitz and Furlong, 2012).

Despite the clear role of enhancers in driving gene expression programs, the rules by which the number, placement and affinity of binding sites within them dictate transcriptional dynamics remain unknown. To uncover these rules, however, it is first necessary to find the binding sites within an enhancer with high precision, and then to measure how systematic modulation of this binding site arrangement controls gene expression.

Massively parallel approaches for mapping regulatory sequence variants to gene expression have made it possible to identify transcription factor binding sites at high throughput. The most widely used implementation of this strategy in the context of cells in culture and in some multicellular organisms is the massively parallel reporter assay (MPRA) (Arnold et al., 2013; Inoue and Ahituv, 2015; Ireland et al., 2020; Kheradpour et al., 2013; Kinney et al., 2010). Here, a barcoded library of reporter genes, each driven by a mutated regulatory region or genome fragment, is generated in vitro and transfected into cells. RNA sequencing is then used to correlate enhancer sequence and gene expression, revealing activator and repressor binding sites (Ireland et al., 2020). Further, once binding sites are identified, libraries can be computationally designed to systematically modulate regulatory parameters such as the relative placement of binding sites (de Almeida et al., 2022; Qi et al., 2020; Sharon et al., 2012; Yu et al., 2021). These approaches have resulted in a pipeline that allows for the rapid identification and experimental validation of binding sites within an enhancer, and for the engagement in a theory–experiment dialogue aimed at reaching a predictive understanding of how binding site architecture dictates gene expression (Pan et al., 2024; Tareen et al., 2022).

Despite these advances, extending MPRA approaches to multicellular organisms requires overcoming three key challenges. First, large libraries of animals must be generated, each carrying a unique set of mutations within a targeted regulatory region. Second, those mutations must lead to measurable phenotypic differences, such as changes in gene expression patterns during development. Third, the mutated regulatory sequence present in each individual must be linked to its corresponding phenotype in order to correlate sequence variation with functional output. Only by satisfying these three requirements can sequence–phenotype relationships be obtained, enabling the inference of regulatory mechanisms.

Despite their promise, the application of MPRAs in animals remains constrained in two fundamental ways. First, because MPRAs rely on the incorporation of in vitro–generated DNA libraries, they can only be deployed in systems amenable to transfection or viral delivery. As a result, with some notable exceptions—-including adult mouse brains, *C. elegans*, *Ciona intestinalis* and tobacco leaves (Chan et al., 2023; Farley et al., 2015; Jores et al., 2020; Lagunas et al., 2023; Ren et al., 2026)— the field lacks a reliable pipeline to incorporate DNA diversity into most animal and plant model organisms. For example, in the fruit fly the number of variants tested in stable transgenic lines is typically on the order of 40 per study (Falo-Sanjuan et al., 2019; Ip et al., 1992; Sayal et al., 2016; Swanson et al., 2010) with a notable exception testing ∼750 variants over multiple years (Fuqua et al., 2020).

Second, even when MPRAs are implemented, they typically rely on reporter constructs in which enhancers are assayed outside their endogenous genomic context (Chan et al., 2023; Farley et al., 2015; Kheradpour et al., 2013; Lagunas et al., 2023). These reporter constructs are often episomal and lack native chromatin organization, limiting their ability to capture regulatory effects such as local chromatin state or histone modifications. Recent advances in tile editing have allowed the introduction of mutations in endogenous genomic loci in cell culture (Becerra et al., 2024; Roh et al., 2025). While it has been possible to xenograft edited cells into living mice (Ren et al., 2026), in vivo endogenous editing has not yet been feasible.

Here we develop an approach that enables massively parallel sequence–function mapping to endogenous regulatory loci in vivo. This approach overcomes the key limitations that have prevented the application of MPRAs in multicellular organisms and at endogenous loci. As an alternative to generating enhancer libraries in vitro and incorporating them into organisms, we implemented a system that directly induces random mutations in vivo at targeted genomic loci, using the fruit fly *Drosophila melanogaster* as a case study. Here, mutator flies generate libraries of enhancer variants—”Flybraries”—in their germline cells. Each embryo inherits a unique realization of the mutated enhancer, enabling direct correlation of sequence and transcriptional activity. Of note, throughout this paper we refer to “in vivo*”* to indicate mutations within a multicellular organism, as opposed to cultured cells, used for system optimization.

To enable in vivo mutagenesis in the genome, we assayed multiple DNA editing tools such as base editors and error-prone DNA polymerases. Most notably, inspired by tiled base editing approaches in cell culture (Becerra et al., 2024; Ravi et al., 2022), we used arrays of gRNAs to target nickase Cas9 (nCas9) fused to deaminases derived from the somatic hypermutation enzyme AID to stochastically produce cytosine to thymine (C→T) transitions (Berríos et al., 2024; Kohli et al., 2021). Through iterative optimization in *Drosophila* animals and cell culture, combined with mathematical modeling of the mutational process, we identify the sea lamprey AID evoCDA1 as a highly efficient mutator in vivo, in contrast to commonly used mammalian variants (Anzalone et al., 2020; Doll et al., 2023; Thuronyi et al., 2019). Further, we optimize the spatial window of mutagenesis by modulating the length, number and overlap of guide RNAs (gRNA) that mediate the targeting of this fusion to the DNA. Finally, we generate enhancer variants in vivo, measure their effect on gene expression in high throughput, and directly link sequence variation and to gene expression output in order to draw mechanistic insights about the regulatory logic of the targeted enhancer. Together, these advances establish a framework for MPRA-like dissection of endogenous regulatory loci in multicellular organisms.

## Results

### Mammalian AID–nCas9 base editors are active in cell culture but fail to induce mutations in vivo

In the last few years, multiple tools have been developed for targeted mutagenesis such as prime editing, base editing, and the error-prone polymerase system EvolvR. However, prime editing requires a pegRNA encoding each specific desired mutation (Bosch et al., 2020), such that generating a library of variants would require a large number of distinct pegRNAs, one per mutation. Because such pegRNA libraries cannot be efficiently introduced into most multicellular organisms, this approach is not suitable for high-throughput mutagenesis in vivo.

In contrast to prime editing, base editing and EvolvR can, in principle, be used for high-throughput mutagenesis in animals, as they do not require pre-specifying individual mutations. Adenine deaminases convert A→I, resulting in A→G mutations (and T→C on the antisense strand), while cytosine deaminases that convert C→U, resulting in C→T mutations (and G→A on the antisense strand). Both kinds of deaminases have been engineered for high efficiency and precision with the goal of reverting disease-causing mutations when targeted using fusions to nCas9 (Anzalone et al., 2020; Fu et al., 2021; Newby et al., 2021; Thuronyi et al., 2019). Importantly, despite their limited mutational spectra, recent theoretical work suggests that mutations introduced by adenine and cytosine deaminases are still sufficient for the dissection of regulatory regions such as identifying the binding sites within it (Pan et al., 2024). Finally, to minimize indel formation, many base editors include a uracil glycosylase inhibitor (UGI), which suppresses base excision repair (Wang et al., 2017).

Further, the EvolvR system relies on fusing enCas9 (enhanced-nickase Cas9) to an engineered error prone *E. coli* DNA polymerase I with an error rate 12,000 fold compared to wild type (Figure 1 - figure supplement 1). The mutational spectrum of EvolvR is not as restricted as that of base editing: it can mutate all bases to similar degrees (Tou et al., 2020).

**Figure 1.**
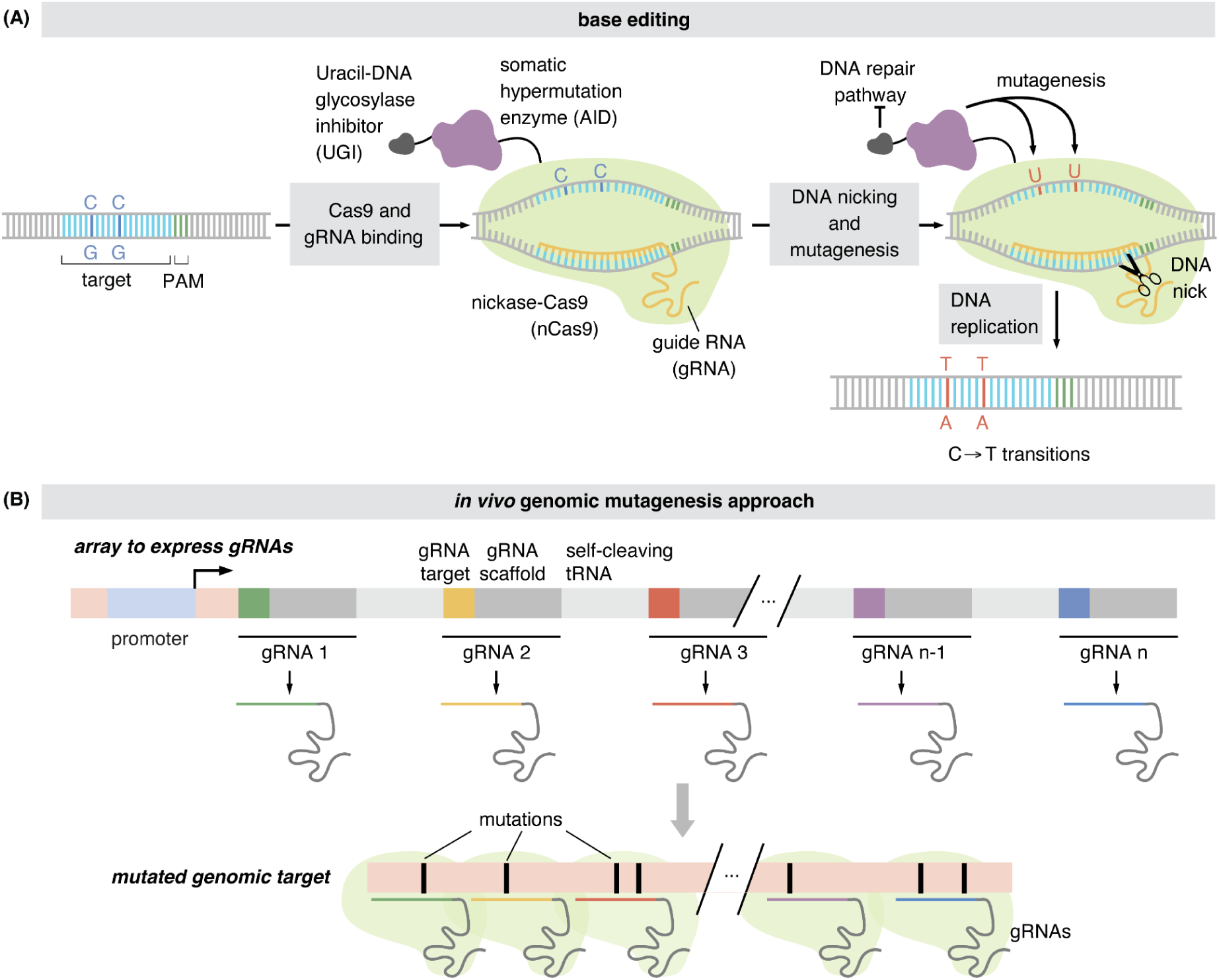
Tools to induce mutations in vivo. **(A)** In base editing, (i) a guide RNA (gRNA) is used to target nickase Cas9 (nCas9) to a specific DNA sequence. Importantly, targetable DNA sequences are constrained to those next to a protospacer adjacent motif (PAM). (ii) Upon nCas9 binding, AID fused to this protein mutates C→T on the strand not bound by the gRNA. The DNA nick introduced by nCas9 increases the efficiency of mutagenesis. (iii) Uracil glycosylase inhibitor (UGI) inhibits uracil excision during DNA repair, thereby avoiding indels**. (B)** To mutagenize a long stretch of DNA, multiple gRNAs are expressed in a single RNA transcript flanked by tRNAs for self cleaving of each individual gRNA. Different colors indicate the different protospacers targeting DNA and dark gray lines indicate the shared gRNA scaffolds.

We tested these two mutagenesis approaches. First, we examined several variants of human AID (AID12* or AID123*), fused to nCas9, with or without uracil glycosylase inhibitor (Berríos et al., 2024; Kohli et al., 2021) (Figure 1A). Second, we tested the EvolvR system (Tou et al., 2020) (Figure 1 - figure supplement 1). To mutagenize extended genome sequences, and inspired by approaches to fluorescently label loci by recruiting Cas9 proteins gRNAs arrays (Gu et al., 2018), we expressed arrays of gRNAs targeting a single locus, enabling the simultaneous recruitment of multiple nCas9–AID or nCas9–EvolvR complexes across the region (Figure 1B).

To test whether multiple gRNAs can drive mutagenesis across a genomic region, we generated a construct expressing four gRNAs targeting GFP. The gRNAs were arranged in a single transcript under a U6 promoter and flanked by tRNA sequences to enable their processing into individual guides. This 4xgRNA plasmid was transfected into *Drosophila* Kc cell lines expressing GFP off of the chromosome (Kc-NLS-GFP, (Cherbas et al., 2015)), along with plasmids expressing the different AID-nCas9 and EvolvR-enCas9 variants. These variants were driven by distinct promoters to achieve different expression levels: the ubiquitous *act5C* or the Cu^2+^ inducible *pMT* promoters (Figure 2A, Figure 2 - figure supplement 1, see Methods - Cell Culture) (Bunch et al., 1998).

**Figure 2.**
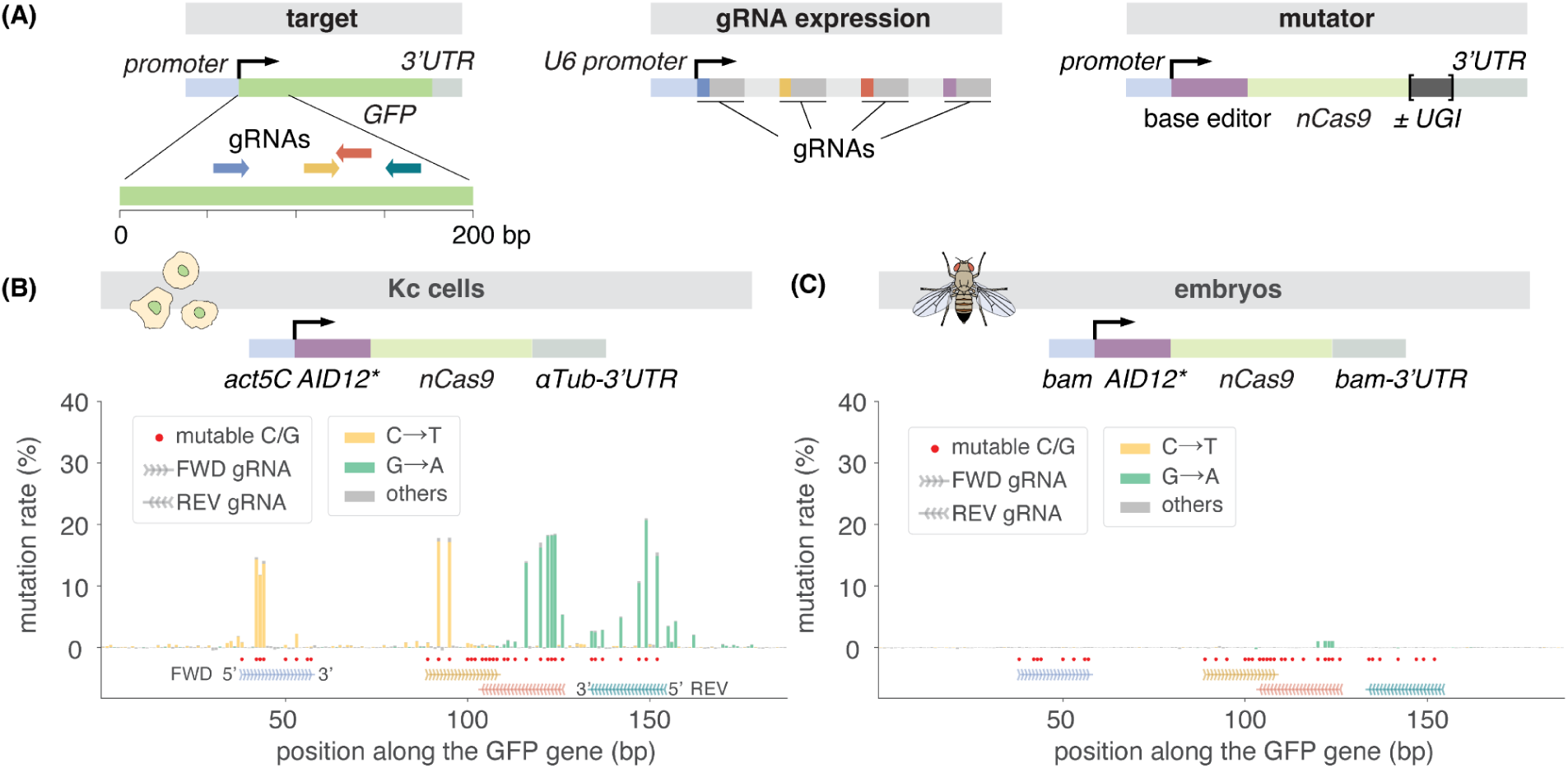
Human AID variants introduce C→T transitions at high rates at gRNA target sites in cell culture, but not in animals. **(A)** Plasmids used to mutagenize a segment of the GFP sequence in the genome of Kc cells or fly embryos. Four gRNAs were used to target GFP with nCas9 fused to mutagenic proteins AID or EvolvR. Different AID variants, promoters and 3’UTRs were tested. gRNAs were expressed as an array from a U6 ubiquitous promoter. **(B)** Barplots showing mismatches to the targeted GFP segment classified by base substitution generated by AID12* in Kc-NLS-GFP cells four weeks post-transfection. Most mismatches are C→T (yellow) on forward gRNAs or G→A (green) on reverse gRNAs. See Figure 2 - figure supplement 2 for other AID variants and promoter conditions. **(C)** Barplots showing mismatches to GFP classified by base substitution generated by AID12* in F2 embryos. Just a few G→A (green) mutations were detected. In (B) and (D), red dots mark all “mutable bases” (Cs on FWD gRNAs and Gs on REV gRNAs).

To assess the extent of mutagenesis at the chromosomal GFP locus in cultured cells, we extracted genomic DNA at multiple timepoints after transfection and performed amplicon sequencing of GFP using Nanopore technology. Mismatches to the reference sequence were detected across all AID samples and timepoints, with AID12* at 4 weeks post-transfection producing the highest rate, reaching ∼20% at some gRNA-targeted positions within GFP (Figure 2B, Figure 2 - figure supplement 2). The observed mutations were C→T in forward (FWD) gRNAs binding to the ‘minus’ strand, and G→A mutations in reverse (REV) gRNAs binding to the ‘plus’ strand. This mutational profile is consistent with AID almost exclusively being able to mutate the strand of DNA not bound by the gRNA. Mutations were biased towards the 5’ half of the gRNAs, with occasional edits 1–3 bp upstream (i.e., 5’) of the protospacer (Figure 2 - figure supplement 3), consistent with previous observations (Kohli et al., 2021; Marr and Potter, 2021). Notably, very few indels were detected even in samples without UGI (Figure 2 - figure supplement 2B), consistent with recent reports that, unlike hemimetabolous insects, flies do not have the UNG gene required for the uracil excision repair pathway (Doll et al., 2023). No mutations were detected in EvolvR samples (Figure 2 - figure supplement 2B).

Having established that AID-nCas9 induces mutations in *Drosophila* cell culture, we next tested its activity in *Drosophila* embryos. To do so, it was important to control the timing of the mutagenesis: mutations generated during the embryonic blastoderm stage could generate mosaic embryos, while somatic mutations in adults might cause toxicity and fertility loss. Further, mutations occurring early on in the germline could become fixed in the parent flies and all offspring, reducing library diversity. We therefore restricted mutagenesis to oogenesis or spermatogenesis by ensuring that AID is expressed only after germline stem cell division throughout gametogenesis. To make this possible, we expressed the AID-nCas9 variants using the *bag of marbles* (*bam)* promoter, which is only active in the germline cyst stage following germ stem cell division in both males and females (Chen et al., 2020).

We targeted a genomic GFP in the fly genome using the same four gRNAs as in our cell culture experiment and sequenced GFP from F2 embryos obtained from parents expressing both AID-nCas9 or EvolvR-enCas9 and 4xgRNAs. Mutagenesis was minimal in vivo: No mutations were detected for AID123*, and AID12* produced mutations at frequencies below 1% at only a few positions (Figure 2C). Further, similarly to cell culture, no mutations were detected in the EvolvR samples (Figure 2 - figure supplement 4).

The observed in vivo mutation rates of at most 1% are far below the 10% mutation rate necessary to recover information such as binding site locations in bacterial MPRAs (Ireland et al., 2020; Pan et al., 2024). As a result, because AID performed well in cell culture, we focused on this approach as opposed to EvolvR and sought to optimize base editing to increase mutation rates in vivo.

### Lamprey evoCDA1 is highly mutagenic in vivo

We explored several potential causes for the observed discrepancy in AID–nCas9 mutagenic activity between cell culture and flies. First, to ensure that the gRNAs were correctly expressed and functional in flies, we crossed our fly line expressing the four gRNAs targeting GFP with a line expressing catalytically active Cas9. We obtained a high rate of indels (100% in some positions, Figure 3 - figure supplement 1A), indicating that the gRNAs are expressed and can successfully target the GFP gene.

**Figure 3.**
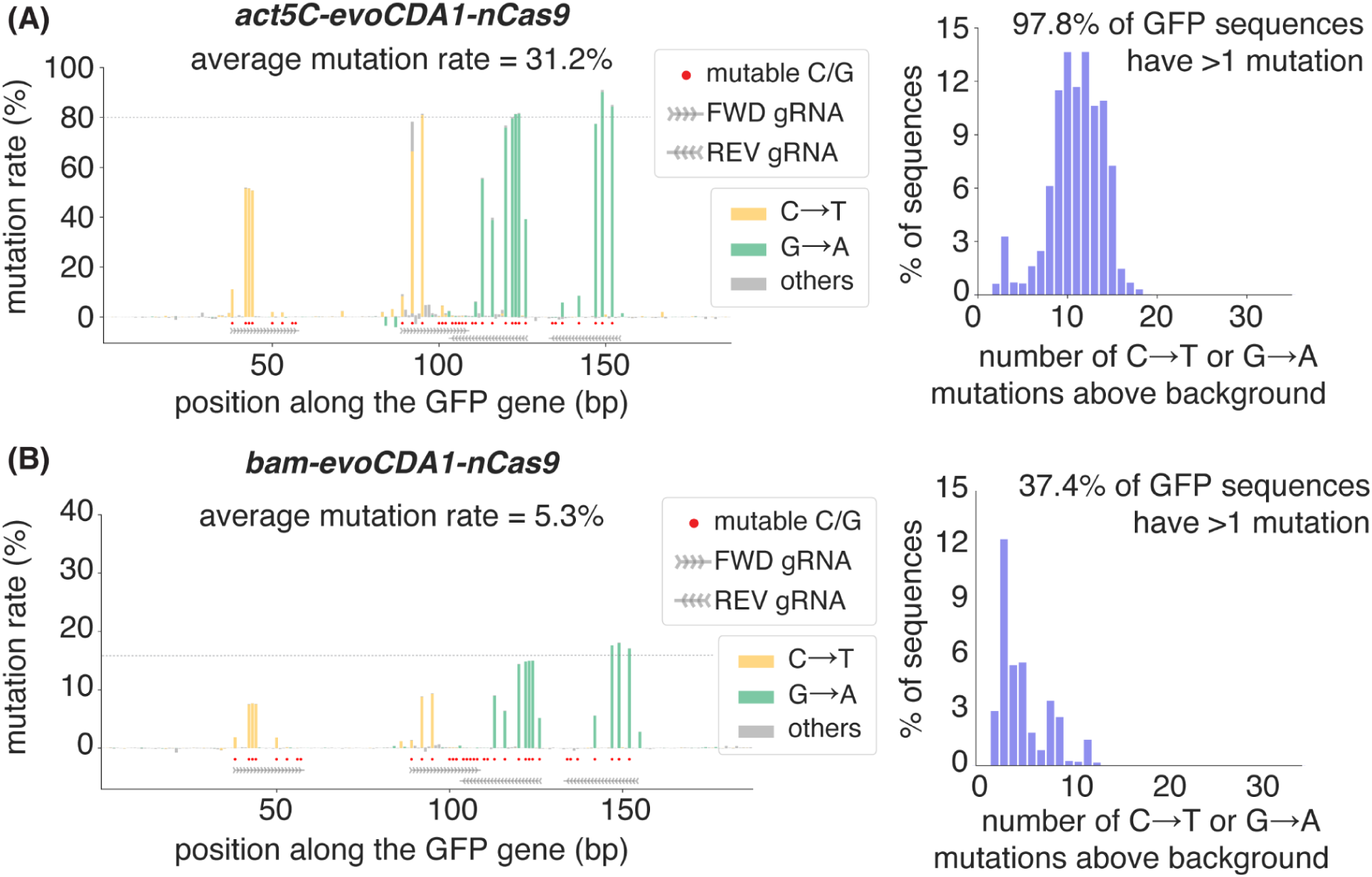
Sea lamprey evoCDA1 introduces C→T transitions at high rates in embryos. **(A)** Mismatches in GFP classified by base substitution in adults where *act5C*-evoCDA1-nCas9 introduced mutations from the paternal germline (left), and distribution of the observed number of mutations per read after subtracting the number of mismatches observed in control samples (right). Red dots mark all “mutable bases” (Cs on FWD gRNAs and Gs on REV gRNAs). **(B)** Mismatches in GFP classified by base substitution in adults where *bam*-evoCDA1-nCas9 introduced mutations from the paternal germline (left), and distribution of the observed number of mutations per read after subtracting the number of mismatches observed in control samples (right). Red dots mark all “mutable bases” (Cs on FWD gRNAs and Gs on REV gRNAs).

Second, because fly experiments are performed at 25°C and fly cell culture experiments at 27°C, it is possible that the difference in mutation rate we observed is due to temperature sensitivity of AID activity, as it has been recently reported for some AID variants (Dancyger et al., 2012; Doll et al., 2023). To determine the effect of temperature on in vivo mutagenesis, we repeated our experiments at 29°C—towards the high range of fly viability. However, we saw no significant difference in mutation rate (Figure 3 - figure supplement 1B, compare to Figure 2 - figure supplement 4), suggesting that temperature is not the key factor dictating AID functionality.

Third, cells were mutated for up to 4 weeks, while the *bam* promoter is only active for four days during gametogenesis in the animal (Rust et al., 2020). To determine whether more mutations could be accumulated over time in the fly genome, we created stable stocks with each of our human AID variants and let these variants mutate for up to 10 generations, corresponding to a total mutation period of ∼40 days. We collected embryos and sequenced GFP at generations 4, 6, 8 and 10. Although more mutations were detected over time (Figure 3 - figure supplement 2, Figure 3 - figure supplement 3B), the maximum mutation rate detected was 2%, and the average mutation rate of 0.3% across all mutable base pairs was still significantly below the 10% that would be needed to extract regulatory information from an enhancer.

Lastly, we explored whether differences in AID expression levels could explain the difference in mutagenesis rate between our cell culture and in vivo experiments. To achieve higher levels of AID expression, we created transgenic flies where the *bam* promoter was replaced with the *act5C* promoter, which we had used in cell culture. Further, we also replaced the *bam* promoter with the strong germline UASz promoter, and drove this promoter with either *bam-Gal4* or *osk-Gal4* (Figure 3 - figure supplement 3) (Deluca and Spradling, 2018). However, none of the *act5C* or *UASz* versions of our AIDs constructs produced any significant increase in mutation rate (Figure 3 - figure supplement 3C).

Our inability to to generate significant mutations in the fly using widespread human AID variants led us to explore variants from other species. Specifically, we used the *act5C* promoter to express three other AID/APOBEC variants, two from rat (rAPOBEC1 and evoAPOBEC1) and one from sea lamprey (evoCDA1) (Figure 3 - figure supplement 3A), all of which were recently implemented in *Drosophila* (Doll et al., 2023; Marr and Potter, 2021). While the rat APOBEC1 variant did not result in significant mutations, the two in vitro evolved evoAPOBEC1 and evoCDA1 variants exhibited a significant increase in mutation rate. Strikingly, evoCDA1, the AID variant evolved from the lamprey *Petromyzon marinus* pmCDA1 (Thuronyi et al., 2019), led to a remarkable amount of mutagenesis in sequenced embryos (Figure 3 - figure supplement 3C, 19% across all mutable base pairs).

Given the high mutation rate of the lamprey AID, we sought to further characterize the mutagenesis induced by this deaminase. First, since, unlike the *bam* promoter, the *act5C* promoter we used to express this AID is active in all cells, we determined to what degree mutations were produced in the somatic or germline tissues of the mutagenic parents. To make this possible, we set up crosses that allowed us to distinguish flies where mutations could only have occurred in the soma or germline (see Methods - Fly Strains and Genetics). This experiment confirmed that mutations were indeed occurring in the germline to a similar degree as soma, suggesting that each of our embryos is isogenic. Further, our results show that mutations are twice as likely to occur in the paternal germline, reaching 80% in some positions and 98% reads harboring at least one mutation (Figure 3A), compared to 40% and 73%, respectively, for mutations from the maternal germline (Figure 3 - figure supplement 4A).

The observed difference in mutation rate between males and females could stem from different sources. First, sex-specific differences in AID efficiency or *act5C* expression could exist between males and females. Second, the fact that male germline cells go through six divisions rather than four in the female germline to generate gametes (Hempel et al., 2008) could mean that AID has more time to act on males than females. Regardless of the source of this sex-specific difference, our results suggest that mutations generated in either germline are high enough to constitute a suitable strategy for modulating mutation efficiency.

As shown above, the sea lamprey evoCDA1, which we refer to as AID for the remainder of the paper, was remarkably efficient in introducing mutations in a short period of time. However, mutation rates between 40% and 80% at many positions (Figure 3A) could be too high for our purposes: if too many mutations are introduced in a given enhancer it will not be possible to clearly discern which position(s) along the sequence are important for the modulation of expression levels (Pan et al., 2024). Replacing the *act5C* promoter with the less active *bam* promoter resulted in a mutation rate of 5.3% over all mutable base pairs from the paternal germline, with maximum rates of 15% at certain positions (Figure 3B, Figure 4 - figure supplement 4B). However, these mutation rates could also change as we go beyond mutagenizing a short GFP segment and deploy many gRNAs in order to generate libraries of enhancer sequences. We therefore spend the next two sections further investigating the sequence diversity generated by AID driven by the *act5C* and *bam* promoters, and how they perform when the number of gRNAs increases.

**Figure 4.**
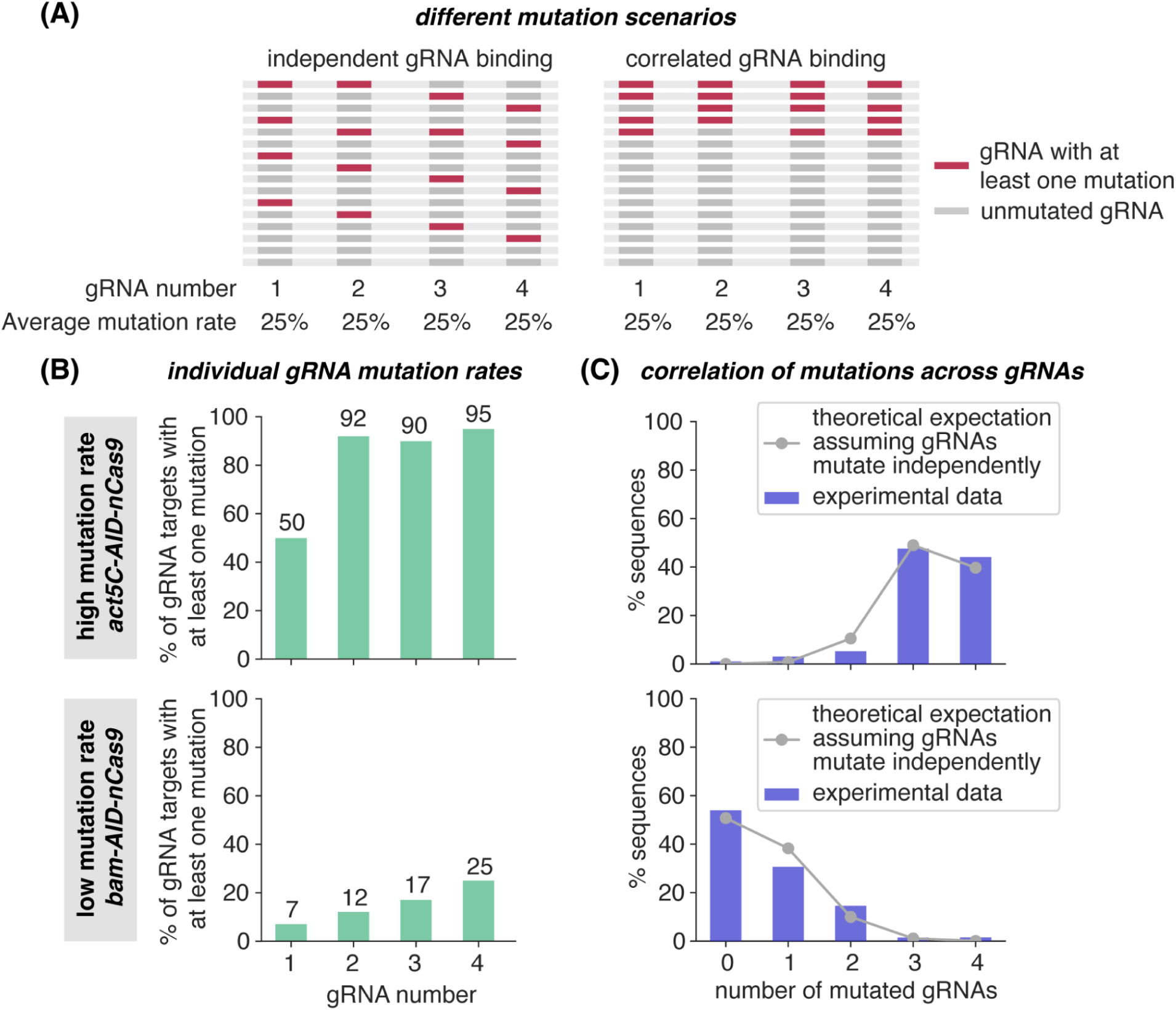
Quantifying the variability in mutations generated by AID and four gRNAs in embryos. **(A)** Two different scenarios for correlation between mutations generated by different gRNAs: completely uncorrelated and correlated. **(B)** Average mutation rates in each gRNA, defined as the percentage of reads with at least one mutation within the gRNA target sequence, for embryos expressing gRNAs and *act5C-AID-nCas9-UGI* (top) or *bam-AID-nCas9-UGI* (bottom). **(C)** Quantification of the correlation in mutations across gRNAs for the *act5C* (top) and *bam* (bottom) constructs. Line plots show the expected distribution of the number of mutated gRNAs per read assuming mutation rates from (A) and that each gRNA mutates a DNA molecule independently of each other (Poisson Binomial Distribution, see Methods - Quantification Mutational Variability). Blue bars indicate the measured number of mutated gRNAs per read.

### Quantitative analysis reveals that gRNAs mutate independently of each other in embryos

To maximize sequence diversity across embryos, mutations must arise independently in each germ cell rather than being shared across lineages. However, promoter choice could introduce correlations between mutations. For example, while the *bam* promoter is only active in later stages of gamete development, the *act5C* promoter is active in the stem cell stage during germline development. As a result, mutations driven by *act5C-*AID could become fixed in the germline progenitors and inherited by many offspring, producing embryos with identical or very similar sequences. Thus, while *act5C-AID* mutates at a higher rate, its early activity could reduce the overall sequence diversity in the resulting animal libraries.

To determine and compare the degree of sequence diversity generated by *act5C-AID* and *bam-AID*, it is necessary to assess the degree of correlation between mutations occurring within a given gRNA and between different gRNAs. To make this possible, we went beyond the low accuracy Nanopore sequencing used to characterize *average* mutation rate and re-sequenced our libraries using higher accuracy Illumina technology.

To determine whether gRNAs mutated on the same DNA molecule independently of each other, we compared the measured distribution of the number of mutated gRNAs per read to a simple theoretical model that assumes no correlations in mutagenesis between gRNAs (Figure 4A). We first measured the fraction of gRNA targets with at least one mutation (Figure 4B). We then used these mutation rates to predict the probability of obtaining reads containing 1, 2, 3 or 4 mutated gRNAs (see Methods - Quantification of Mutational Variability). A comparison of this theoretical expectation to our data for low (*bam*) and high (*act5C*) mutation rates shows an agreement between theory and experiment, suggesting that, indeed, each gRNA mutates independently of each other (Figure 4C).

To quantify correlations using a complementary approach, we calculated the normalized pointwise mutual information (NPMI; see Methods) between pairs of gRNAs (Figure 4—figure supplement 1A), as well as between mutable bases within each gRNA (Figure 4—figure supplement 1B). Across gRNAs, most pairs showed little to no correlation, suggesting that gRNA binding events occur largely independently. When considering mutations within individual gRNAs, many mutations co-occurred in the same DNA molecule, especially at the high (*act5C*) mutation rate. However, the NPMI analysis showed that most mutable bases were also uncorrelated within individual gRNAs. This suggests that mutations at different positions could arise independently despite co-occurring in the same DNA molecule. An exception to this lack of correlation was observed for gRNAs 2 and 3, which exhibited positive correlations both between each other and within their mutable bases. Because these 2 gRNAs overlap, it is possible that binding of one facilitates the binding of the other—for example by lowering the energy required to open the bubble of ssDNA (Corsi et al., 2022)—thereby facilitating access of AID to nearby bases. Thus, our two quantification methods indicate that mutations across and within gRNAs are largely uncorrelated, with a potential positive correlation when gRNAs overlap. Notably, similar correlation patterns were observed for both low (*bam*) and high (*act5C*) mutation rates, suggesting that both constructs will lead to similar degrees of mutational diversity in an enhancer when targeted with many gRNAs.

### gRNA arrays enable uniform mutagenesis across extended genomic regions in cell culture

It is important to note that, while our initial experiments were conducted using only four gRNAs, extending this approach to mutagenize full enhancers requires tiling many guides across hundreds of base pairs to achieve uniform mutation rates approaching 10% while maintaining uniform coverage along the entire sequence. This dense tiling inevitably leads to overlapping gRNA target sites. Because mutations disrupt gRNA binding, edits introduced by one guide can prevent subsequent binding of overlapping guides. Effectively, overlaps could render whole stretches of DNA refractory to further mutations. Thus, increasing gRNA density introduces a potential tradeoff between coverage and achievable mutation rate.

To explore how gRNA overlap informs our design strategy for blanketing an enhancer, we developed a simple model of AID–nCas9 mutagenesis that predicts how mutations accumulate over time and how edited DNA becomes refractory to further targeting. Briefly, our model posits that each nCas9 molecule binds a gRNA target in each cell cycle with a probability *p_bound_*. Once bound, AID mutates each available C or G (depending on whether the gRNA is FWD or REV) with a probability *p_mut_* (Figure 5A). Mutations that arise in one cell cycle prevent gRNA binding in subsequent cell cycles, preventing further edits at overlapping sites.

**Figure 5.**
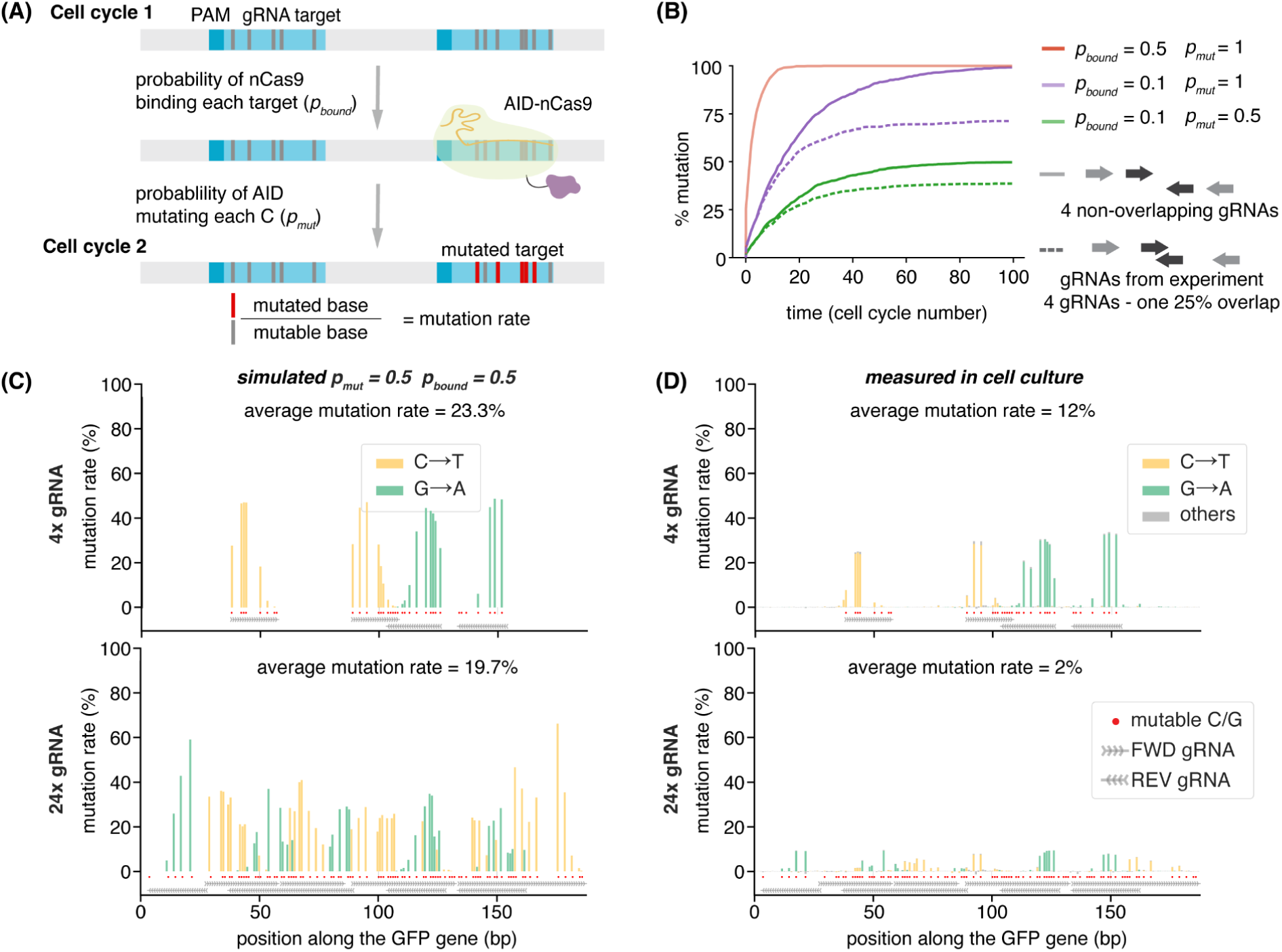
Simulating the expected number and distribution of mutations. **(A)** Simple model of mutations caused by AID-nCas9 in two probabilistic steps. First, each gRNA guides the binding of nCas9 to an intact target site with probability *p_bound_*per cell cycle. Upon binding, AID mutates each of available C (or G in REV gRNAs) with a probability *p_mut_*. After mutagenesis, the *p_bound_*of any gRNA overlapping the mutated sequence becomes zero. The mutation rate is defined as the mean fraction of mutated bases over all the mutable bases, averaged over multiple simulated DNA molecules. **(B)** Average fraction of mutations in all mutable bases at each generation when four non-overlapping gRNAs with uniform mutation rates target GFP under different simulation conditions: *p_bound_ =* 0.5 and *p_mut_* = 1 (orange), *p_bound_ =* 0.1 and *p_mut_* = 1 (purple solid), *p_bound_ =* 0.1 and *p_mut_* = 0.5 (green solid) or with 4 gRNAs where there is one 25% overlap equivalent to those used in experiments *p_bound_ =* 0.1 and *p_mut_*= 1 (purple dashed), *p_bound_ =* 0.1 and *p_mut_* = 0.5 (green dashed). Simulations consist of 200 cells mutating over 100 cell cycles. **(C)** Predicted mutation profile when the GFP fragment is targeted with 4 gRNAs (top) or 24 gRNAs (bottom,) using *p_bound_ =* 0.5 and *p_mut_* = 0.5. Average mutation rate in each position is calculated over 1000 simulated mutated DNA molecules mutating over 10 cell cycles to mimic the number of generations that would have occurred in cell culture after two weeks as in our experiments. **(D)** Measured mutational profile of GFP classified by base substitution in Kc cells transfected with lamprey *act5C-AID-nCas9* and 4 (top) or 24 (bottom) gRNAs, 2 weeks post-transfection. To experimentally test the hypothesis that gRNA overlaps lead to a small reduction in mutation rate, we cloned arrays of gRNAs containing these 24 gRNAs and assessed both the overall mutation rate and its distribution along the targeted sequence. Because of the challenge of cloning a single repetitive array, we distributed the gRNAs across three different plasmids (containing 6, 6 and 12 gRNAs). In cell culture, we co-transfected these arrays of 24 gRNAs with *act5C-AID-nCas9* and compared mutagenesis to the 4 gRNA condition.

To build intuition for our model, we simulated the effects of different *p_bound_* and *p_mut_* values, and of gRNA overlap, on mutagenesis over many cell cycles. We first tested this model for the case of our 4 non-overlapping gRNAs targeting GFP and incorporated the 5’ mutation bias observed in each gRNA (Figure 2 - figure supplement 3). We simulated thousands of cells mutating over multiple cell cycles, and calculated the average mutation rate over time, defined as the mean fraction of mutated bases over all the mutable bases after each cell cycle (Figure 5B). As expected, the fraction of mutated bases saturates after a certain number of cycles (Figure 5B, Figure 5 - figure supplement 1A). By systematically modulating parameters, we determined that *p_mut_*sets the maximum achievable mutation rate while the *p_bound_* controls the time to reach this maximum (Figure 5B, Figure 5 - figure supplement 1A). Importantly, introducing overlaps between gRNA molecules reduces the maximum mutation rate, with stronger effects at high *p_mut_* (Figure 5B, Figure 5 - figure supplement 1B). Together, these results confirm gRNA overlap as an important constraint on mutagenesis efficiency and provide a quantitative framework to guide the design of gRNA arrays for uniformly mutating extended genomic regions.

To more realistically model the mutagenesis of a longer DNA sequence such as an enhancer, we designed gRNAs for all possible protospacer adjacent motif (PAM) sites required for Cas9 binding in a 200 bp sequence of GFP. We then removed those gRNAs that were almost identical, ending up with 24 gRNAs that cover the whole 200 bp sequence in FWD and REV (Figure 5 - figure supplement 1C). In agreement with the observation above that overlaps have a bigger impact at high *p_mut_*, the mutation rates with 24 gRNAs remained similar or slightly decreased compared to 4 gRNAs (Figure 5 - figure supplement 1D). Examples of mutational profiles obtained are shown in Figure 5C.

In the 24 gRNA condition, we detected mutations along the 200 bp region. Our measurements revealed a substantially more uniform mutation profile, with reduced 5’ bias compared to the 4 gRNA condition (compare top and bottom in Figure 5D; Figure 5 - figure supplement 2). This improved uniformity indicates that gRNA arrays can effectively tile extended genomic regions and approach the goal of evenly distributed mutagenesis across an enhancer. However, despite this improved coverage, the overall mutation rates were lower than predicted by our simulations. This discrepancy suggests that factors beyond gRNA overlap—such as limiting concentrations of nCas9-AID or gRNAs—may further constrain mutagenesis efficiency when many guides are expressed simultaneously.

Finally, we sought to optimize another parameter: the gRNA length. Although standard CRISPR uses gRNAs 20 bp in length (Jinek et al., 2013), we noticed that one of the four gRNAs was inadvertently 23 bp long (Ma et al., 2016). This longer gRNA showed higher mutagenic activity, extending into these three additional bases. To systematically determine the effect of gRNA length, we generated plasmids expressing either the 20 or 23 bp version of the same gRNA and transfected these plasmids together with *act5C-AID-nCas9* in cell culture. Confirming our hypothesis, the 23 bp gRNA indeed extended the mutation window 5’ with respect to the 20 bp gRNA (Figure 5 - figure supplement 3A). We next asked whether further increasing gRNA length could expand this mutagenic window and reduce the number required to tile a whole enhancer. However, we found that increasing the gRNA length beyond 23 bp actually decreased the mutagenesis efficiency and restored the narrower editing window of 20 bp guides (Figure 5 - figure supplement 3B). Thus, we concluded that 23 bp is the optimal balance between gRNA editing window and efficiency in cell culture.

Together, our simulations and cell culture experiments demonstrate that gRNA arrays can achieve broad and relatively uniform mutagenesis across extended DNA sequences. We next asked whether this behavior extends to in vivo conditions and which promoter enables mutation rates closest to our target of ∼10%.

### In vivo mutagenesis of the *eve2* enhancer identifies optimal conditions for uniform editing

Having obtained mutations along 200bp of DNA with arrays of gRNAs in cell culture, we then sought to apply AID-nCas9 to mutate an enhancer in vivo. As a case study, we chose the enhancer regulating stripe 2 of the seven *even-skipped* (*eve*) stripes during early *Drosophila* embryogenesis. Specifically, we targeted a 484 bp long stretch of DNA that has been identified as the minimal *eve2* enhancer, which we will refer to hereafter as *eve2* (Figure 6A). The location of the binding sites for the several activators and repressors regulating this stripe have been characterized (Arnosti et al., 1996; Bishop et al., 2023; Ludwig et al., 1998; Small et al., 1992; Vincent et al., 2018, 2016)(Figure 6B, C). As a result, *eve2* serves as a “gold standard” that offers the opportunity to confirm that any binding sites discovered by randomly mutagenizing this enhancer are consistent with previous annotations, and to discover any new binding sites that have escaped identification to date.

**Figure 6.**
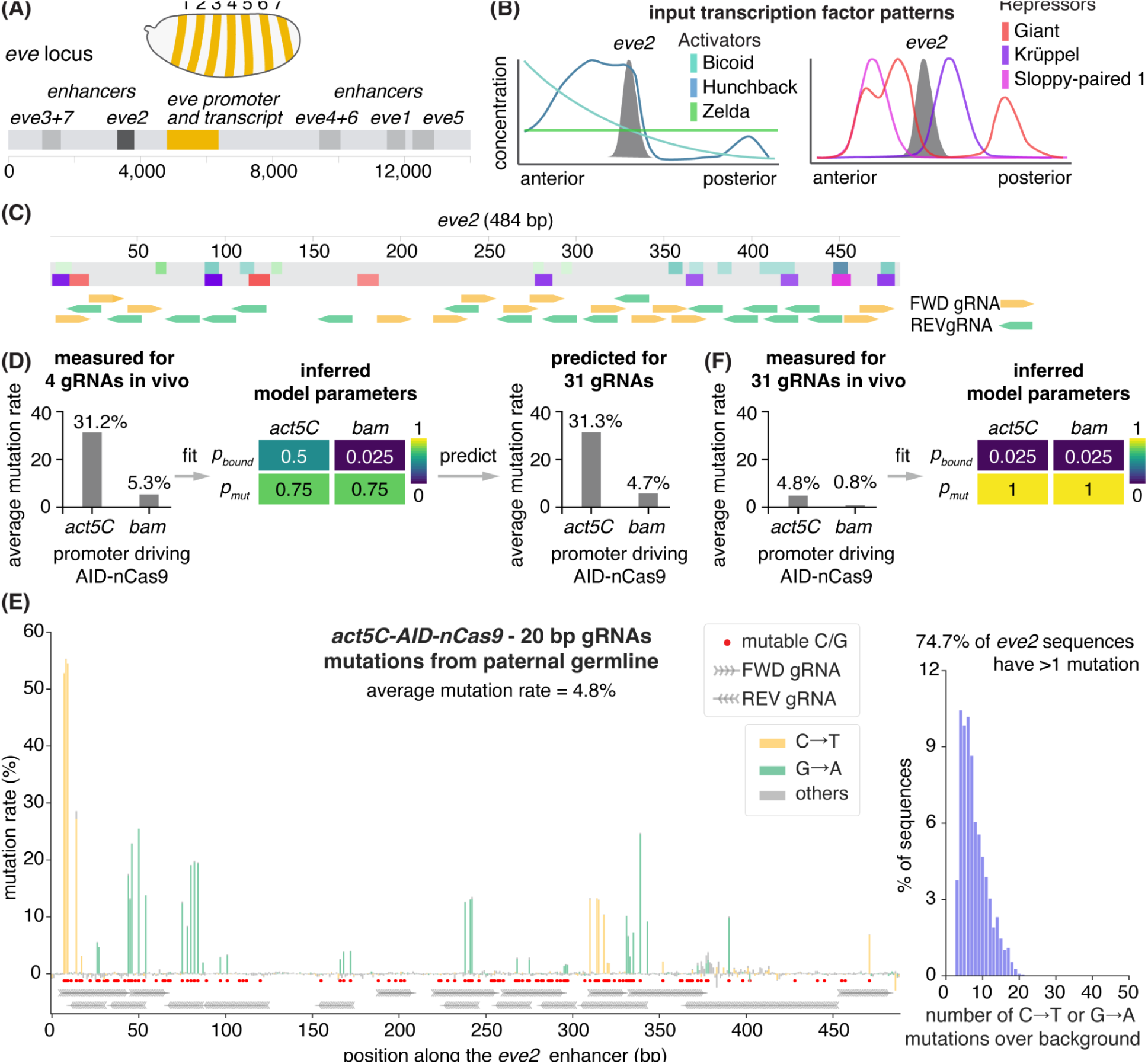
Generating sequence variability in an enhancer in vivo. **(A)** Transcription of the seven *even-skipped* stripes is controlled by different enhancers that respond to unique combinations of transcription factor inputs. **(B)** The enhancer dictating the second stripe of *even-skipped* activity (*eve2*) is controlled by the pioneer factor Zelda and activators produced in the anterior part of the embryo: Bicoid and Hunchback. Repressors Krüppel, Sloppy-paired 1 and Giant delimit the boundaries of the stripe from the anterior and posterior sides. **(C)** Predicted location of activator and repressor binding sites in *eve2* based on functional experiments and bioinformatic inference. The shading of the binding sites is proportional to the site PATSER score, a bioinformatic proxy for binding affinity (Methods - Finding putative binding sites with PATSER). The sequence is overlaid with all gRNAs designed to target the enhancer. **(D)** Measured in vivo mutation rate for the four gRNA experiment were used to infer model parameters for the different promoters controlling AID-nCas9 expression. These parameters were used to infer mutation rates for the 31 gRNAs targeting *eve2*. **(E)** Measured in vivo mutation rate for the 31 gRNAs targeting *eve2*, together with the corresponding parameters inferred using our model. **(F)** Average mutation rate generated in the paternal germline by *act5C-AID-nCas9* and 31 20 bp long gRNAs targeting *eve2* (left) and distribution of observed number of mutations per read after subtracting the number of mismatches observed in control samples (right). Red dots mark all “mutable bases” (Cs on FWD gRNAs and Gs on REV gRNAs).

To target the whole *eve2*enhancer, we identified all potential PAM sequences within it and designed gRNAs adjacent to those sequences. After excluding some overlapping gRNAs that targeted the same Cs this process resulted in 31 gRNAs that covered most of the sequence in the FWD and REV directions (Figure 6C). Importantly, some TA-rich regions in the enhancer remained untargeted due to the lack of NGG PAM sequences (Figure 6C).

We optimized our mutagenesis of *eve2* along two axes. First, based on our cell culture results showing that 23 bp gRNAs expand the mutational window relative to 20 bp gRNAs (Figure 5 - figure supplement 3), we compared the performance of these two lengths in vivo. Once again, because of difficulties in cloning repetitive sequences in the gRNA arrays, subsets of guides were cloned onto multiple plasmids and integrated into distinct landing sites in the fly genome (see Methods – Plasmids).

Consistent with our cell culture results, 23 bp gRNAs extended the mutation window toward the 5’ end. However, these longer guides resulted in lower overall mutation rates than their 20 bp counterparts (Figure 6E; Figure 6—figure supplement 4B,C). We therefore focused on 20 bp gRNAs for subsequent experiments, prioritizing higher overall mutation rates over a broader mutational window.

Second, we examined how *AID-nCas9* expression level, controlled by promoter choice, affects mutagenesis. In vivo, our four gRNAs yielded mutation rates that were too high with *act5C* (31.2%) and too low with *bam* (5.3%) (Figure 3A, B)). In principle, our simulations predicted that these rates should not decrease substantially when scaling from four to 31 gRNAs. However, in the previous section, we showed that larger gRNA arrays in cell culture produced lower-than-expected mutation rates (Figure 5). These results indicate that additional constraints, such as mutagenesis operating in a regime where gRNAs and/or AID–nCas9 become limiting, may emerge when many guides are expressed simultaneously.

To establish a quantitative baseline that captures the effective mutagenesis regime in vivo, we first inferred the mutagenesis parameters from the four gRNA in vivo data to then predict mutation rates for *eve2* targeted by 31 gRNAs. To infer these parameters, we compare the experimentally observed distribution of mutations per DNA molecule with 4 gRNAs, and *act5C* or *bam* promoters, to distributions obtained with our model using a range of *p_bound_*and *p_mut_*. This approach is informative because different combinations of *p_bound_* and *p_mut_* can yield distinct distributions of mutations per DNA molecule despite similar average mutation rates (Figure 6 - figure supplement 1, Figure 6 - figure supplement 2, Methods - Fitting simulated data to experiments). This comparison allows us to uncover the best fit parameters. For the 4 gRNA experiment in vivo with a high mutation rate (*act5C*) these parameters were *p_mut_* = 0.75 and *p_bound_* = 0.5, whereas for the experiment with a low overall mutation rate (*bam*) we obtained *p_mut_* = 0.75 and *p_bound_* = 0.025 (Figure 6D, Figure 6 - figure supplement 3A).

We then used these inferred parameters to predict mutation rates when targeting *eve2* with the 31 gRNAs. Assuming constant levels of AID-nCas9 and each gRNA, the model predicted mutation rates that were too high with *act5C-AID-nCas9* (around 31%), and too low, but closer to our goal of 10%, with *bam-AID-nCas9* (6.4%) (Figure 6D, Figure 6 - figure supplement 3B). We tested these predictions experimentally by comparing mutagenesis of *eve2* under these two promoter conditions.

In vivo, *act5C-AID–nCas9* consistently produced higher mutation rates than *bam*. Using 20 bp gRNAs, *act5C* yielded an average mutation rate of 4.7% across mutable bases, whereas *bam* reached a maximum of 0.8% (Figure 6E, F; Figure 6—figure supplement 4A). Thus, *bam*-driven expression is insufficient to achieve the needed levels of mutagenesis in vivo. However, in both conditions, mutation rates were substantially lower than predicted by our simulations: even for *act5C*, where a mutation rate of 31% was expected, observed rates remained below our target 10% (Figure 6F - figure supplement 3A).

To understand this discrepancy, we fit our model to the experimental data to infer the effective mutagenesis parameters for the 31 gRNA in vivo condition and compared them to those inferred for 4 gRNAs. For *act5C-AID-nCas9* the best fit parameters were *p_mut_* = 1 and *p_bound_*= 0.025 (Figure 6F, Figure 6 - figure supplement 3C). Notably, *p_mut_* remained high (0.75-1) across all conditions. However, *p_bound_* for 31 gRNAs were much lower than the *p_bound_*≈(0.5) observed for the *act5C* condition with 4 gRNAs (Figure 6E). These results indicate that our experiments operate in a regime where the availability of both gRNAs and/or AID–nCas9 is limiting, leading to reduced DNA binding. In contrast, once bound, AID–nCas9 mutates DNA with similar efficiency across conditions. These results indicate that binding, rather than catalytic activity, is the primary limiting step in mutagenesis under these conditions.

We hypothesize that the reduced *p_bound_* reflects limiting gRNA availability when many guides are expressed from arrays. In support of this idea, expressing the same gRNAs at high copy number in cell culture—where each cell contains thousands of plasmids instead of a single genomic integration—resulted in more uniform mutation rates (Figure 6 - figure supplement 4D, Figure 6 - figure supplement 5A).

Our model predicts similar mutation rates across gRNAs and thus relatively uniform mutagenesis along the targeted sequence (Figure 6–- figure supplement 3D). However, in vivo we observed substantial variability in mutation rates across the different positions, with some gRNAs exhibiting little to no mutagenesis (Figure 6 - figure supplement 5). We hypothesized that this loss of uniformity arises from the same limiting regime identified above: differences in gRNA binding efficiency or expression levels—such as positional effects along the transcript in the array where guides located further toward the 3′ end might be expressed at lower levels—lead to unequal effective concentrations of individual guides and thus variable mutation rates.

To test whether differences in gRNA binding efficiency and/or positional effects within the gRNA array underlie this lack of mutational uniformity, we correlated the mutation rate of each gRNA with its ‘binding score’, a proxy for binding energy (Doench et al., 2014; Labuhn et al., 2018; Moreno-Mateos et al., 2015). We found no correlation between mutation rates and gRNA binding scores across multiple scoring metrics or position in the array (Figure 6 - figure supplement 6, Methods - Correlation between mutation rates, gRNA score and position).Our results suggest that factors other than gRNA binding and position within the array are responsible for the observed variability in mutation rates across gRNAs. While the precise source of this variability remains unclear, these findings indicate that achieving fully uniform mutagenesis across long sequences will require additional optimization beyond the current design.

Despite this variability, our approach is sufficient to generate informative levels of mutagenesis in vivo. Specifically, when considering only the mutable bases with a detectable mutation rate (above 0.5%) we obtain average mutation rates of 11.5% and 7.1% for *act5C-AID-nCas9* with 20 bp and 23 bp gRNAs, respectively. Moreover ∼75% of embryos generated by *act5C-AID-nCas9* with 20 bp gRNAs have at least one mutation, indicating that the majority of sequenced embryos contribute information, and thus minimizing sequencing effort spent on unedited samples in high-throughput experiments (Figure 6F, right). Thus, while there is certainly room to improve the coverage and uniformity (see Discussion), the mutation rates achieved at the sites that are frequently mutated are sufficient to meet the requirements of MPRA-like experiments. We therefore proceeded to test whether the mutations generated by this system produce measurable functional effects.

### Random mutagenesis in *eve2* Flybraries reveals changes in expression patterns

We next asked whether random mutations in the *eve2* enhancer generate measurable changes in gene expression that can be used to link sequence to function. In particular, identifying transcription factor binding sites requires measuring both the sequence of each variant and its corresponding levels and spatial pattern of activity.

Because mutations generated in the germline would be inherited alongside a wild-type allele, measuring the effect of individual variants at the endogenous locus would require disentangling heterozygous contributions. To avoid this issue, we generated mutations in the *eve2* enhancer contained within a BAC (bacterial artificial chromosome) reporter construct that encompasses the whole *eve* locus. In this BAC, the *eve coding sequence* has been replaced by 24 MS2 stem loops followed by the *yellow* reporter, enabling quantitative measurement of transcriptional activity (Berrocal et al., 2020).

As an initial step, we mutagenized the *eve2* enhancer within the BAC construct by targeting *act5C-AID-nCas9 to the* enhancer with 20 or 23 bp gRNAs. We then removed the mutagenesis system and generated 22 stable stocks, each carrying a unique mutated version of *eve2* in the BAC. To quantify the effect of those mutations on gene expression, we imaged the seven *eve* stripes using single molecule fluorescent *in situ* hybridization (smFISH, (Calvo et al., 2021)). Specifically, we used hybridizing probes against *yellow* intronic mRNA to detect nascent transcription from the BAC reporter as bright puncta at sites of transcription.

Quantification of embryos carrying the wild-type, unmutated BAC construct revealed a pattern of expression consistent with previous works (Figure 7 - figure supplement 1, (Berrocal et al., 2020), providing a baseline for comparison. Expression profiles were extracted along the anterior–posterior axis and normalized to enable comparison across embryos (see Methods - Image analysis and gene expression quantification). Because *eve* expression is highly dynamic (Berrocal et al., 2020), we divided embryos into four timepoints during nuclear cycle 14, and compared the expression profiles to those of control embryos within the same time class (see Methods - Image analysis and gene expression quantification).

**Figure 7.**
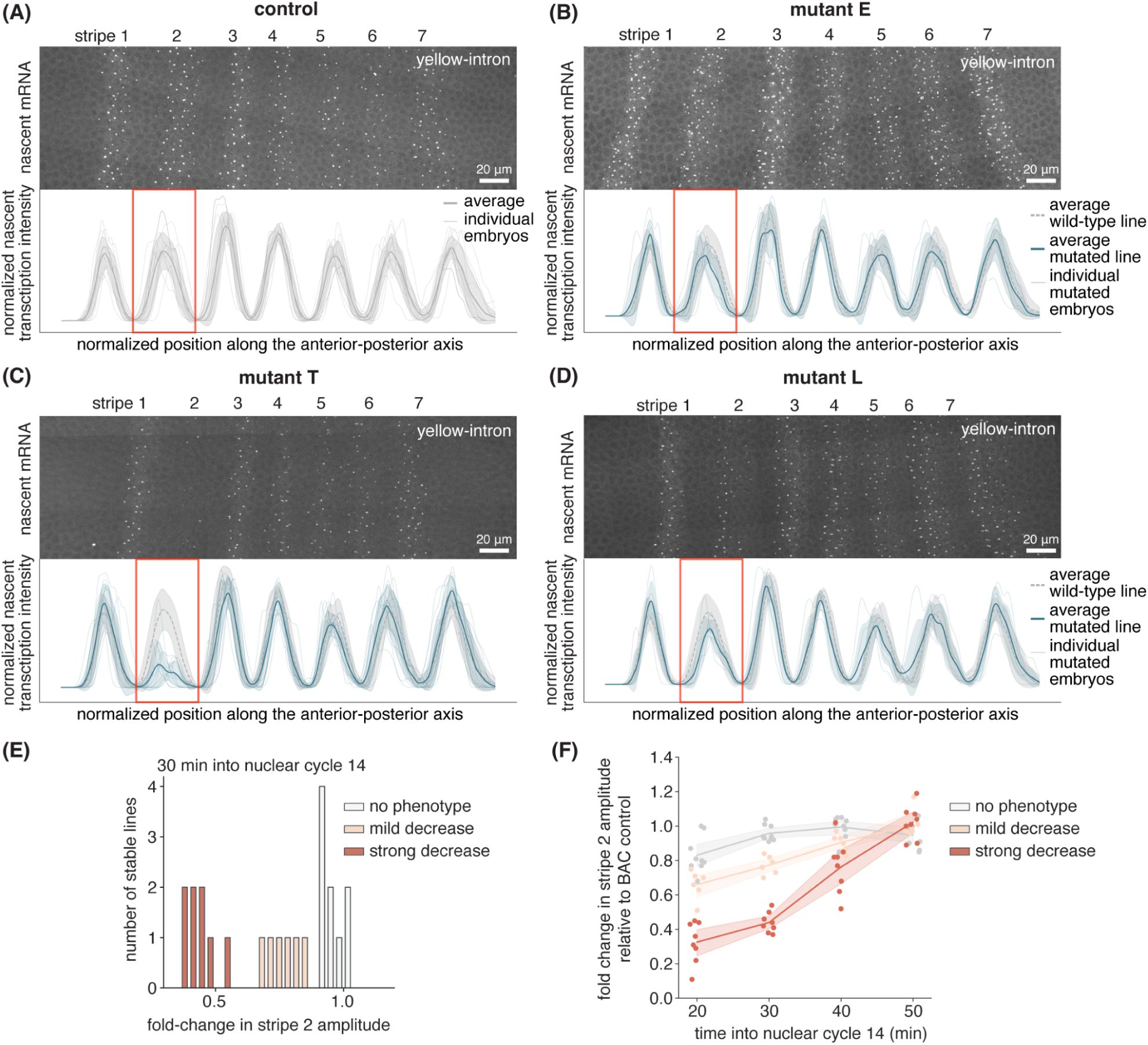
Mutations in the *eve2* enhancer generated by base editing lead to phenotypic changes. (A-D) Expression of *eve-MS2-yellow* measured by smFISH and quantified nascent transcription profiles (smoothed smFISH intensity of the nascent transcription spot summed at each position along the embryo after straightening the stripes and normalizing for the average of all stripe peaks except stripe 2, Methods - Image analysis and gene expression quantification) in control embryos (A), lines with no phenotypic change (B) and lines with mutations that lead to phenotypic changes (C, D). Thick blue lines and shaded areas indicate mean and SD of expression profiles from the indicated line at 30 min into nuclear cycle 14, respectively. Thick grey lines and shaded areas indicate mean and SD of expression profiles at the same timepoint in control embryos with wild-type *eve2* sequences, respectively. Thin blue or grey lines indicate the expression profile from an individual embryo. **(E)** Distribution of fold-change values in stripe 2 amplitude (peak of the second stripe in the nascent transcription profile) compared to controls at 30 min into nuclear cycle 14, used to classify lines into 3 phenotypic categories. **(F)** Changes in stripe 2 amplitude over time for the three categories: no phenotype, mild decrease and strong decrease in expression. Dots indicate the average fold change for all each mutated line. Lines and shaded areas indicate mean and 95% confidence intervals for each category at each timepoint, respectively.

Strikingly, some lines exhibited little to no stripe 2 activity, particularly in the early timepoints (Figure 7A-D). To quantify these phenotypes, we calculated the fold-change in stripe 2 amplitude—defined as the second peak in nascent transcription signal relative to the average of the other stripes—and compared to controls at the same timepoint. At 30 min into nuclear cycle 14, this metric revealed three phenotypic classes (Figure 7E). First, a set of lines (8/22) displayed fold-changes clustered around 1, corresponding to no detectable phenotype (Figure 7B). Second, a comparable number of BAC lines (8/22) exhibited markedly reduced levels of stripe 2 expression (fold-change < 0.55), which we term ‘strong decrease’ (Figure 7C, Figure 7 - figure supplement 2). Finally, the remaining lines (6/22) displayed a smaller but consistent reduction, which we called ‘mild decrease’ (fold-change 0.7–0.84, Figure 7D, Figure 7 - figure supplement 3).

Our quantification revealed a striking temporal phenotype: differences between lines were more obvious at earlier timepoints (20 and 30 min into nuclear cycle 14), with expression reduced as much as 80% in some lines (Figure 7E-F, Figure 7 - figure supplement 4, 5). However, at 40 and 50 min, nascent transcription returned to levels equivalent to controls. These phenotypes were highly reproducible, with nearly all embryos carrying the same mutation exhibiting similar reduced levels of transcription at a given time point. Together, these results indicate that systematic dissection of enhancer function in vivo requires quantification not only of spatial patterns of gene expression, but also of their temporal dynamics.

To further dissect the quantitative changes in transcriptional activity that led to reduced stripe 2 expression in our mutated embryos, we measured average spot intensity, stripe width and fraction of active nuclei in stripe 2 at the 30 min timepoint (see Methods - Image analysis and gene expression quantification). We found that reduced expression was not due to a decrease in the overall brightness of individual transcription spots, but due to a decrease in the number of transcription spots (Figure 7 - figure supplement 6). This data is therefore consistent with a model where many mutations affect the probability or timing of transcriptional activation rather than the transcriptional output of already active nuclei. Interestingly, live imaging studies of *eve* stripe 2 expression have also revealed that both of these transcriptional features are under regulatory control (Berrocal et al., 2020; Bothma et al., 2014; Lammers et al., 2019). Consistent with the transient nature of the phenotype, all parameters returned to wild type levels at later timepoints (Figure 7 - figure supplement 7).

Together, these results demonstrate that random enhancer mutagenesis in vivo generates reproducible and quantifiable changes in transcriptional dynamics that can be linked to specific enhancer variants. Further, the transient nature of the phenotype we uncovered suggests that systematic dissection of enhancer function in vivo requires quantitative measurements of both the spatial and temporal dynamics of gene expression. Thus, Flybraries can constitute the basis of a platform for mapping sequence–function relationships directly in endogenous developmental contexts.

### Using *eve2* Flybraries to draw insights into regulatory architecture

Having identified mutants that led to quantifiable phenotypes, we next sought to shed light on the changes in transcription factor binding site architecture responsible for these phenotypes. To make this possible, we sequenced the *eve2* enhancer within the BAC reporter construct of each of our 22 fly lines (see Methods - Genotyping stable lines). We then grouped these lines by phenotypic category to look for correlations between phenotype and DNA sequence (Figure 8A). To calculate these correlations, we computed the mutual information between mutations at each position along the enhancer and the corresponding stripe 2 amplitude phenotypic category. This measure has been used in MPRA experiments to find regions likely bound by repressors or activators (Ireland et al., 2020). Despite our limited mutational coverage and statistics, this approach was useful in identifying the C-to-T mutation at position 13 (C13T) as the most informative position (Figure 8B). Indeed, 8 out of 9 fly lines in which this mutation occurs correspond to the ‘strong decrease’ category.

**Figure 8.**
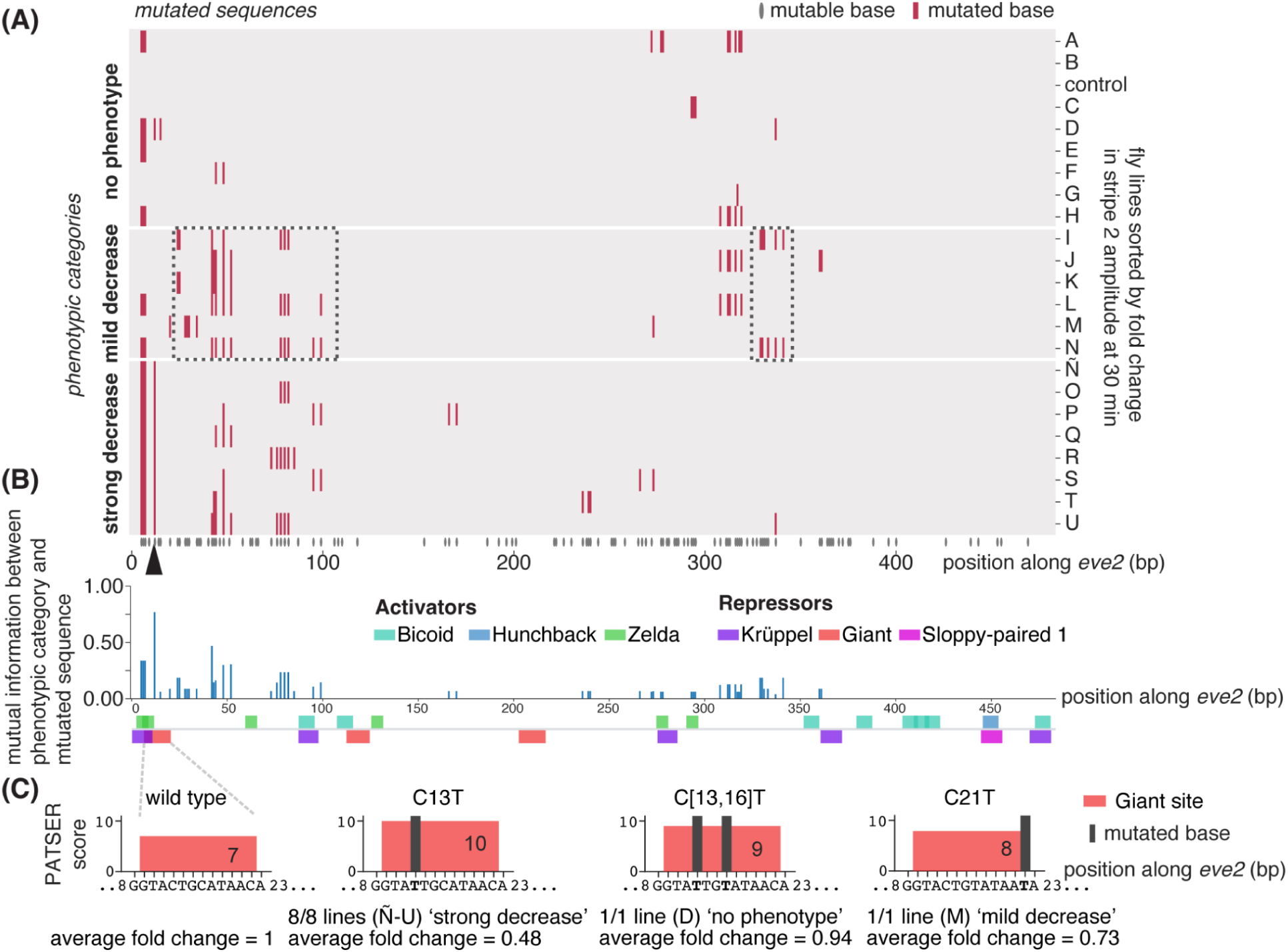
Identifying DNA positions responsible for the phenotypic changes. **(A)** Quantification of changes in gene expression over the 22 stable fly lines analyzed. The position of mutations for each sequence is shown as red lines. Lines are sorted by the fold change in stripe amplitude at 30 min into nuclear cycle 14. 8/22 lines show a ‘strong decrease’ in activity and 6/22 a ‘mild decrease’. The mutation on position 13 (C13T) that highly correlates with ‘strong decrease’ lines is marked with an arrowhead. All mutable bases are shown by grey lines below the plot. **(B)** Mutual information between sequence and stripe 2 amplitude phenotypic category calculated at each position. Transcription factor binding sites predicted by PATSER are shown below. The fourth C→T mutation (C13T) is the position that contains the most information, consistent with (A). **(C)** Changes in binding sites predicted by PATSER in the different mutations obtained at the 5’ end of the *eve2* enhancer impacting a Giant site binding site. Mutations are indicated by the vertical blue lines. Predicted Giant sites are shown as red boxes, with their height indicating the PATSER score for the wild-type binding site and for the three kinds of mutations in this region. Mutations are shown in boldface. The C→T mutation at position 13 increases the score of a Giant site. Other mutations lead to smaller increases on the Giant site score.

Using these sequences, we then ran the PATSER algorithm (see Methods - Finding putative binding sites with PATSER) on all mutated sequences to predict changes to the binding sites for all known regulators of *eve* stripe 2 (Figure 8 - figure supplement 1). All the lines with the lowest expression (8/8 of the ‘strong decrease’ category) contained the C13T mutation identified above. Our bioinformatic analysis predicts that this mutation should increase the binding affinity of this site for the Giant repressor (Figure 8A, arrowhead; Figure 8B; Figure 8 - figure supplement 1). Interestingly, other mutations that increased the PATSER score of this Giant site, but to a lesser extent, fell into the ‘no phenotype’ or ‘mild decrease’ categories (Figure 8C, lines D and M).

We note all the ‘strong decrease’ lines also contain three additional C-to-T mutations 5’ of C13T mutation. These mutations (C[6,7,8]T) are predicted to disrupt sites for the Krüppel repressor and pioneer factor Zelda (Figure 8 - figure supplement 1 B). However, these mutations are also present in other lines, such as line E, that were classified as ‘no phenotype’ (Figure 7B; Figure 8A).

Thus, our observations are compatible with the reduced stripe 2 amplitude in our mutants originating from enhanced repression of *eve2* due to an increase in affinity of the Giant site. Further, these results also indicate that stripe 2 expression is largely insensitive to mutations of the predicted Krüppel and Zelda sites at this location. This insensitivity suggests that these bioinformatically identified sites, which have not been individually tested in the context of functional assays, are not functional (Arnosti et al., 1996; Bishop et al., 2023; Small et al., 1992).

None of the 6 lines classified as ‘mild decrease’ contained the C13T Giant mutation that seems to be responsible for stripe amplitude reduction in the ‘strong decrease’ lines. Instead, these fly lines presented mutations in regions devoid of any predicted binding sites (Figure 8A, boxes; Figure 8 - figure supplement 1). Our bioinformatic analysis predicts that some of these mutations lead to the creation of putative Krüppel, Sloppy-paired 1 or Hunchback sites, or to the modification of existing Giant, Krüppel or Bicoid sites throughout the sequence (Figure 8 - figure supplement 1). However, no clear patterns between these mutations and phenotypes were observed.

Overall, our results show that it is possible to generate mutations within an enhancer by high-throughput in vivo mutagenesis, to sequence those mutations, and to measure their functional consequences on gene expression. Most importantly, all of these measurements can be brought together to perform a systematic dissection of transcription factor binding sites within an enhancer, and of the regulatory logic by which those sites dictate patterns of gene expression in space and time.

## Discussion

The advent of MPRAs has revolutionized our ability to query biological phenomena, from dissecting gene regulatory regions to systematically studying protein function (Arnold et al., 2013; Inoue and Ahituv, 2015; Ireland et al., 2020; Kheradpour et al., 2013; Kinney et al., 2010; Staller et al., 2022). Yet, these advances have been mostly relegated to cells in culture, where the transfection of massive libraries of DNA is feasible. Indeed, with some notable exceptions (Brown et al., 2022; Chan et al., 2023; Farley et al., 2015; Jores et al., 2020; Lagunas et al., 2023; Stevenson et al., 2023), multicellular organisms have remained on the sidelines of the great progress heralded by MPRAs. Further, because of their reliance on reporter constructs, it has been challenging to deploy MPRA-like technologies in endogenous genomic *loci* neither in the context of single cells, nor in the context of multicellular organisms.

Whole genome CRISPR approaches such as Perturb-seq have emerged to probe the effect of modulating the expression of endogenous genes by recruiting activating or repressing to their regulatory regions (Alda-Catalinas et al., 2020; Dixit et al., 2016; Zheng et al., 2025). Recently, the generation of DNA variants of endogenous regulatory regions has been made possible using prime editing and tiled based editing arrays. While mostly applied in cell culture (Becerra et al., 2024; Lue and Liau, 2023; Ravi et al., 2022), a recent study has applied tiled based editing arrays in vivo using mouse xenographs (Ren et al., 2026).

In this work, we chose to explore the use of tiled base editing approaches to generate variant libraries in multicellular organisms. We made significant progress towards realizing this goal by introducing technology for targeted high-throughput in vivo genomic mutagenesis in endogenous *loci* in the fruit fly. Specifically, we repurposed base editing tools (AID fused to nCas9) to generate random mutations within a 200-500 bp sequence by using >20 gRNAs expressed in an array.

We found that lamprey AID (evoCDA1, (Thuronyi et al., 2019)) was highly active in *Drosophila* in vivo and in cell culture, whereas other mammalian AID/APOBEC variants (Kohli et al., 2021), heavily used in mammalian cell culture, were active in *Drosophila* cell culture but not in animals. Interestingly, because of their use in precision genome engineering (Koblan et al., 2021; Musunuru et al., 2021; Newby et al., 2021; Villiger et al., 2018), most AIDs have been selected to be highly efficient and accurate at introducing mutations within a narrow editing window (Anzalone et al., 2020; Thuronyi et al., 2019). However, for our purposes, enzymes with low mutation rates (∼10%) and with a wide editing window are needed. Thus, our discovery of lamprey AID being more appropriate for our purposes in animals emphasizes the need to scout different sources of mutagenesis enzymes, and suggests that optimization of these enzymes for our specific purposes might be necessary. Further, this realization is also a stark reminder that what is learned in a cell culture context does not always translate directly to the context of the full, intact organism.

### Current limitations of our approach and further optimization

The ability to tune mutation rate is critical to extracting the most information out of our variant libraries. For example, computational studies suggest that mutations generated randomly along a DNA sequence with an 10% mutation rate are optimal for finding transcription factor binding sites and transcription factor binding energy matrices in MPRAs in bacteria, and that this combinatoriality is needed to uncover cooperative sites (Ireland et al., 2020; Pan et al., 2024). Yet, while in vitro-generated libraries for MPRAs can be designed with a prescribed mutation rate, this is not the case in our approach to in vivo genomic mutagenesis. Fortunately, using different promoters to tune the timing and level of expression of our AID-nCas9 fusion, and informed by a simple mathematical model, we have demonstrated the ability to tune average mutation rates between 5 and 25% at many sites.

However, in addition to having the right overall mutation rate, uniform coverage is also essential to extract information from variant libraries. Indeed, while our mutation rates were in line with our goals in many sites, we found that small regions of the *eve2* enhancer remained untargeted due to the NGG PAM sequence constraints of nCas9 (Figure 6C). Here, Cas9 variants, such as Cas12 (Chen et al., 2022) or the nearly PAMless SpRY-nCas9 (Walton et al., 2020) could be used to break free this NGG constraint. Alternatively, other targeting systems, such as those that rely on fusions to T7 polymerase (Álvarez et al., 2020; Cravens et al., 2021; Moore et al., 2018; Park and Kim, 2021) or helicase-assisted (Chen et al., 2024) could also be deployed.

Another significant difference between our approach and the generation of libraries in vitro is related to the mutational spectrum. In vitro DNA libraries can be designed to contain all possible base substitutions. However, our AID-nCas9 approach can only lead to C→T or G→A mutations. While it remains to be experimentally determined whether this limited mutational spectrum can provide the same information as in vitro generated libraries, one recent computational study in bacteria suggests that mutations produced by C and A base editors, but not just C editors, can indeed reveal where binding sites are in bacterial promoters (Pan et al., 2024). While our experiments only used C base editors, we envision that in future work other base editors could be combined with AID to increase the mutational spectrum. For example, the adenine deaminase ABE8e (Fu et al., 2021) could be used to incorporate A→G and T→C mutations, allowing the substitution of all four base pairs, and newly engineered glycosylases could generate G→C and G→T mutations (Tong et al., 2023).

### Flybraries shows the potential for high-throughput in vivo regulatory dissection in animals

Our approach offers the possibility of breaking free from the low-throughput that has characterized the study of DNA regulatory regions in the context of animals. For example, over the last forty years, and with only one exception that required herculean effort (Fuqua et al., 2020), no work in the fruit fly has produced more than 50 DNA regulatory variants that are incorporated into stable, transgenic lines (Falo-Sanjuan et al., 2019; Ip et al., 1992, 1992; Sayal et al., 2016; Small et al., 1992; Swanson et al., 2010; Szymanski and Levine, 1995). In contrast, because each fly mother produces hundreds of eggs (Vijayan et al., 2022), we envision that our system could be easily scaled up to generate tens of thousands of embryos, each with a unique set of mutations at a DNA locus of interest, in a single experiment.

Importantly, the ultimate aim of this technology is to enable the quantification of phenotypic changes in individual embryos, each with its own mutated sequence. However, as a first step toward establishing the feasibility of this approach, we generated 22 stable fly lines, each carrying a unique set of mutations. These flies made it possible to characterize the robustness of the resulting phenotypes and the sensitivity of our fluorescence *in situ* hybridization quantification approach. Using these stable lines, we showed that mutations lead to quantifiable changes in gene expression. Specifically, our experiments revealed that, while these changes displayed a dynamical phenotype, once that timing was considered, even subtle phenotypic changes in gene expression patterns became detectable and quantifiable. As a result of the precision and sensitivity demonstrated in measuring the gene expression patterns of embryos of stable lines, we envision that the quantification of phenotypes in single embryos, each with its own mutation, should be possible.

It is interesting that our mutations of the *eve2* enhancer only impact transcription over a 10-20 min developmental window. It is possible that this mutation delays chromatin accessibility or transcription initiation. Alternatively, it is possible the mutations in *eve2* decrease activity throughout nuclear cycle 14 but we are not able to detect it at later timepoints because endogenous Eve protein is still able to activate the *eve autoregulatory element* that is also contained in the BAC. Recent observations also suggest that *eve* expression recovers at the end of nuclear cycle 14, even in the absence of the *eve2* enhancer (Fischer et al., 2025). As a result, it is also possible that at late timepoints we are detecting expression regulated by other unknown DNA regions.

Finally, beyond showing that we can generate unique sequence variants and that these variants can lead to quantifiable gene expression phenotypes, we also demonstrated that correlations between sequence and phenotype can lead to molecular insight. Specifically, bioinformatic analyses suggest that the mutations that lead to a strong decrease in stripe 2 expression result from the increase in affinity of a Giant repressor binding site. Additionally, we have identified areas with no predicted binding sites that lead to small decreases in expression when mutated. This observation is consistent with the work from (Fuqua et al., 2020) revealing that almost all positions in an enhancer carry some information, and is a stark reminder of the importance of performing unbiased mutational scans of enhancer sequences.

### Future applications of MPRAs in animals and in endogenous loci

We anticipate that our technology will make it possible to extract the same information from hundreds of individual embryos harboring slightly different combinations of mutations as we obtained from the 22 stable lines. Specifically, using the microscopy-based platform featured in this work, we aim to be able to quantify the gene expression patterns of ∼1,000 embryos in this mid-throughput manner. To be able to test even more variants and rapidly scan an enhancer to uncover binding sites, sequencing approaches will be needed.

As a first step to increase throughput in the number of variants tested via sequencing, our technology could be used to dissect the regulatory content of enhancers that are contained within a STARRseq (self-transcribing active regulatory region sequencing) construct. In this reporter-based approach, enhancers are placed downstream of promoters, within the intron of a reporter gene. Thus, sequencing of the resulting transcript not only reports on gene expression levels, but also on the mutated enhancer sequence driving those gene expression levels (Arnold et al., 2013).

Further, to go beyond reporter constructs and mutagenize enhancers in their endogenous context we need to simultaneously measure mutated DNA enhancer sequence and the transcriptional activity of the gene regulated by the enhancer. We envision that future developments in single cell DNA and single-cell RNAseq will make this possible (Olsen et al., 2023; Yu et al., 2023). It is important to note that this ability to mutagenize endogenous regulatory regions would even be useful in contexts where reporter-based MPRAs have been traditionally feasible, such as bacteria, yeast or mammalian cell culture. Indeed, our approach , as well as other recent in vitro and in vivo tiled based editing approaches (Becerra et al., 2024; Lue and Liau, 2023; Ravi et al., 2022; Ren et al., 2026; Roh et al., 2025), could be used to mutagenize and test endogenous genomic loci without losing local genomic information such as interactions with nearby sequences or chromatin state that cannot be captured using reporter constructs.

Importantly, our technology is not limited to targeting regulatory sequences. For example, over the last few years MPRAs have been used to shed insights on the function of protein domains such as transactivation domains (DelRosso et al., 2023; Staller et al., 2022; Thurm et al., 2026; Valbuena et al., 2024). We envision that our in vivo targeted mutagenesis approach could also be used to systematically mutagenize such small protein domains in their endogenous context while simultaneously measuring the resulting phenotypic outcome.

Overall, we have developed a versatile tool that can be used to mutagenize regulatory sequences, or virtually any other sequence, in *Drosophila*. We posit that this tool can be readily adapted to other multicellular and single-celled organisms. Building on recent previous works using MPRAs to find the number, placement and affinity of transcription factor binding sites in previously uncharted regulatory regions (de Almeida et al., 2022; Ireland et al., 2020), future work from our lab will harness this new technology to map the regulatory architecture of enhancers relevant to fly embryogenesis in a single experiment.

## Supplementary figures

**Figure 1 figure supplement 1.**
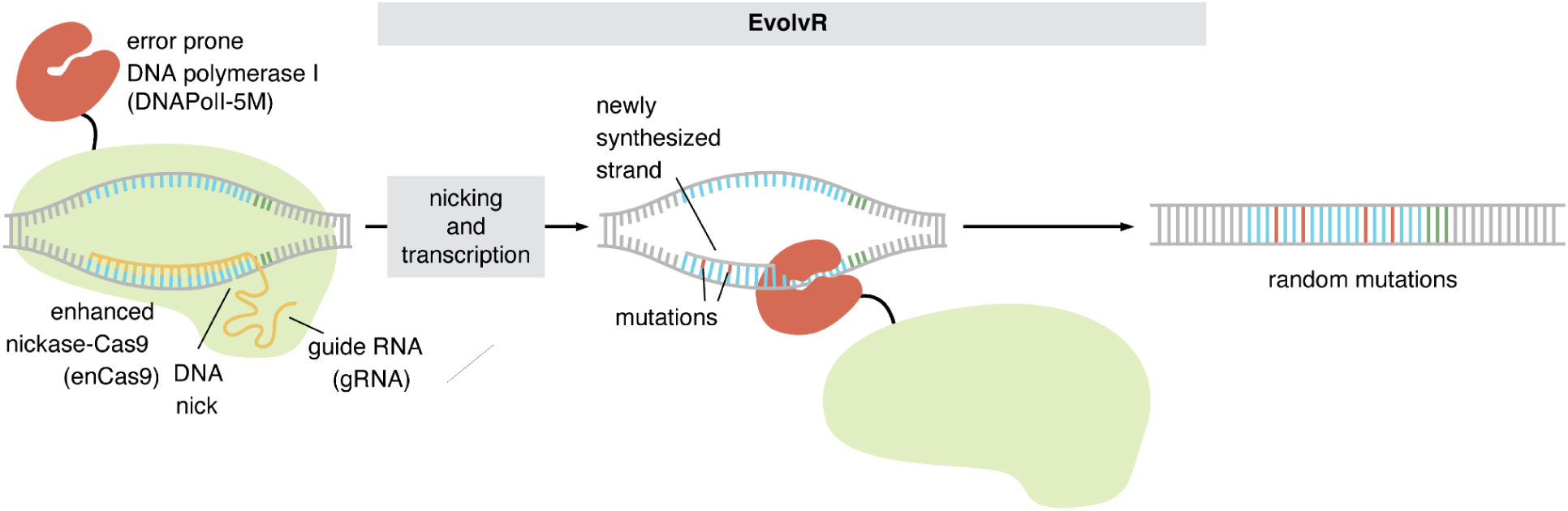
The EvolvR system. In the EvolvR system, enhanced-nickase Cas9 (enCas9) targeted to a locus makes the DNA accessible for an error prone DNA polymerase I from *E. coli* to transcribe the DNA. The DNA nick introduced by enCas9 is also required for mutagenesis.

**Figure 2 figure supplement 1.**
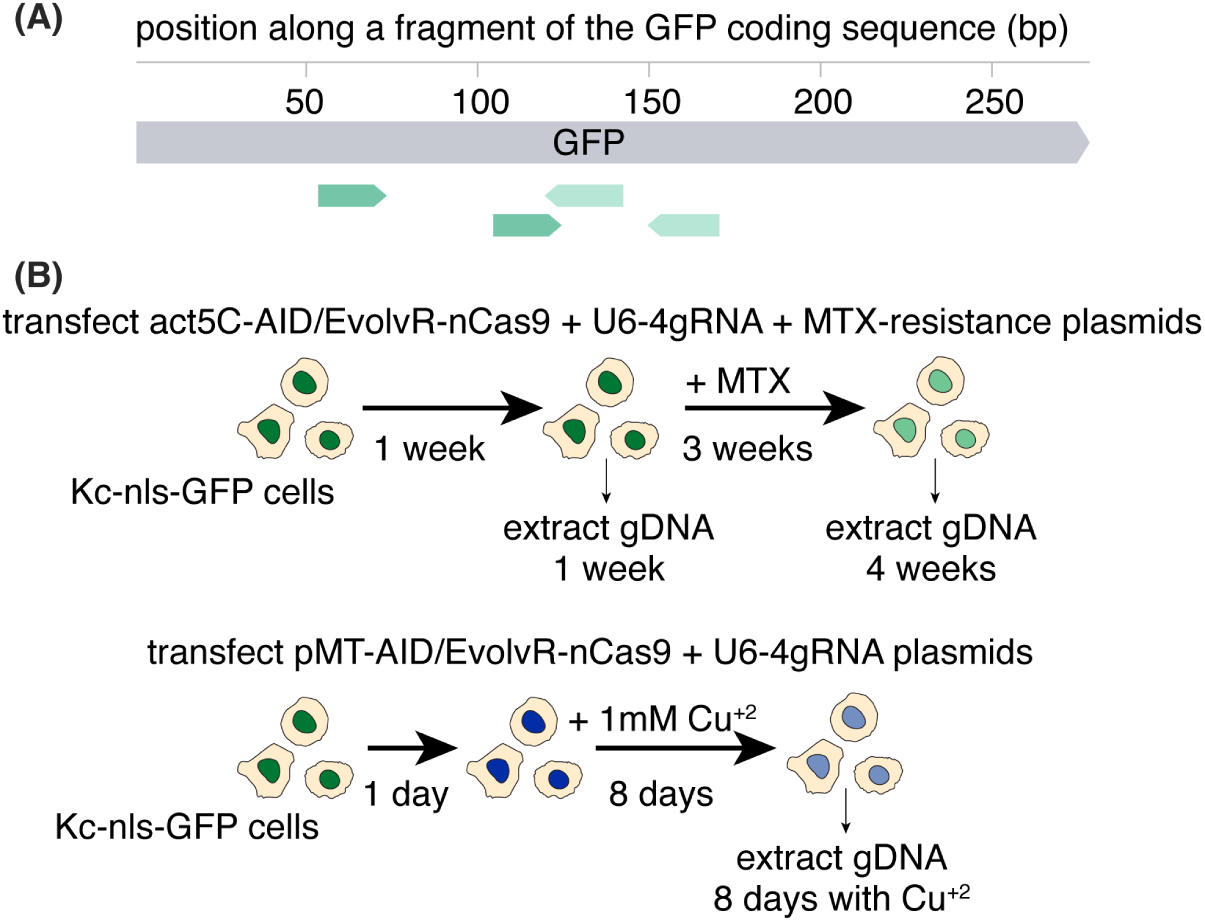
Experimental scheme for cell culture experiments. **(A)** Binding location of the 4 gRNAs targeting the chromosomally integrated GFP gene. **(B)** Experimental protocol to induce and assay mutagenesis in cell culture. Timepoints of genomic DNA collection of Kc-NLS-GFP cells after transfection of gRNAs, AID-nCas9 and MTX resistance plasmids. Genomic DNA was collected after 1 and 4 weeks for *act5C* driven AID-nCas9 or EvolvR-enCas9 (top), and after 9 days in the case of *pMT* (bottom). GFP was then amplified and sequenced using Nanopore sequencing.

**Figure 2 figure supplement 2.**
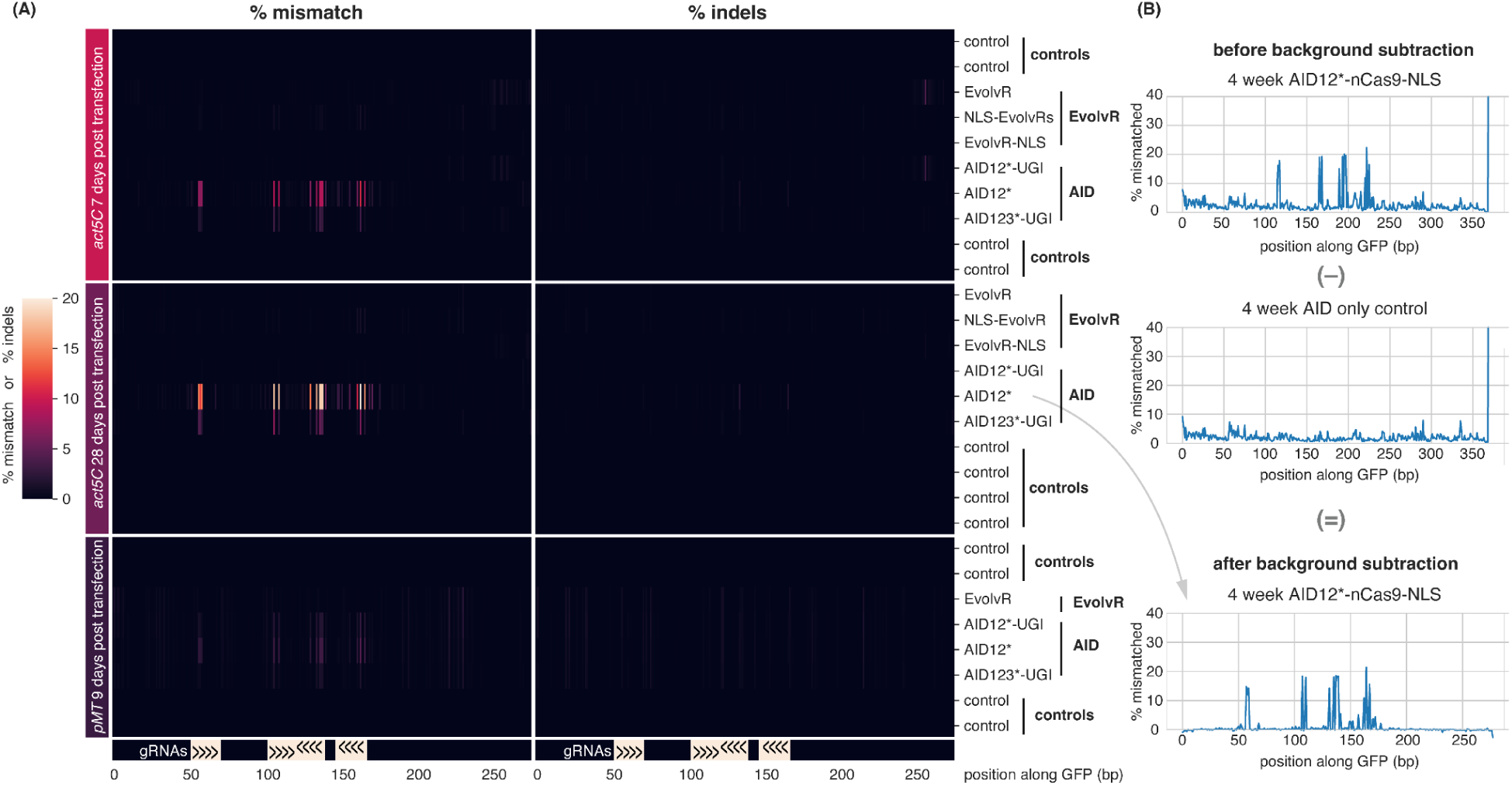
Quantifying mutations by AID in cell culture using Nanopore sequencing. **(A)** Heatmap showing percentage of mismatch (left) and indels (right) to the GFP sequence in the different timepoints and mutagenic enzyme conditions after subtracting background mutations detected in controls (see Methods - Analysis of sequenced Nanopore libraries). AID samples exhibit a high percentage of mismatches aligning with the location of the gRNAs. **(B)** Example mismatch profiles before and after background subtraction.

**Figure 2 figure supplement 3.**
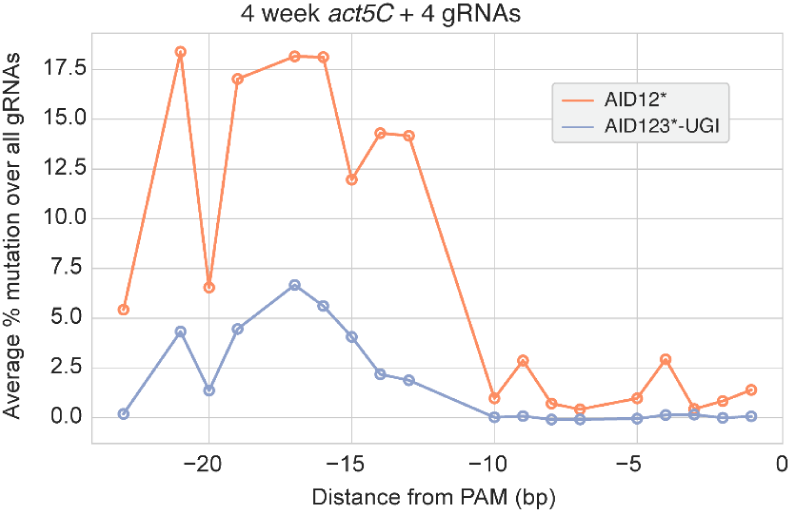
Average mutation profiles along gRNAs. Mutation rates on each position within a gRNA averaged over the four gRNAs targeting GFP. The two AID mutagenic enzymes with the highest rates are shown. Mutations occur primarily in the 5’ half of the gRNAs.

**Figure 2 figure supplement 4.**
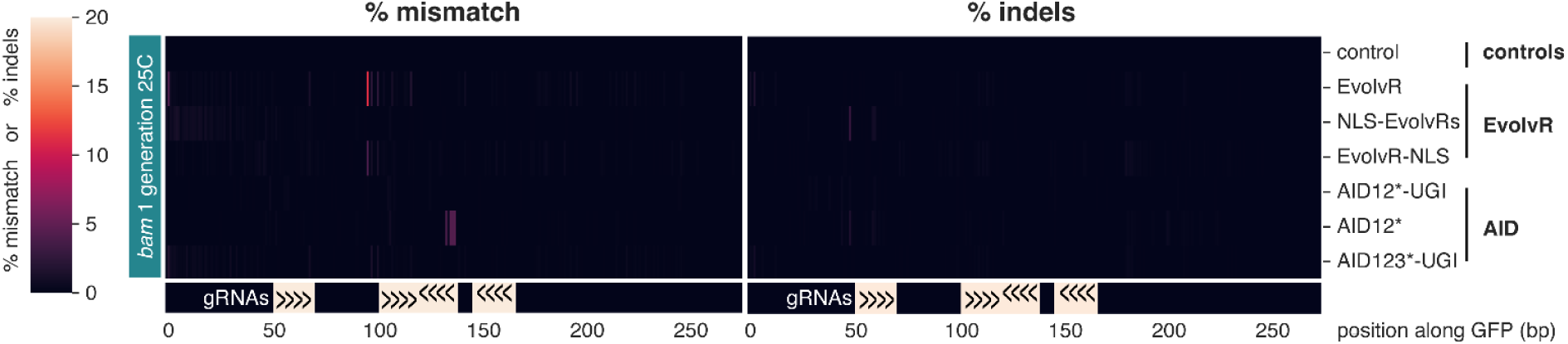
Testing mutagenesis by base editing and EvolvR in vivo. Heatmaps of percentage of mismatch and indels to the GFP sequence after one generation at 25°C. Background mutations detected in controls in embryos are subtracted from each condition using the indicated mutagenic enzymes. Few mismatches were detected in the AID12* sample and only one potential mismatch in the EvolvR sample. Arrows indicate the location of the 4 gRNAs used in this experiment.

**Figure 3 figure supplement 1.**
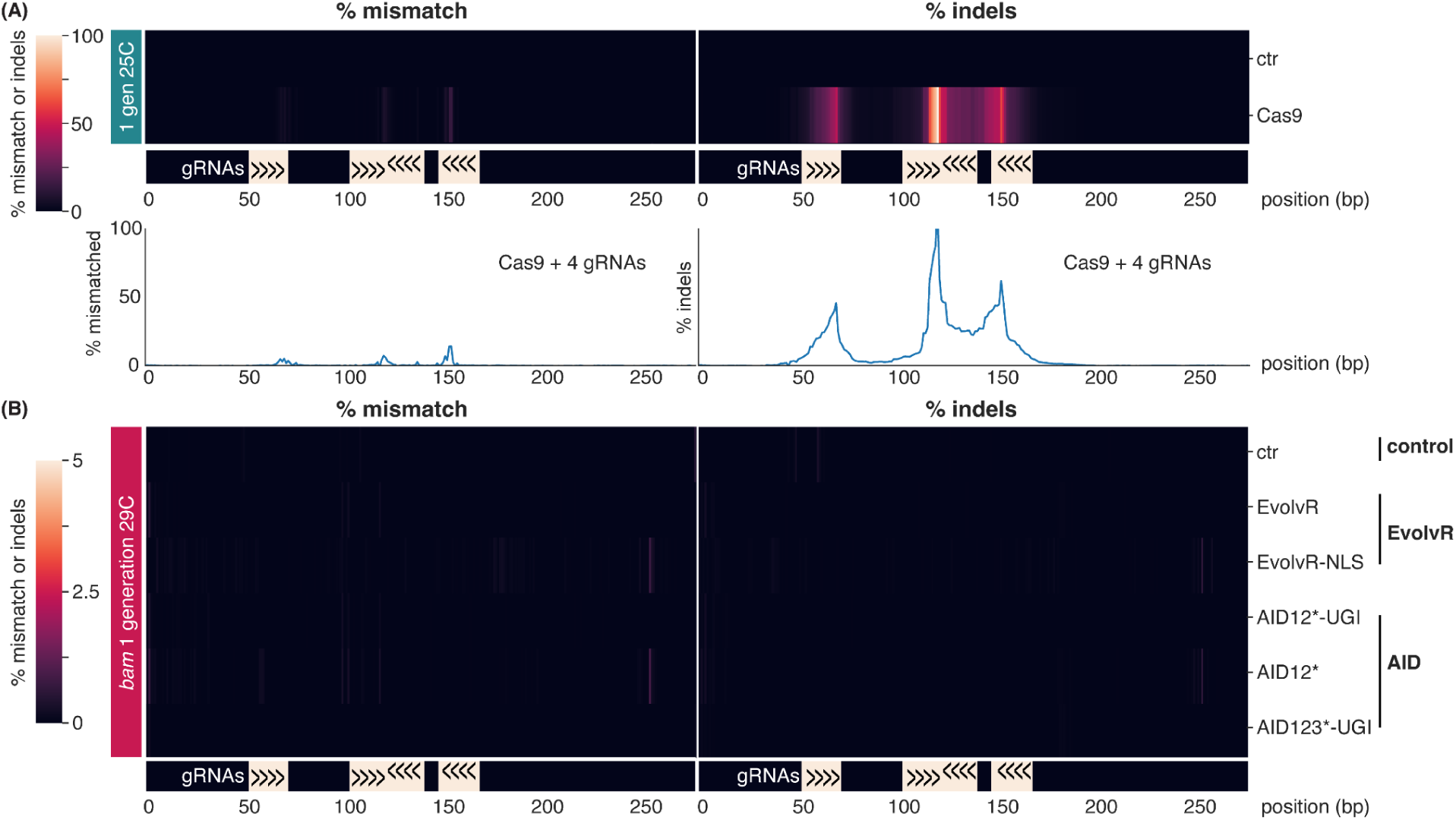
Testing gRNAs in flies using catalytically active Cas9 and higher temperatures. **(A)** Heatmap showing percentage of mismatch and indels to the GFP sequence after subtracting background mutations detected in controls in embryos where GFP was targeted with 4 gRNAs and catalytically active Cas9. Bottom plots show the same profiles. Cas9 introduces indels at target sites with variable efficiencies. **(B)** Heatmap showing percentage of mismatch and indels to the GFP sequence after subtracting background mutations detected in controls in embryos where the indicated enzymes could have generated mutations for one generation at 29°C expressed from the *bam* promoter. No mismatches or indels were detected. Arrows indicate the location of the 4 gRNAs used in this experiment.

**Figure 3 figure supplement 2.**
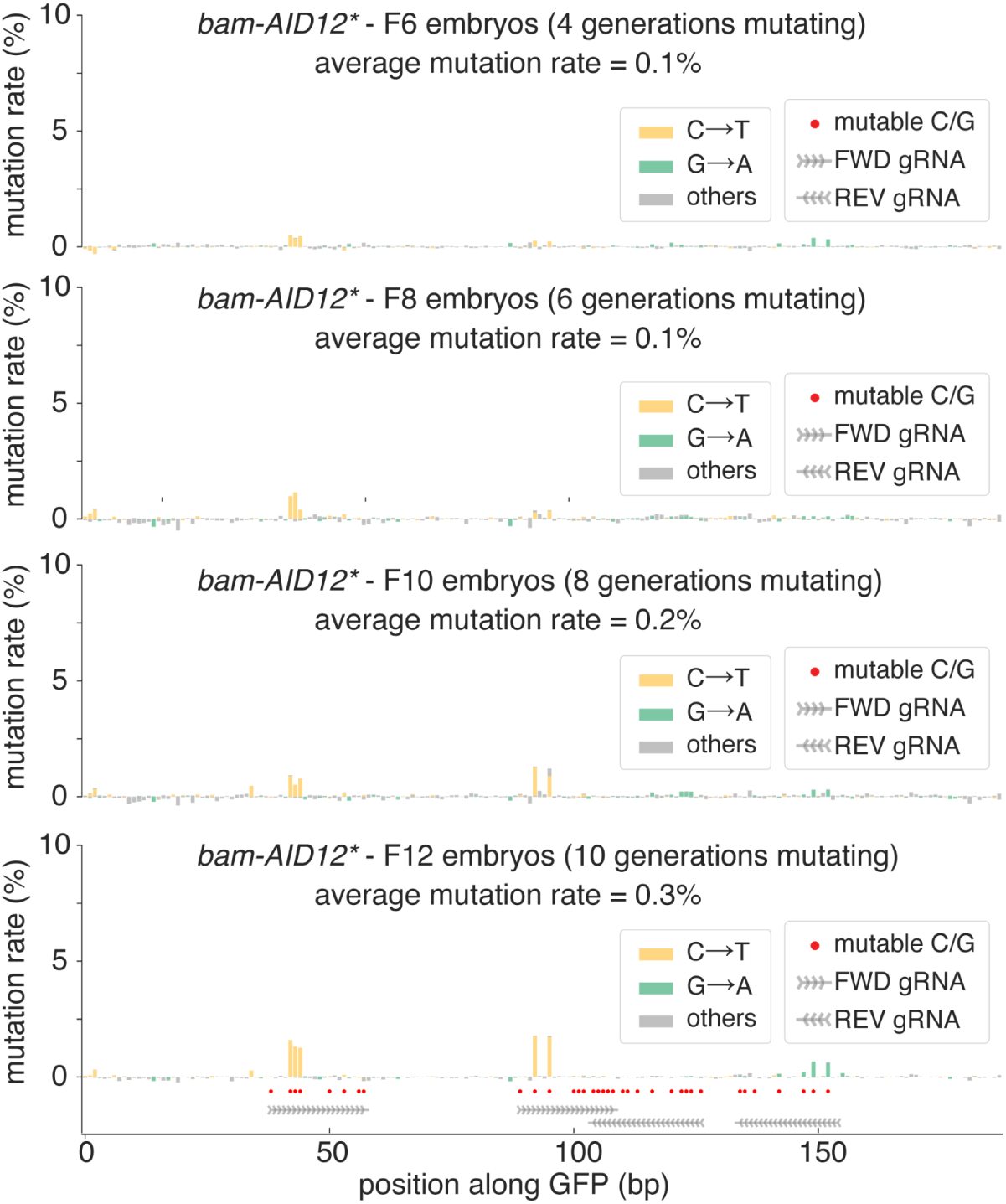
Mutagenesis in flies over multiple generations. Mismatches to GFP classified by base substitution in embryos from stable stocks expressing *bam-AID12\** and 4 gRNAs targeting GFP, after 4, 6, 8 and 10 generations mutating. The average mutation rate is calculated as the mean of detected C→T or G→A mutations over background in all “mutable bases” (Cs on FWD gRNAs and Gs on REV gRNAs, red dots).

**Figure 3 figure supplement 3.**
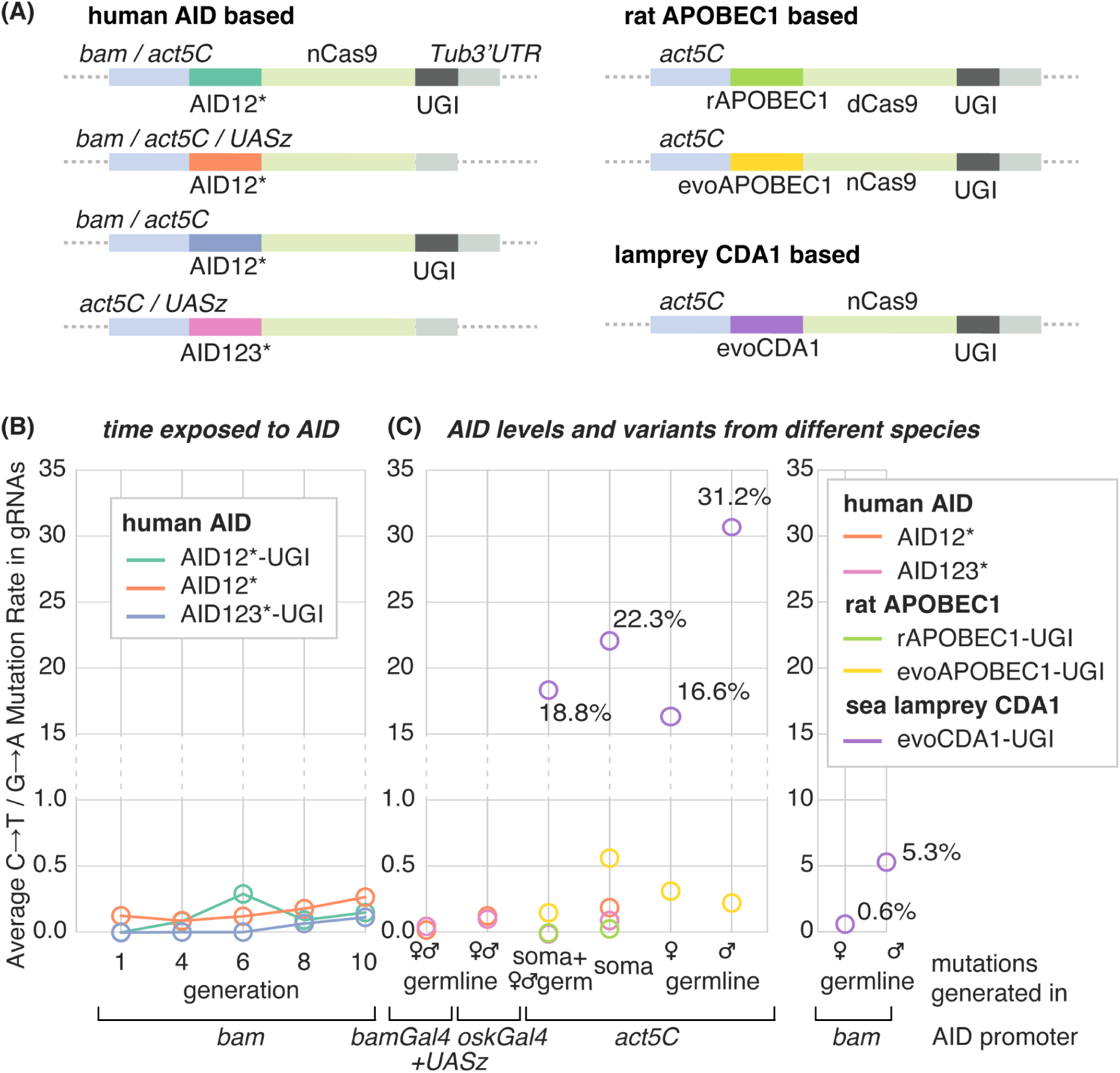
(A) Plasmids with different versions of AID used in experiments in Figure 3 to mutagenize a GFP fragment in the genome of fly embryos using 4 gRNAs. We tested AID stemming from different organisms under a plethora of different promoters. For the inducible UASz constructs, we crossed to *bam-Gal4* and *osk-Gal4* to induce high germline expression. **(B-C)** Average mutation rate (defined as C**→**T or G**→**A mutations over all mutable positions) under different conditions in embryos. Propagating fly lines over multiple generations (A) or replacing the *bam* promoter with *act5C* or *UASz* had no or very little effect on the mutation rate (B). Only *act5C-evoCDA1-nCas9* was successful at producing a high number of mutations in embryos.

**Figure 3 figure supplement 4.**
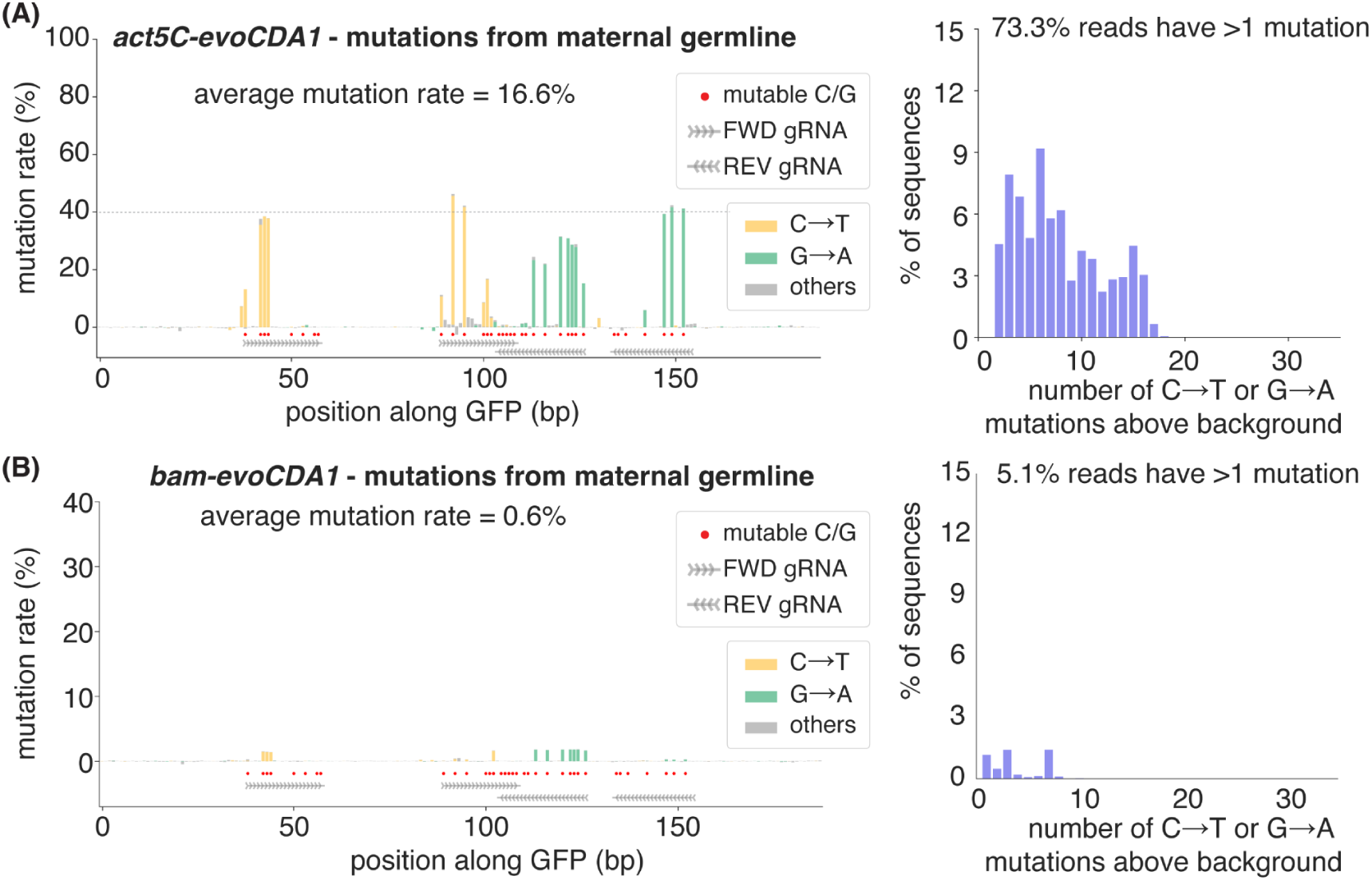
Mutagenesis by lamprey evoCDA1 generated in the maternal germline. Comparison of average mutation rates obtained by evoCDA1 expressed from *act5C* **(A)** or *bam* **(B)** promoters in embryos and adults mutated only in the maternal germline. Mutations are significantly lower with *bam* than *act5C,* and about half as numerous as those obtained in the paternal germline (Figure 3A, B). Left bar plots show mismatches in GFP quantified after background subtraction and classified by base substitution and right histograms show the distribution of the observed number of mutations per read after subtracting the number of mismatches observed in control samples (right). Red dots mark all “mutable bases” (Cs on FWD gRNAs and Gs on REV gRNAs).

**Figure 4 figure supplement 1.**
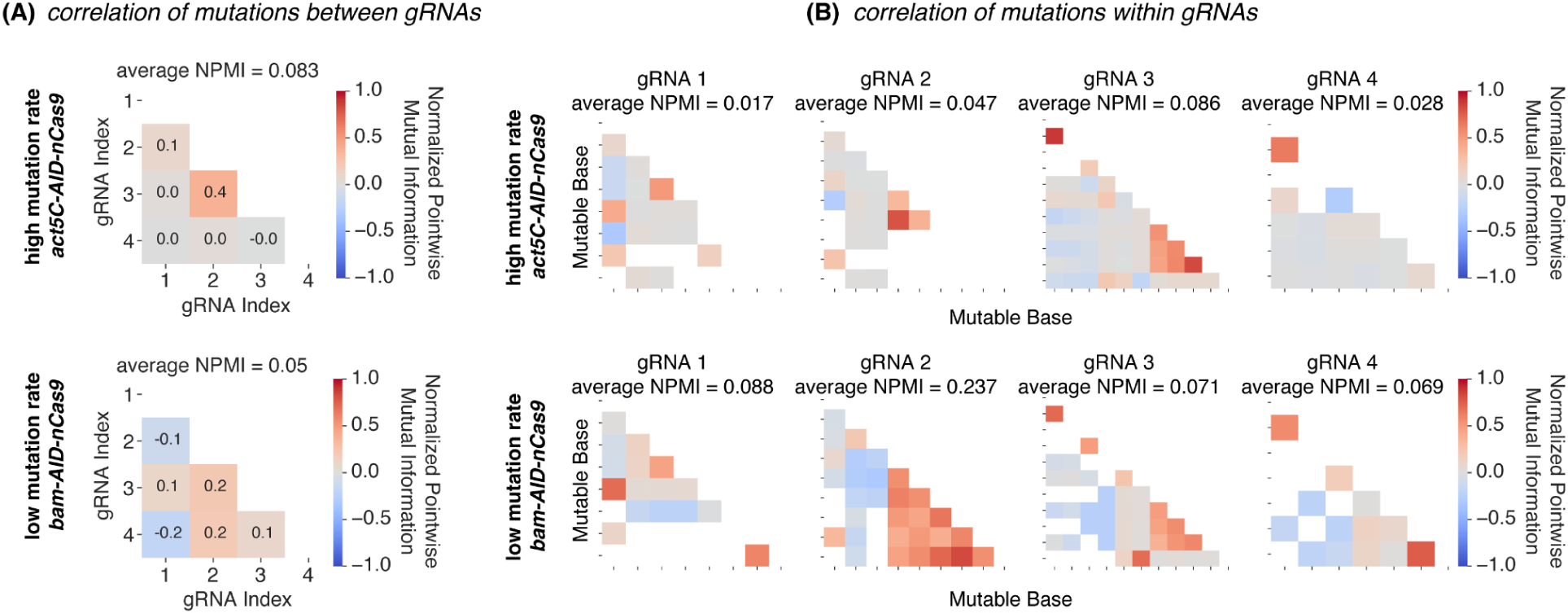
Quantifying the variability in mutations generated between and within gRNAs by mutual information. **(A)** Correlation in mutations between gRNAs for *act5C* (top) and *bam* (bottom) constructs quantified by mutual information. Heatmaps indicate the Normalized Pointwise Mutual Information (NPMI) between each pair of gRNAs, which measures the degree of co-occurrence between two events (0 if they co-occur by chance, 1 if completely correlated, -1 if completely uncorrelated). See Methods – Quantification of Mutational Variability for details. Mutations between gRNAs are mostly uncorrelated, with the exception of mutations between gRNAs 2 and 3 that show a small positive correlation. **(B)** Quantification of the correlation in mutations within each gRNA for *act5C* (top) and *bam* (bottom) constructs. Heatmaps show NPMI across each pair of mutable bases for each gRNA. The average NPMI is calculated across all values and serves as the correlation coefficient. The correlation between mutable bases in the same gRNA is close to 0 in most situations, as well as the average NPMI in a gRNA. Some exceptions are detected in gRNAs 2 and 3, where mutations in some positions are positively correlated.

**Figure 5 figure supplement 1.**
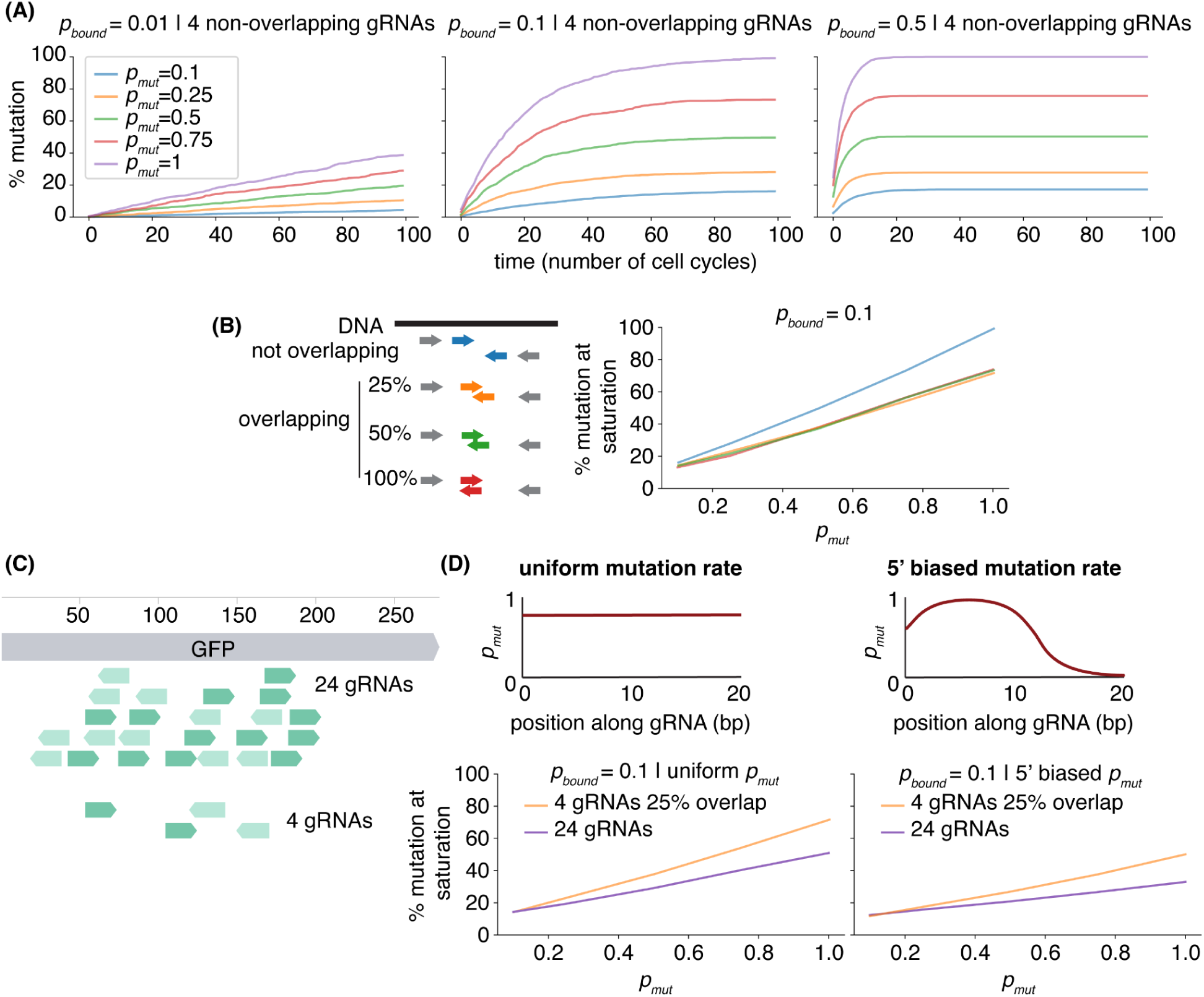
Simulating the effects of uniform and 5’ biased mutation rates. **(A)** Average fraction of mutations in all mutable bases at each generation when four non overlapping gRNAs with uniform mutation rates target GFP, for a range of *p_bound_*(left to right) and *p_mut_* (different colors) values. **(B)** Average fraction of mutations at the end of the simulation considering different amounts of gRNA overlap for *p_bound_* = 0.1 and different values of (different colors) and *p_mut_* (x-axis), in simulations using a uniform mutation rate along the gRNA. The same effect was observed with other *p_bound_* but *p_bound_* = 0.1 shown as an example. **(C)** Diagram of the different gRNA array constructs used to target our GFP segment. **(D)** Average fraction of mutations at the end of the simulation when 4 or 24 gRNAs target GFP for a range of *p_mut_*(x-axis), in simulations using a uniform (left) or 5’ biased (right) mutation rate along the gRNA obtained from the experimental profile of mutations (Figure 2 - figure supplement 3). All simulations consist of 200 cells, mutating over 100 cell cycles in panel A, or over 10000 in panels B and D, to ensure saturation.

**Figure 5 figure supplement 2.**
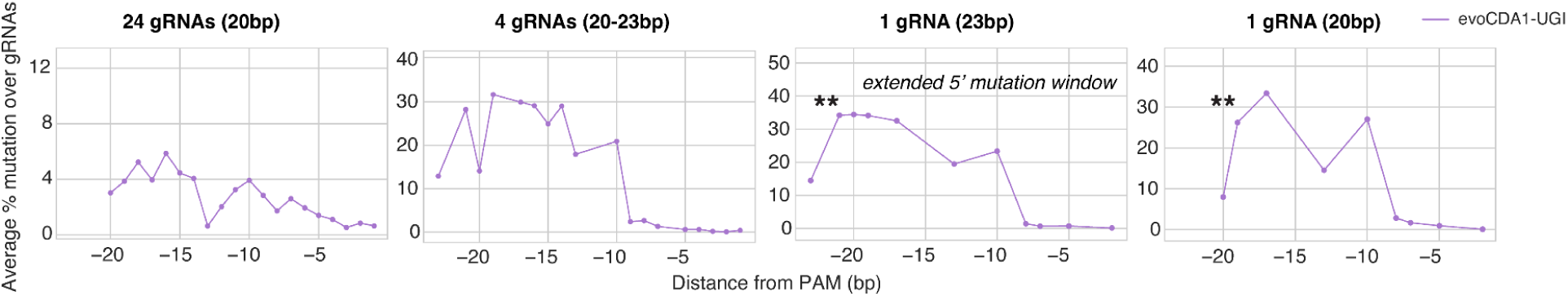
Testing different numbers and lengths of gRNA in cell culture. Average mutation profiles over all gRNAs used in the 24 gRNA, 4 gRNA, one 23 bp or 20 bp gRNA condition with *act5C-AID-nCas9*. 24 gRNAs (left) mutate along the whole 200 bp sequence and, when mutation rates are averaged over all gRNAs, the obtained mutation profiles exhibit less of a 5’ bias than when only 4 gRNAs are used. ** A 23 bp gRNA extends the mutation window 5’ compared to the 20bp version.

**Figure 5 figure supplement 3.**
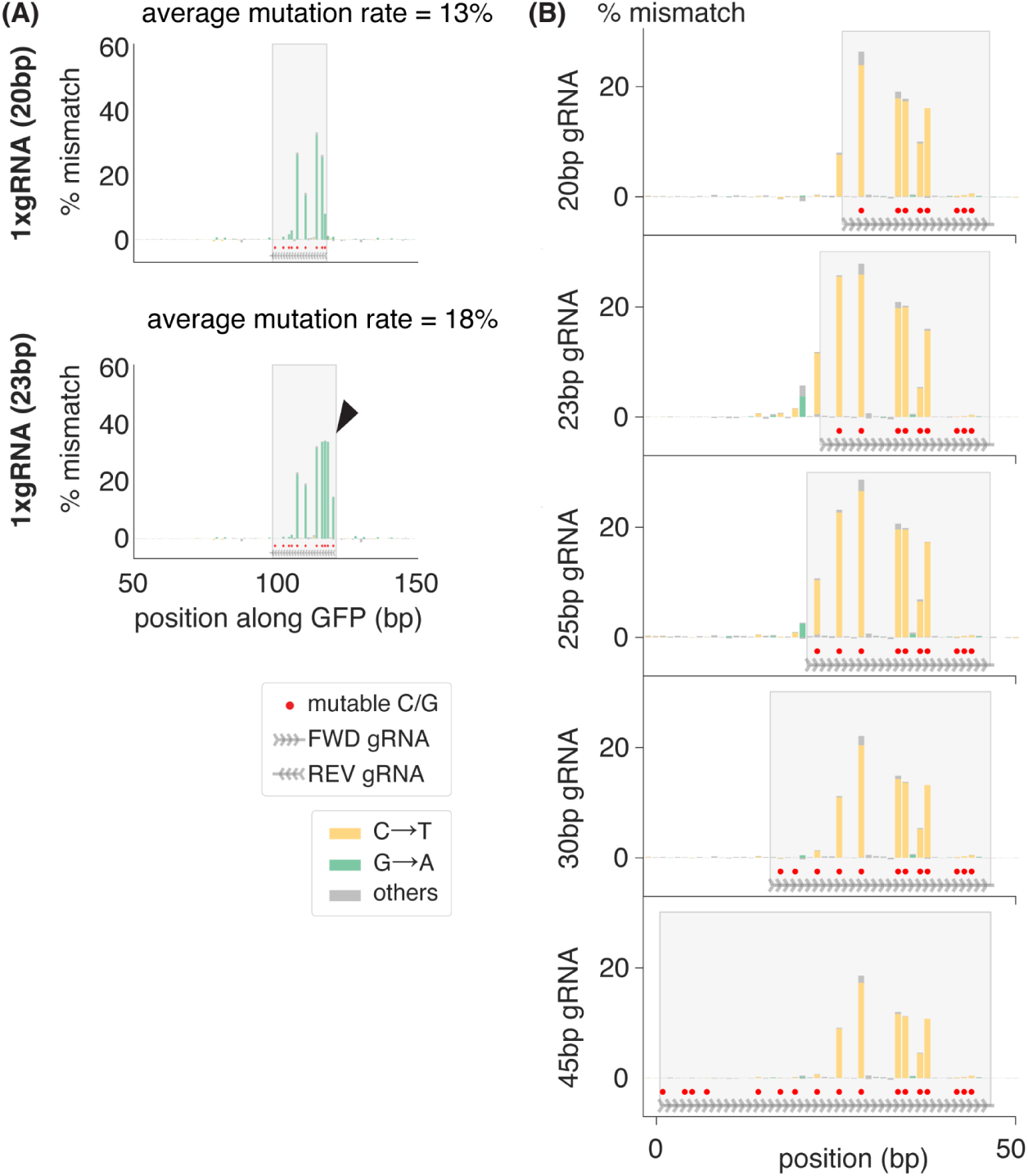
Increasing gRNA length beyond 23 bp does not further extend the mutation window. **(A)** Mismatches to GFP classified by base substitution in Kc cells transfected with *act5C-AID-nCas9* and one 20 bp (top) or 23 bp (bottom) gRNA, 2 weeks post-transfection. Arrowhead and grey box indicate the 5’ extended mutation window with a 23bp gRNA. **(B)** Mutations produced in GFP classified by base substitution in Kc cells transfected with *act5C-AID-nCas9* and one gRNA of 20, 23, 25, 30 or 45 bp in length, 1 week post-transfection.

**Figure 6 figure supplement 1.**
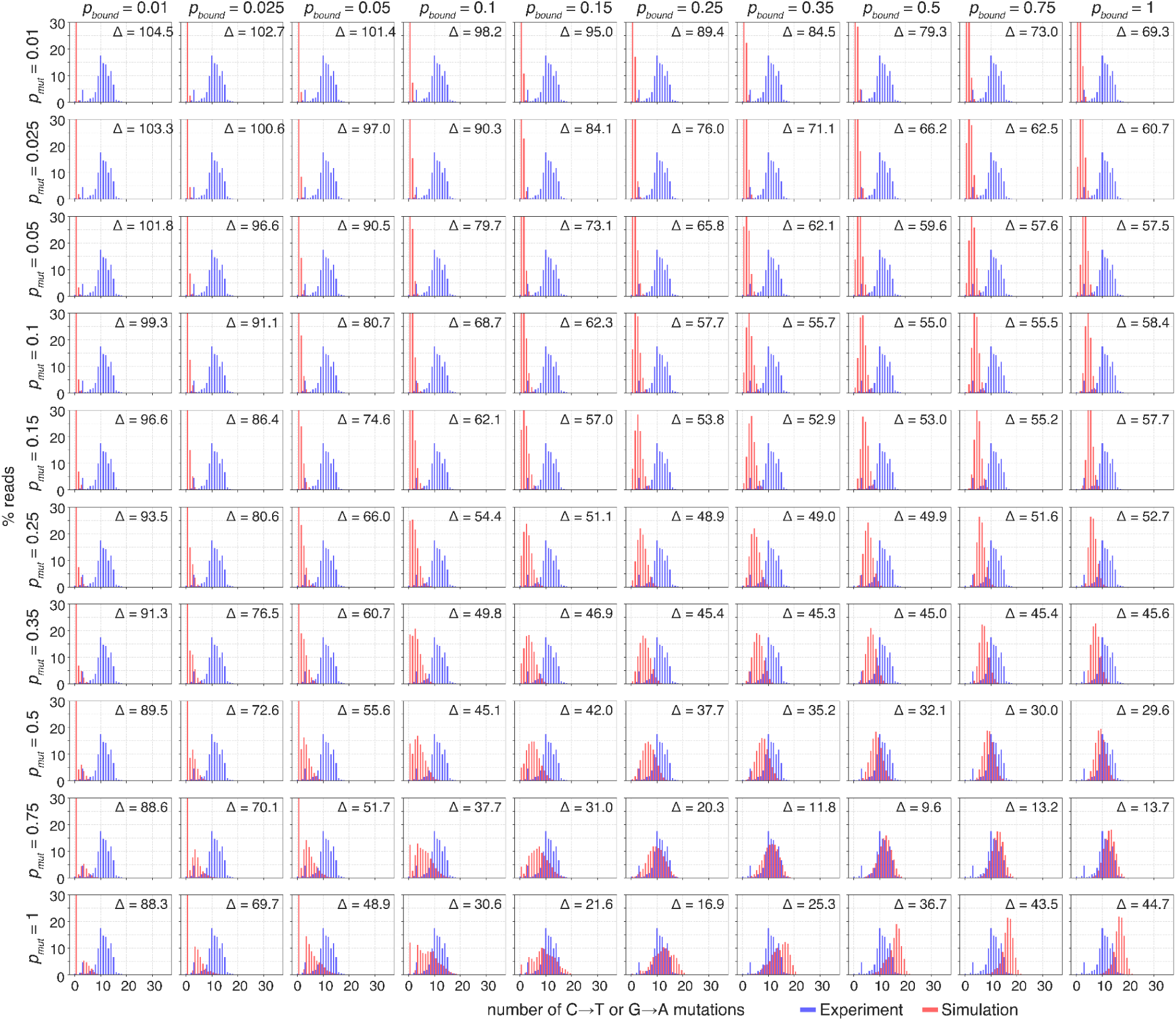
Fits of our simulation to experimental data for *act5C-AID-nCas9*. Comparison of simulated histograms of number of introduced mutations per DNA molecule (red) as we systematically vary *p_bound_* (along columns) and *p_mut_*values (along rows) with the experimental data obtained from the high mutation rate experiment in embryos (blue, *act5C-AID-nCas9,* Figure 3 - figure supplement 3B). Δ indicates the euclidean distances between both histograms on each plot. Each simulated histogram is obtained from 1000 simulated DNA molecules after 10 cell cycles using the indicated *p_mut_* and *p_bound_* parameters. The number of mutations generated in each molecule is then counted and plotted as a histogram.

**Figure 6 figure supplement 2.**
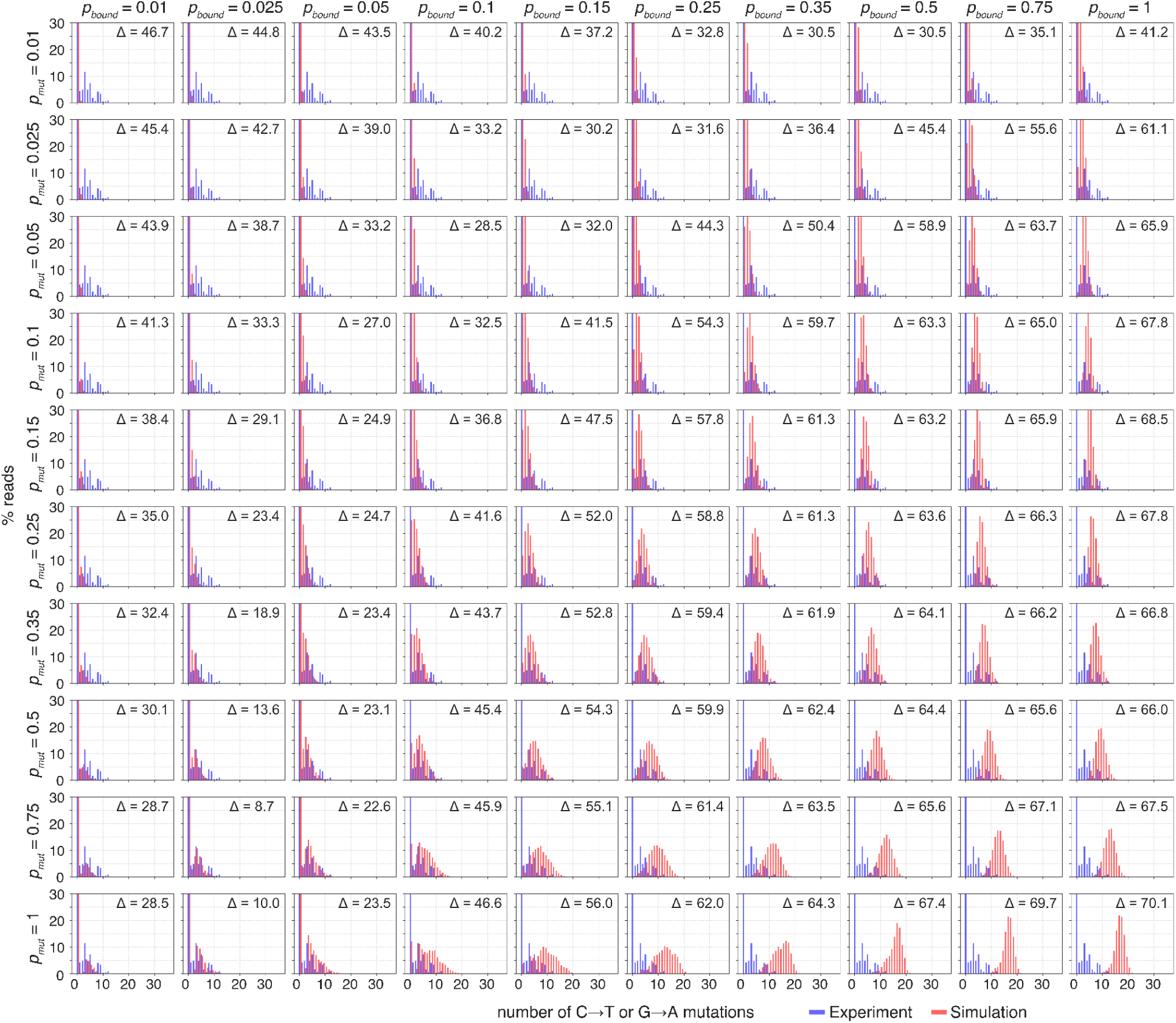
Fits of our simulation to experimental data for *bam-AID-nCas9-UGI*. Comparison of simulated histograms of number of introduced mutations per DNA molecule (red) as we systematically vary *p_bound_* (along columns) and *p_mut_*values (along rows) with the experimental data obtained from the high mutation rate experiment in embryos (blue, *bam-AID-nCas9-UGI,* Figure 3 - figure supplement 3B). Δ indicates the euclidean distances between both histograms on each plot. Each simulated histogram is obtained from 1000 simulated DNA molecules after 10 cell cycles using the indicated *p_mut_* and *p_bound_* parameters. The number of mutations generated in each molecule is then counted and plotted as a histogram.

**Figure 6 figure supplement 3.**
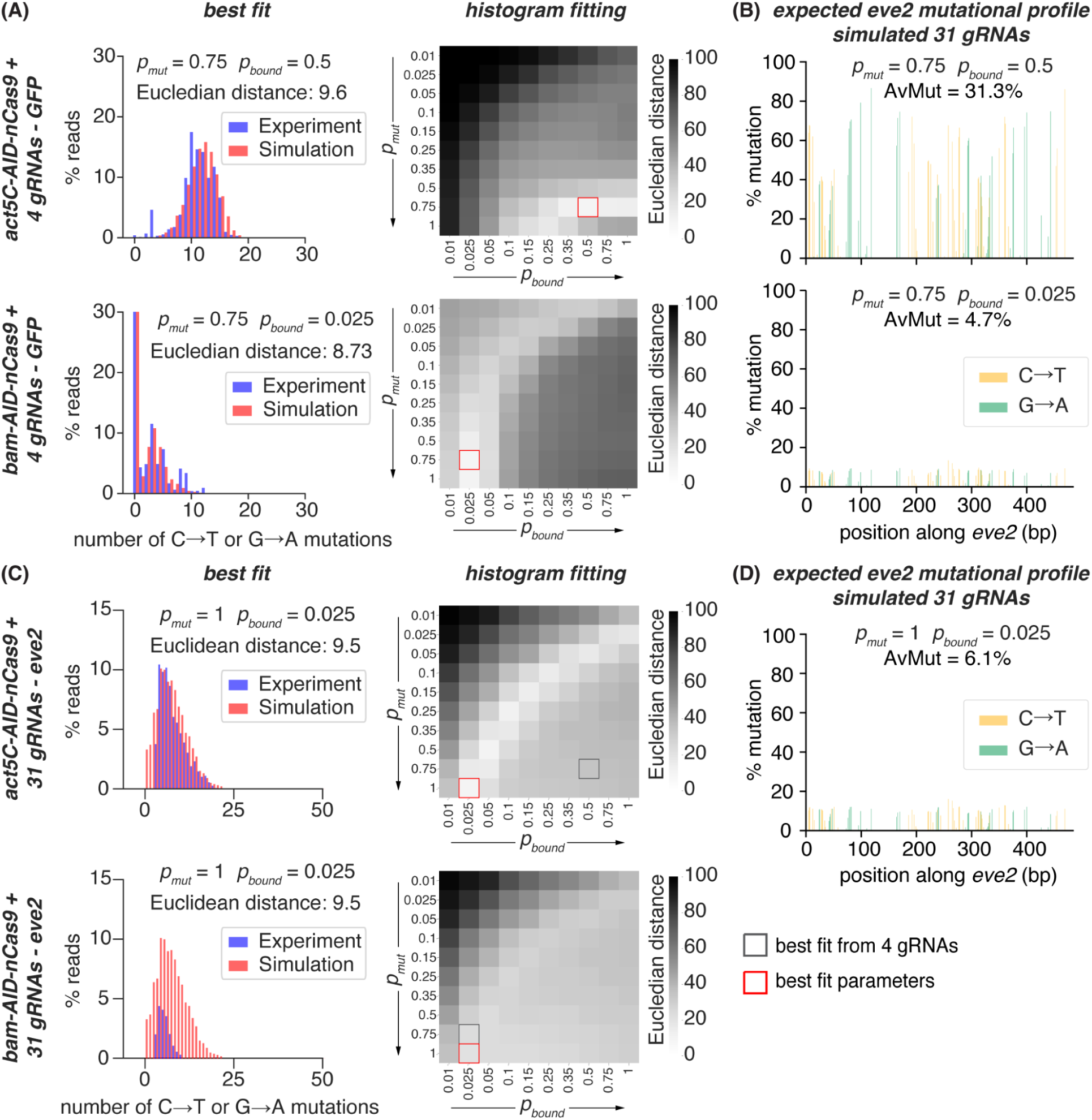
Fitting in vivo mutations in *eve2* to simulations. **(A)** Finding the best fit parameters between simulations and experiments in embryos using the *act5C* or *bam* promoters (Figure 3A, B) by comparing the histograms of the distribution of number of mutations per read. Left plots show best fits for the *act5C* and *bam* experiments, with the experimental distribution of number of mutations shown in blue and best fit from the simulations in red. Left heatmaps show Euclidean distances between the two histograms for 100 combinations of *p_bound_* and *p_mut_* , with the lowest value marked in red (Methods - Fitting simulated data to experiments). Tested *p_bound_* and *p_mut_*values and fits for all combinations are shown in Figure 6 - figure supplement 1 and Figure 6 - figure supplement 2. **(C)** Finding the best fit parameters between simulations and experiments in embryos using the *act5C* promoter and 20 bp gRNAs targeting *eve2* by comparing the histograms of the distribution of number of mutations per read as in Figure 5B, and Figure 5 - figure supplements 2 and 3. Left heatmap shows Euclidean distances between the two histograms for 100 combinations of *p_bound_* and *p_mut_* with the lowest value in red. Right plot shows the best fit, with the experimental distribution of the number of mutations shown in blue and the best fit from the simulations shown in red. **(D)** Expected mutational profile of *eve2* using the best fit parameters from the 4 gRNAs targeting GFP (left) and from the 31x 20bp gRNA arrays targeting *eve2* assuming best fit parameters from the experiment in (A) (right). Plots indicate the average of 1000 mutated simulated reads.

**Figure 6 figure supplement 4.**
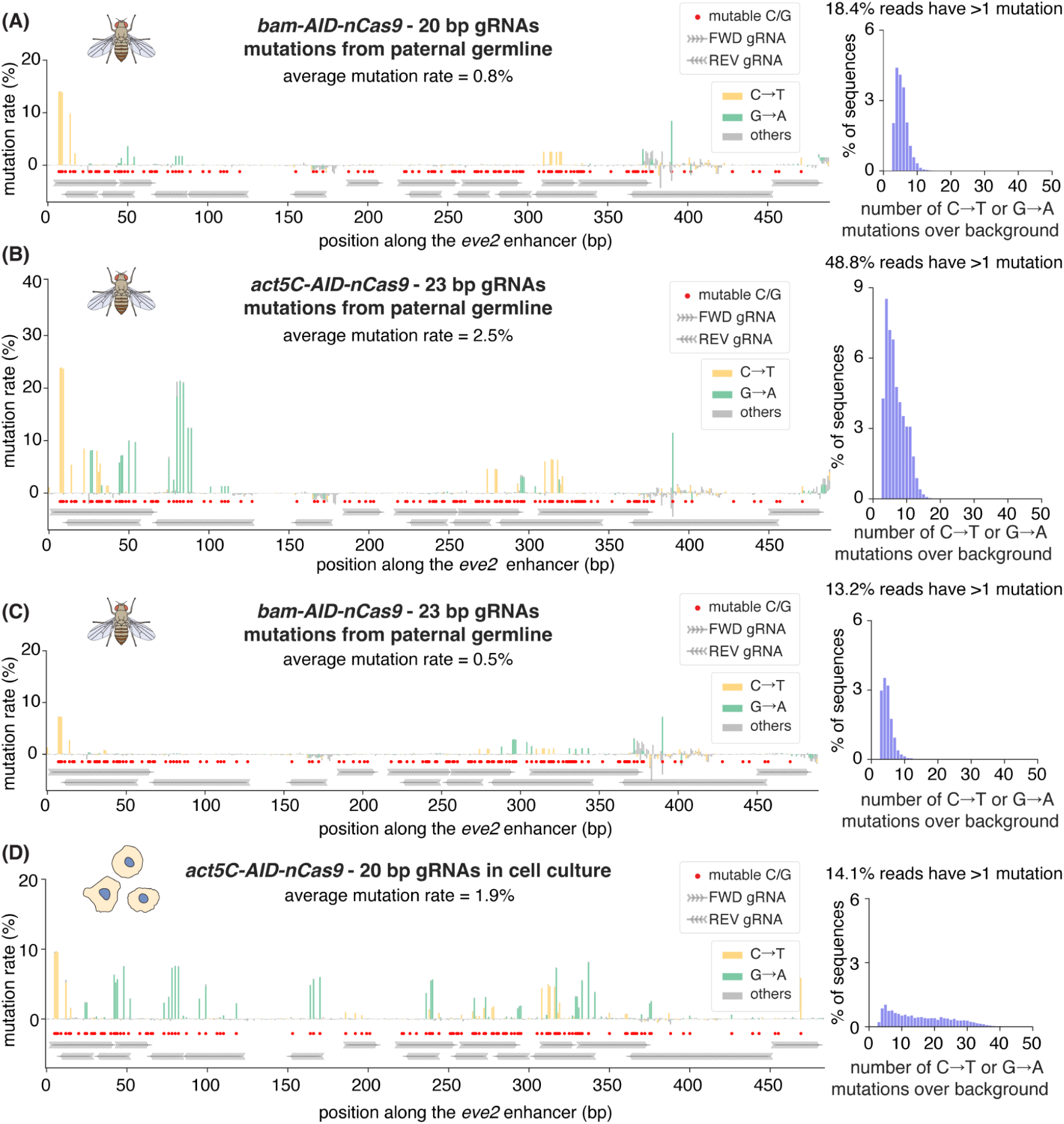
Testing different promoters and gRNA lengths to target an enhancer in vivo, and comparison with cell culture. (A) Average mutation rate (left) and distribution of mutations (right) generated in the paternal germline by *bam-AID-nCas9* and 31 gRNAs of 20 bp in length targeting *eve2*. **(B-C)** Average mutation rate (left) and distribution of mutations (right) generated in the paternal germline by *act5C-AID-nCas9* (B) or *bam-AID-nCas9* (C) and 31 gRNAs of 23 bp in length targeting *eve2* (D) Average mutation rate (left) and distribution of mutations (right) generated in S2 cells in culture 7 days post transfection by *act5C-AID-nCas9* and 31 gRNAs of 20 bp in length targeting *eve2* (left) . Red dots mark all “mutable bases” (Cs on FWD gRNAs and Gs on REV gRNAs). Histograms in all panels show the distribution of observed number of mutations per read after subtracting the number of mismatches observed in control samples in each condition.

**Figure 6 figure supplement 5.**
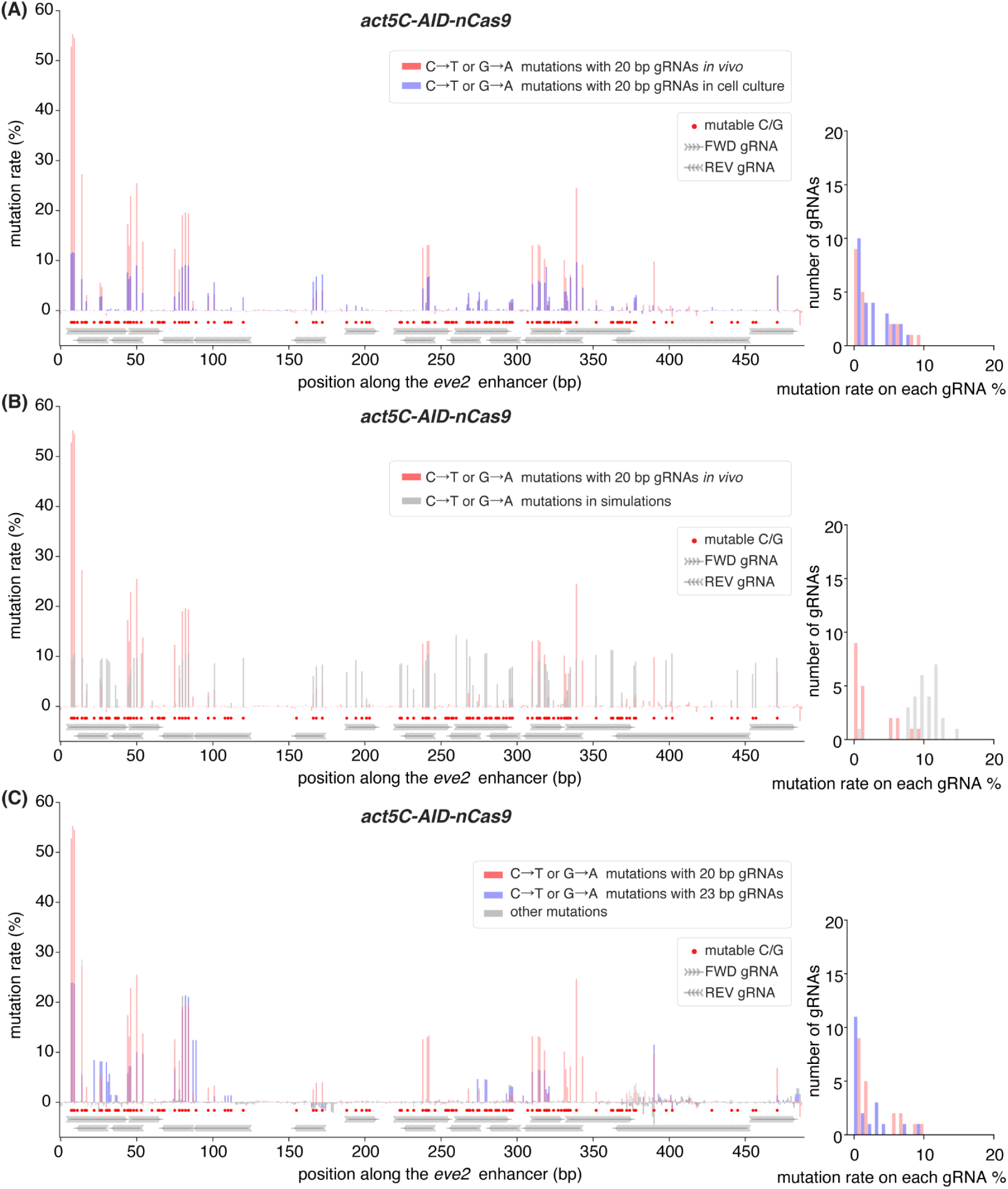
Variability in mutations generated by gRNAs in vivo. **(A)** Comparison between the mutation rates obtained by targeting *eve2* with *act5C-AID-nCas9* and 20 bp gRNAs in vivo (red, same data as Figure 6F) or in cell culture (blue, same data as Figure 6 - figure supplement 4D). **(B)** Comparison between the expected mutation rates from simulations (grey, same data as in Figure 6 - figure supplement 3D) using the best fit parameters to the distribution of mutations obtained by *act5C-AID-nCas9* with 20 bp gRNAs targeting *eve2* compared to the actual mutations obtained by *act5C-AID-nCas9* with 20 bp gRNAs (red, same data as Figure 6F). **(C)** Comparison between the mutation rates obtained by targeting *eve2* with *act5C-AID-nCas9* and 20 bp gRNAs (red, same data as Figure 6F) and 23 bp gRNAs (blue, same data as Figure 6 - figure supplement 4B). Right histograms in all panels show the distribution of mutation rates calculated for each gRNA. To obtain these average mutation rates per gRNA without having gRNA overlaps influence the calculation, we limited the number of bases considered for each gRNA. Specifically, in line with the mutation bias observed in Figure 2 - figure supplement 3, we only considered the mutation rates on Cs in the first 10 bp of each forward gRNA or Gs on the first 10 bp of each reverse gRNA. The mutation rate for each gRNA was the average of the rates for these selected mutable bases.

**Figure 6 figure supplement 6.**
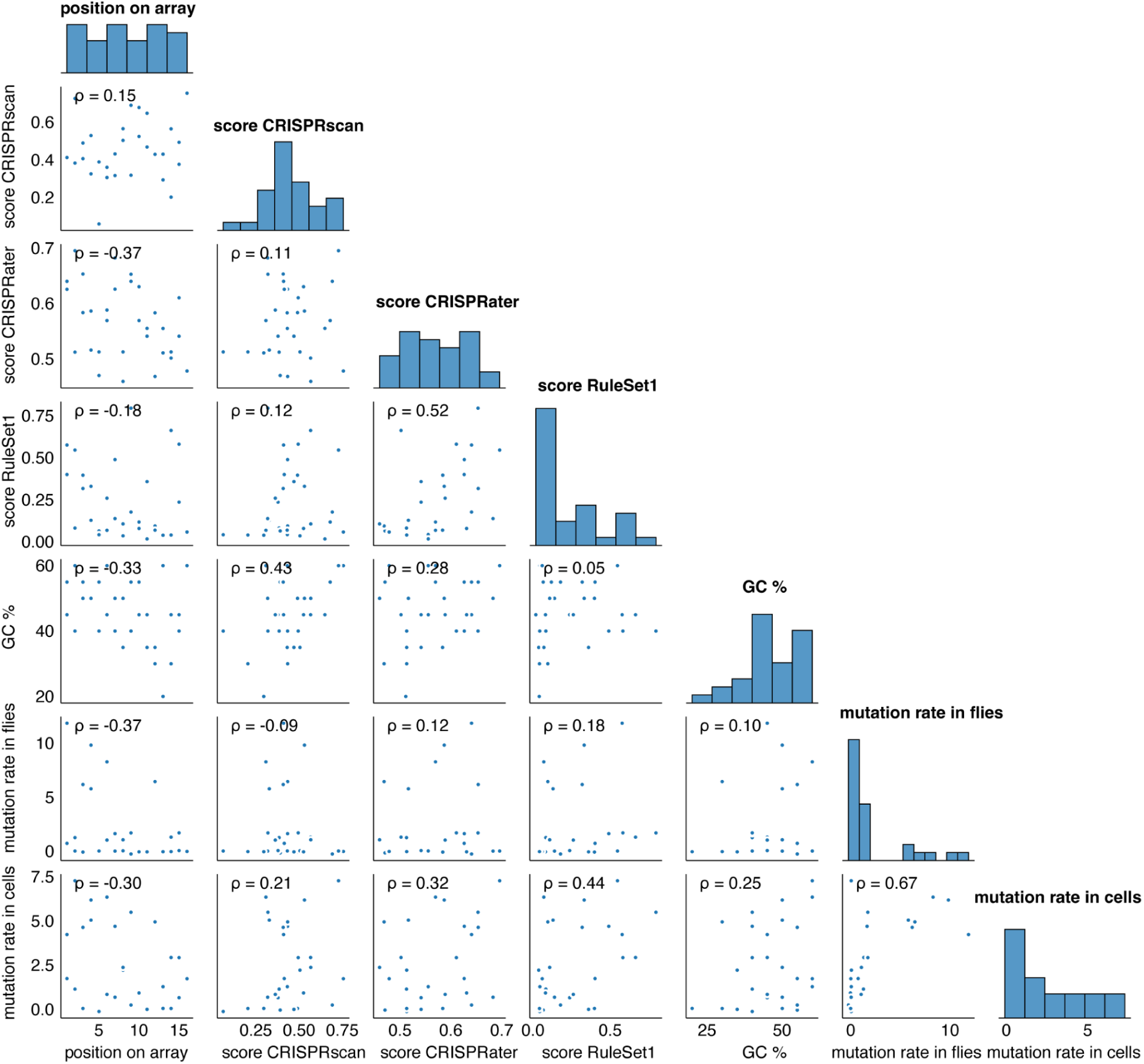
Correlation between average mutation rates on each gRNA, scores from different protospacer design tools and position on the gRNA array. Each plot shows the correlation between two variables: mutation rates in each of the 31 gRNAs targeting *eve2* in cell culture or in vivo, and different properties of each gRNA: position on the gRNA array, binding scores calculated by CRISPRScan, CRISPRater or RuleSet1, and CG percentage. Histograms on the diagonal show the distribution of each of the quantified parameters. No clear correlation is detected between mutation rates in cells or embryos, and any of the calculated parameters (%GC, binding scores, position on the array - bottom two rows). Of note, different gRNA scoring methods also show very little correlation between each other.

**Figure 7 figure supplement 1.**
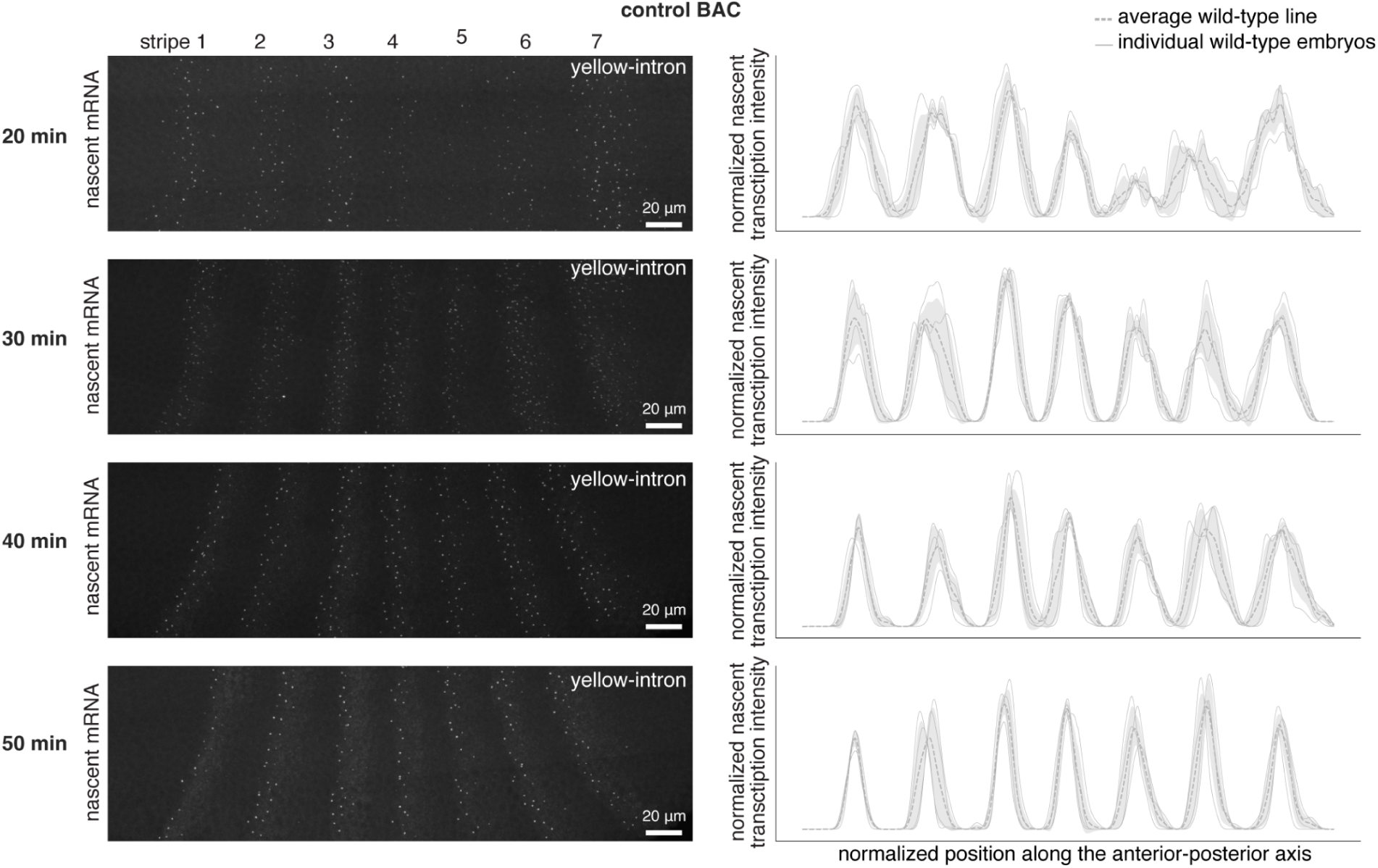
*BAC-eve-MS2-yellow* expression over time. Expression of *eve-MS2-yellow* measured by smFISH and quantified nascent transcription profiles (smoothed smFISH intensity of the nascent transcription spot summed at each AP position after straightening the stripes and normalizing for the average of all stripe peaks except stripe 2, Methods - Image analysis and gene expression quantification) in the wild type BAC. Thick grey lines and shaded areas indicate mean and SD of expression profiles at the indicated timepoints. Thin grey lines indicate the expression profile of individual embryos.

**Figure 7 figure supplement 2.**
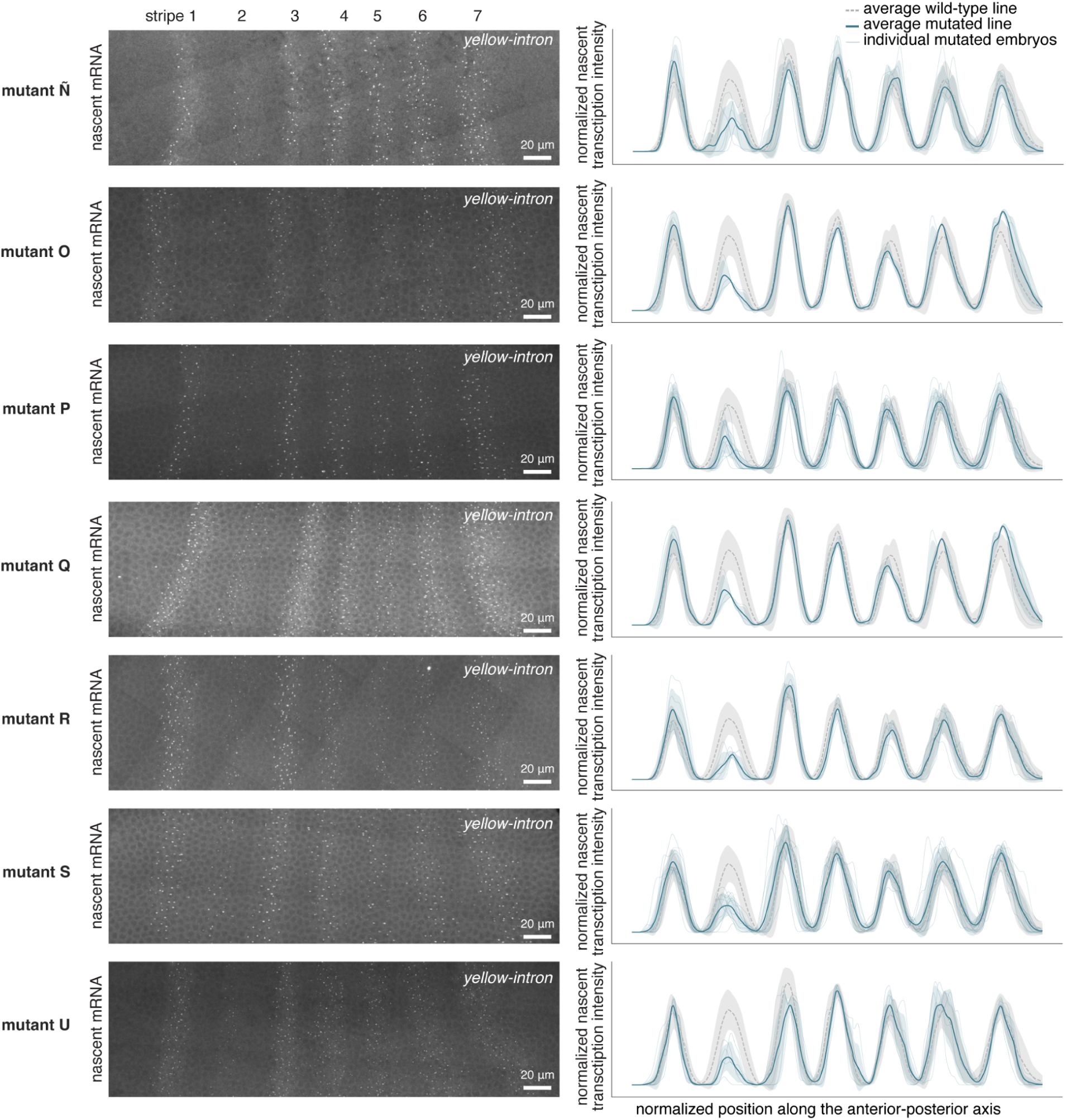
Stable BAC lines with ‘strong decrease’ phenotypes. Expression of *eve-MS2-yellow* measured by smFISH and quantified nascent transcription profiles (smoothed smFISH intensity of the nascent transcription spot summed at each AP position after straightening the stripes and normalizing for the average of all stripe peaks except stripe 2, Methods - Image analysis and gene expression quantification) in stable lines with mutations that lead to a phenotypic change classified as ‘low expression’. Thick blue lines and shaded areas indicate mean and SD of expression profiles from the indicated line at 30 min into nc14, respectively. Thick grey lines and shaded areas indicate mean and SD of expression profiles at the same timepoint in control embryos, respectively. Thin blue lines indicate the expression profile of individual embryos.

**Figure 7 figure supplement 3.**
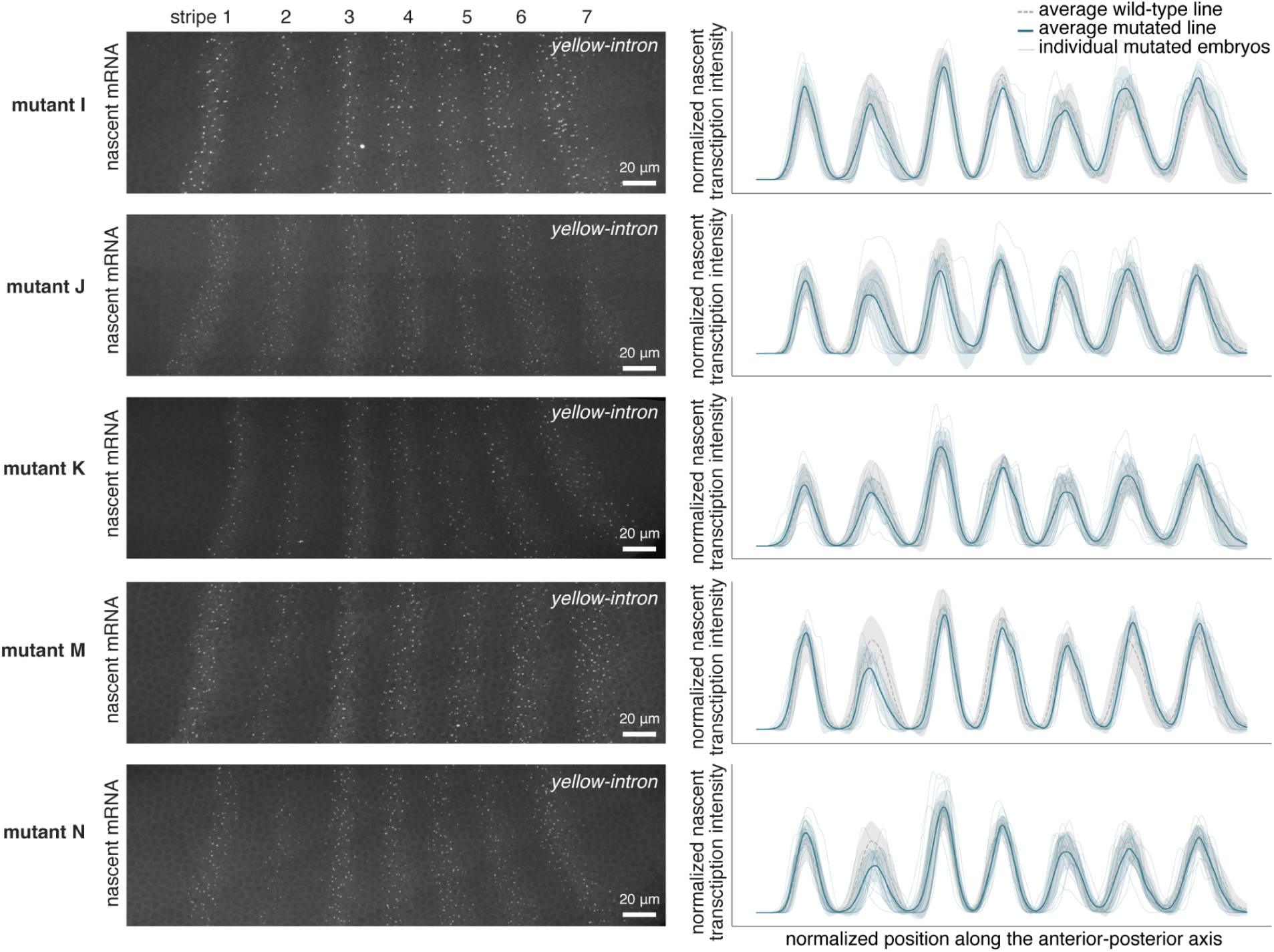
Stable BAC lines with ‘mild decrease’ phenotypes. Expression of *eve-MS2-yellow* measured by smFISH and quantified nascent transcription profiles (smoothed smFISH intensity of the nascent transcription spot summed at each AP position after straightening the stripes and normalizing for the average of all stripe peaks except stripe 2, Methods - Image analysis and gene expression quantification) in lines with mutations that lead to a phenotypic change classified as ‘mild decrease’. Thick blue lines and shaded areas indicate mean and SD of expression profiles from the indicated line at 30 min into nc14, respectively. Thick grey lines and shaded areas indicate mean and SD of expression profiles at the same timepoint in control embryos, respectively. Thin blue lines indicate the expression profile of individual embryos.

**Figure 7 figure supplement 4.**
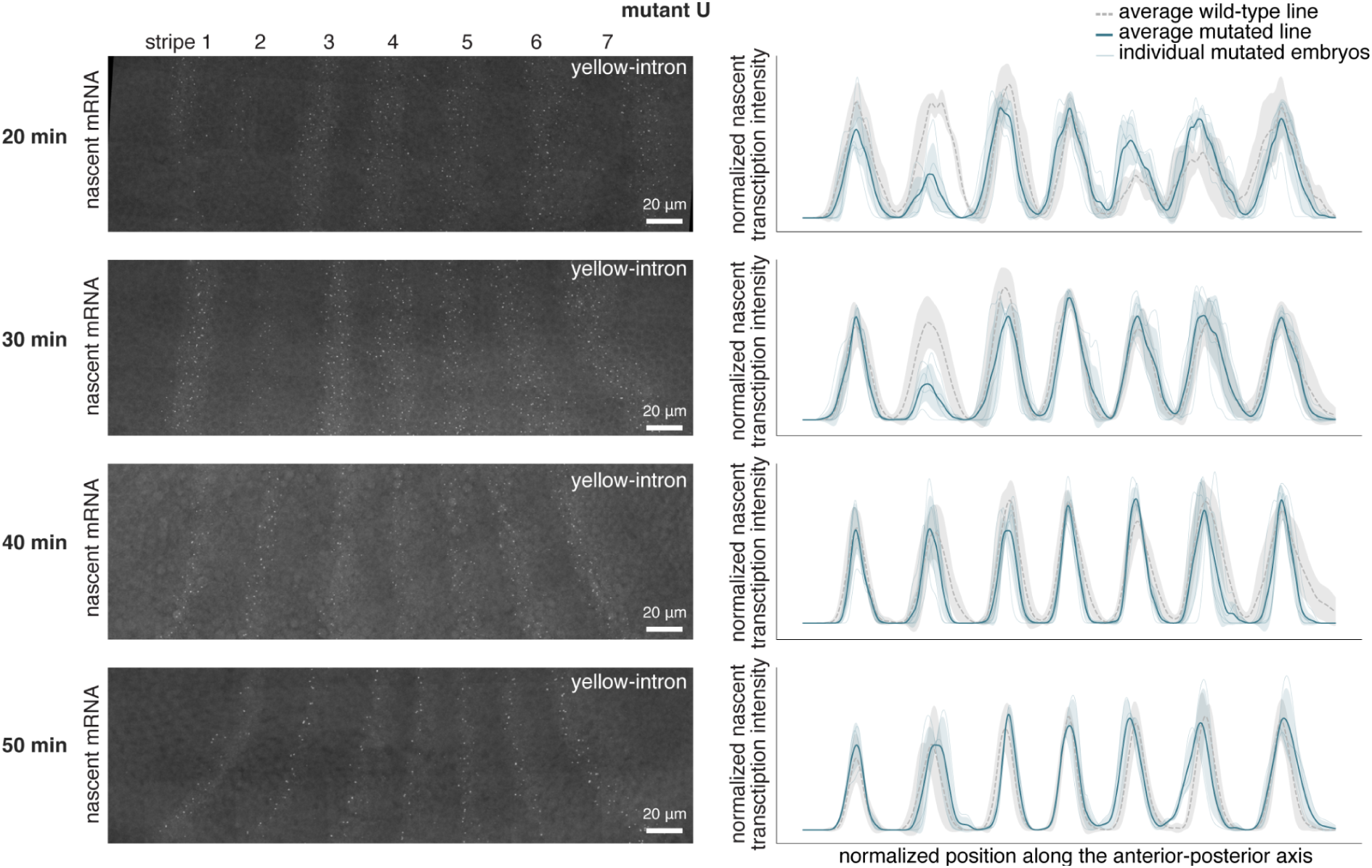
Example of a stable BAC line (mutant U) with ‘strong decrease’ phenotype that returns to wild type levels of expression at the end of nuclear cycle 14. Expression of *eve-MS2-yellow* measured by smFISH and quantified nascent transcription profiles (smoothed smFISH intensity of the nascent transcription spot summed at each anterior-posterior position after straightening the stripes and normalizing for the average of all stripe peaks except stripe 2, Methods - Image analysis and gene expression quantification) in a stable line (BAC_U) in the ‘strong decrease’ category at early timepoints that returns to expression levels similar to wild-type at 40 and 50 min into nuclear cycle 14. Thick blue lines and shaded areas indicate mean and SD of expression profiles at the indicated timepoints. Thick grey lines and shaded areas indicate mean and SD of expression profiles at the same timepoint in control embryos, respectively. Thin blue lines indicate the expression profile from individual embryos.

**Figure 7 figure supplement 5.**
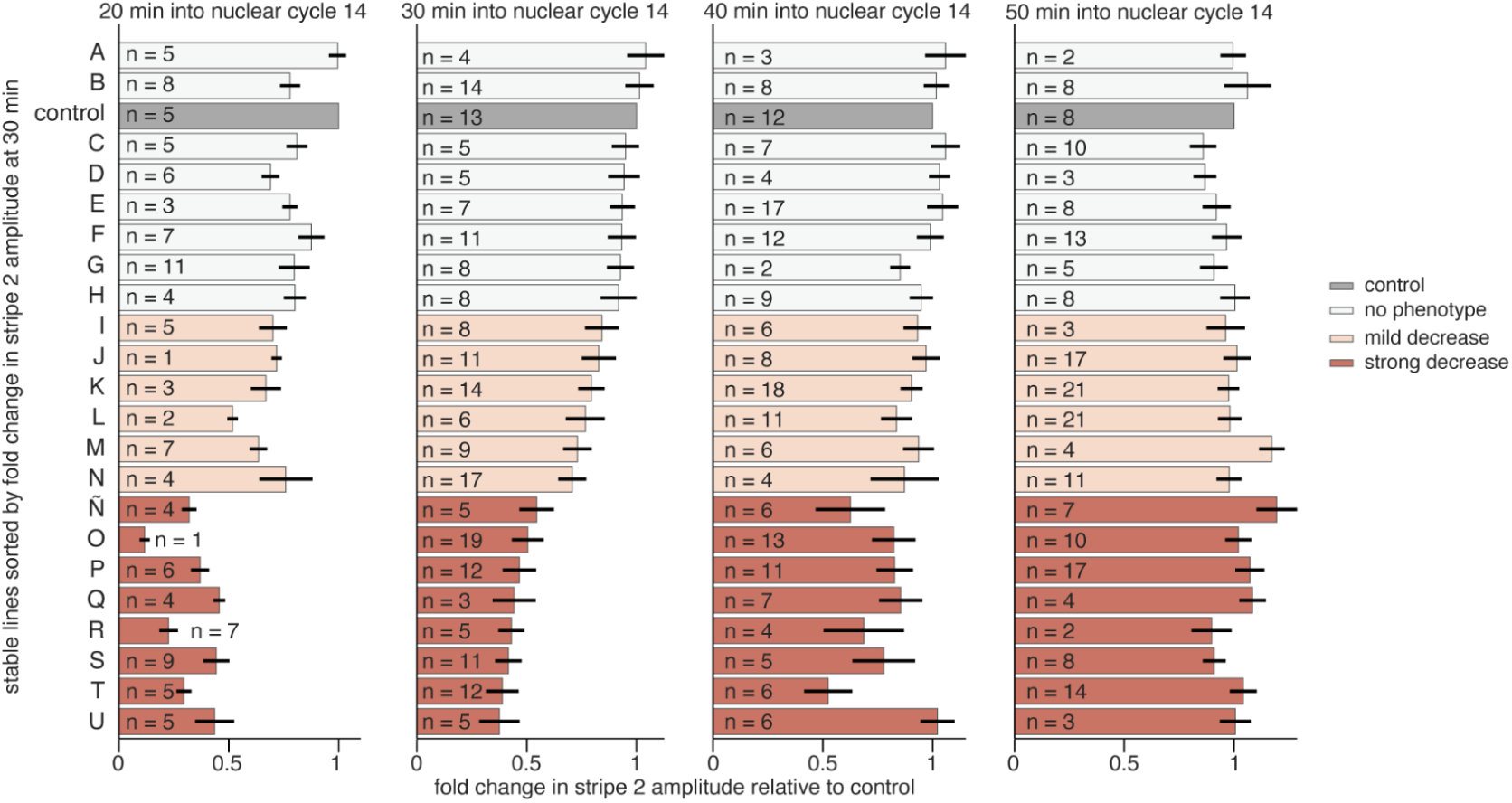
Changes in stripe 2 amplitude over time. Average fold-change in stripe two amplitude (peak of the second stripe in nascent transcription profiles) compared to controls and standard error of the mean shown for each line at the 4 indicated timepoints. Lines are sorted by fold-change at the 30 min timepoint and colored based on the three defined categories: no phenotype, mild decrease and low expression. n indicates the number of embryos imaged for each line at each timepoint.

**Figure 7 figure supplement 6.**
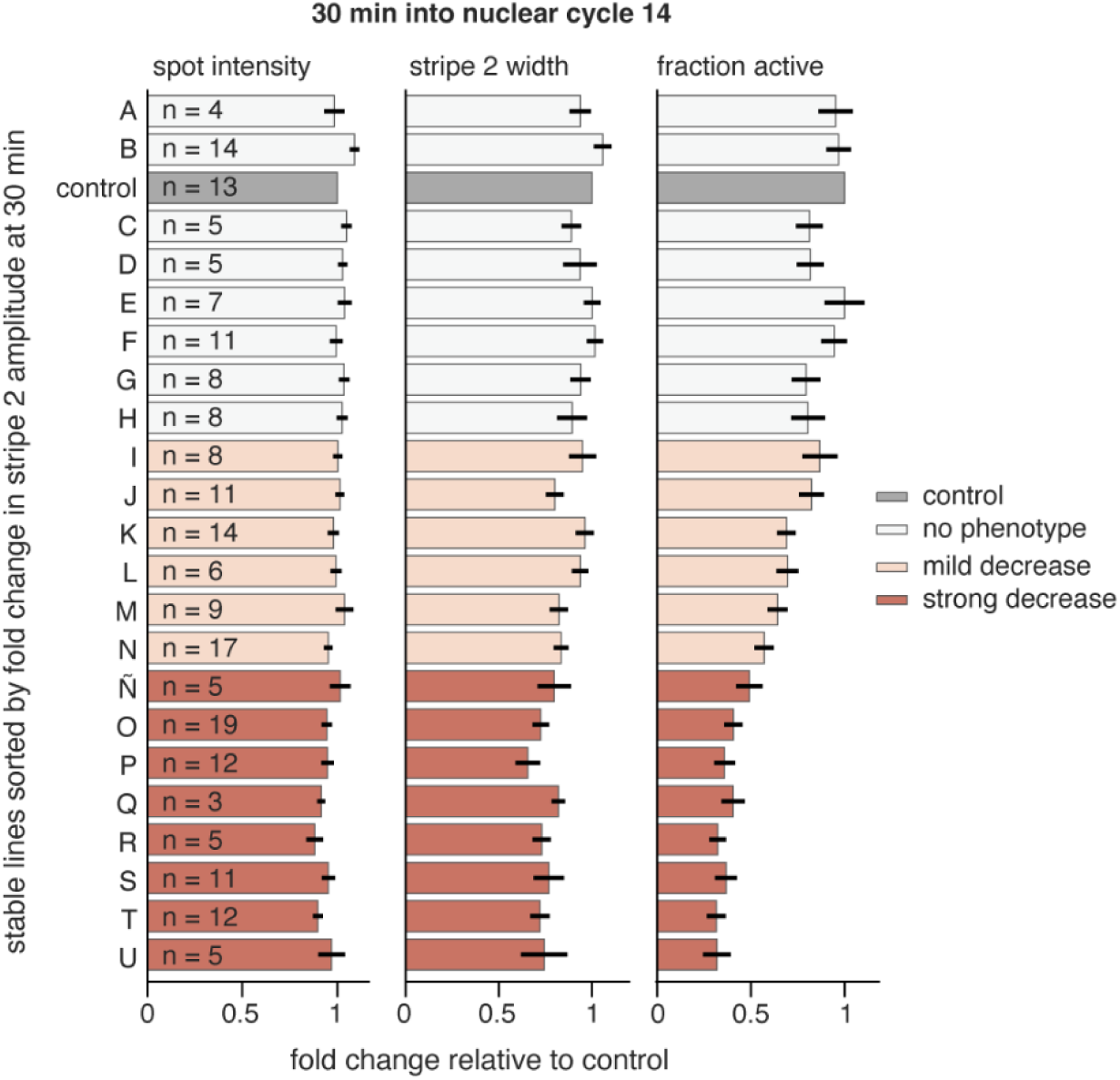
Transcription parameters impacting reduced levels of *eve2* activity. Detailed analysis of parameters impacting *eve2* activity at 30 min into nc14 in our 22 lines. Spot intensity remains relatively constant across the different mutations, but the number of active nuclei in stripe 2 is greatly reduced in some lines. Lines with a decreased fraction of active nuclei exhibit a small reduction in stripe 2 width as well. n indicate the number of embryos imaged for each line at 30 min into nuclear cycle 14. Average fold-change in each parameter compared to controls and standard error of the mean shown.

**Figure 7 figure supplement 7.**
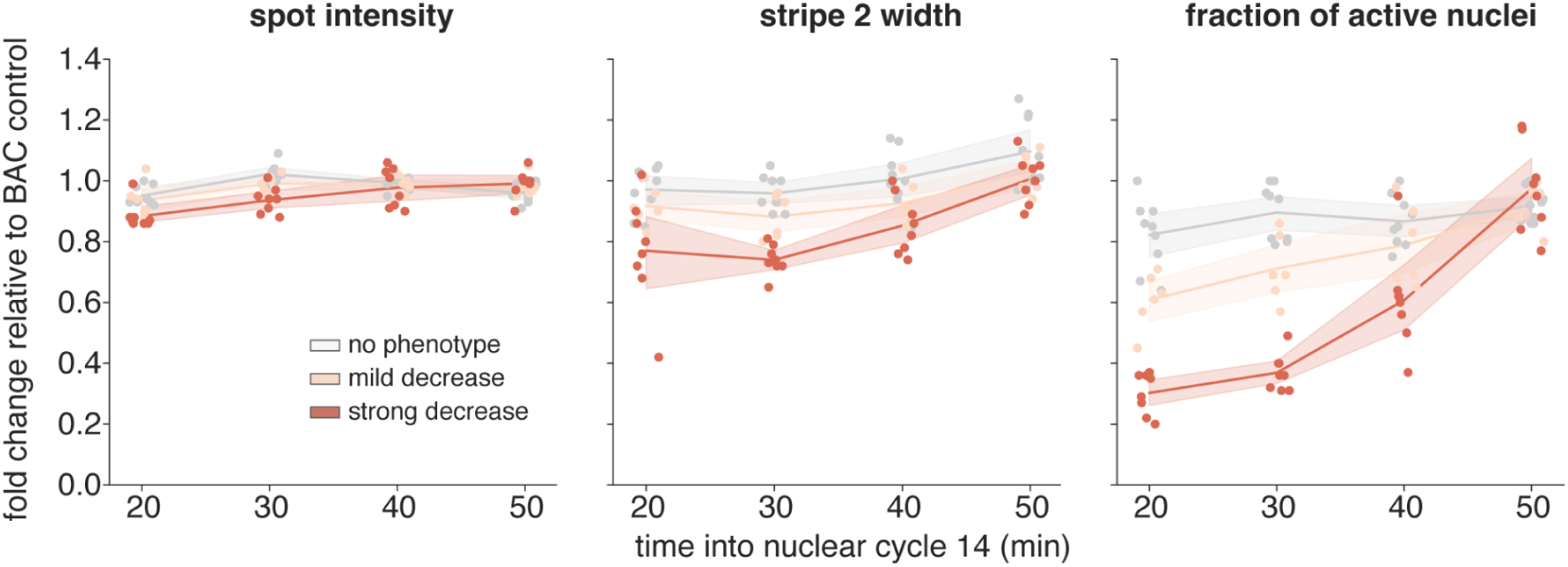
Changes in transcription parameters over time for the three phenotypic categories. Spot intensity (left) quantifies the average intensity of each detected nascent transcription spot in stripe 2 relative to stripe 3. Stripe 2 width (middle) quantifies the stripe width from the positions along the AP axis of all detected stripe 2 spots. Fraction of nuclei active (right) calculates the number of spots assigned to stripe 2 relative to stripe 3. Dots indicate the average fold change for all each mutated line. Lines and shaded areas indicate mean and 95% confidence intervals for each category at each timepoint.

**Figure 7 figure supplement 8.**
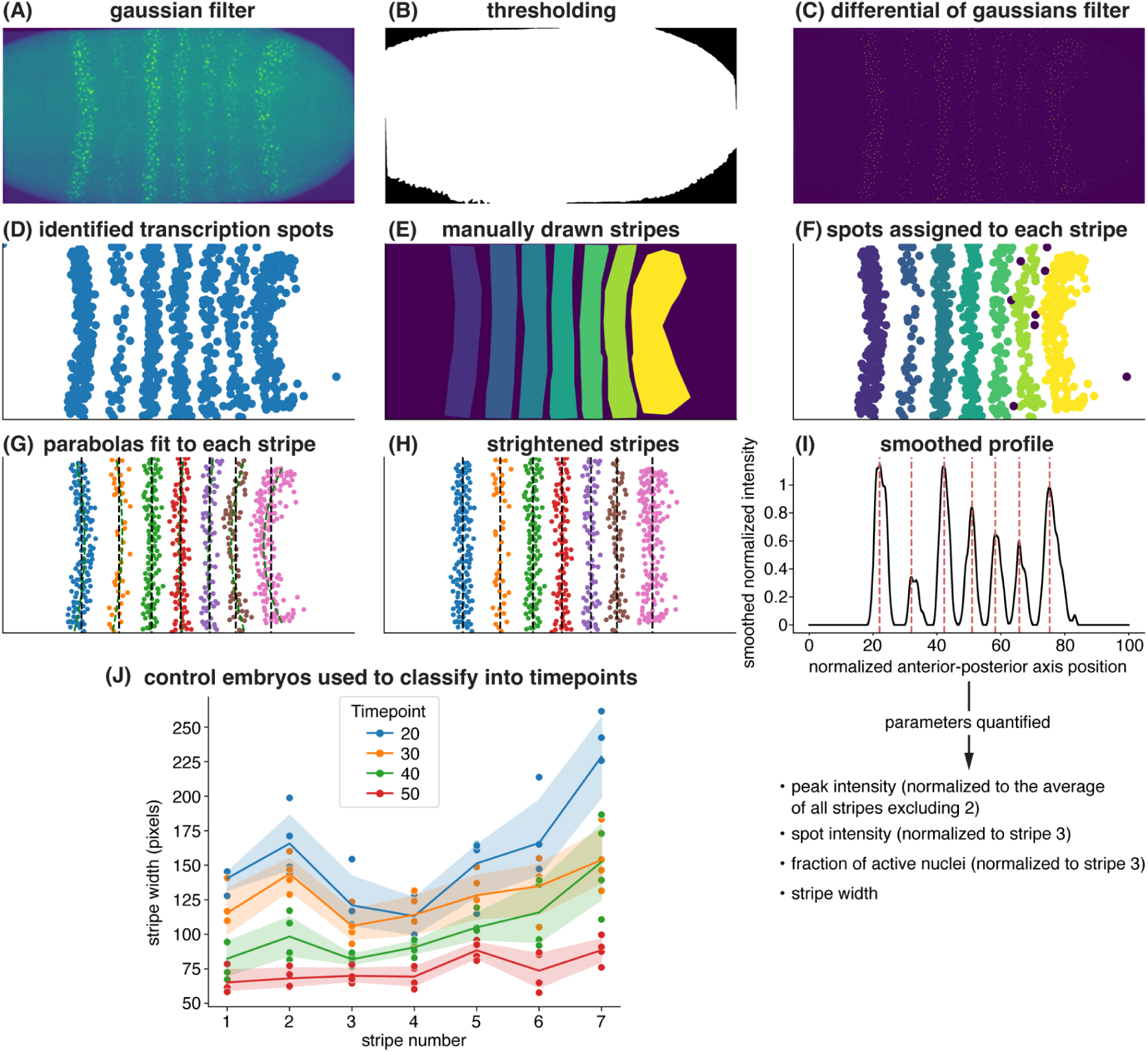
Image analysis and quantification pipeline. **(A)** An example image of an embryo at 30 min into nuclear cycle14, showing reduced expression of *eve* stripe 2. Images are rotated by aligning the fourth stripe and flipped so that the anterior is left and dorsal is up. **(B)** A mask is generated to define the embryo boundaries. **(C)** A difference of Gaussians filter is applied to detect fluorescent puncta corresponding to sites of nascent transcript formation as reported by *eve-MS2-y* using smFISH. **(D)** Detected spots are identified and plotted. **(E)** Stripe boundaries are manually drawn using a lasso tool. **(F)** Spots are binned and assigned to the corresponding stripes. **(G)** The positions of spots within each stripe are fitted to a quadratic function. **(H)** Stripes are straightened by setting the average x-positions of the spots as the center of the stripe. Then, the y-position for each spot is maintained and the x-position shifted to keep the same distance from the original x-spot to the parabola as to the new center of the stripe. **(I)** From the aligned spots, a smoothing Gaussian filter of 20 pixels is calculated to generate the expression profile. Quantified parameters include peak intensity (normalized to the average of all peaks excluding the one corresponding to stripe 2), stripe width, spot intensity (normalized to stripe 3), and fraction of active nuclei (normalized to stripe 3). All parameters are analyzed for experimental and control embryos using the same pipeline, allowing direct comparison between conditions. X–Y axes are labeled in pixels for (A–H). **(J)** Graph plotting changes in stripe width over time in BAC control embryos. Stripe width is plotted versus stripe number and classified by timepoints (20, 30, 40 or 50 min into nuclear cycle 14). These distributions were used to fit other embryos into the different temporal categories. Lines represent the average across embryos of the same time point, and shaded regions represent the 95% confidence interval (1.96 * standard error of the mean). y-axis shows stripe width in pixels, with resolution corresponding to 0.396 µm/pixel.

**Figure 8 figure supplement 1.**
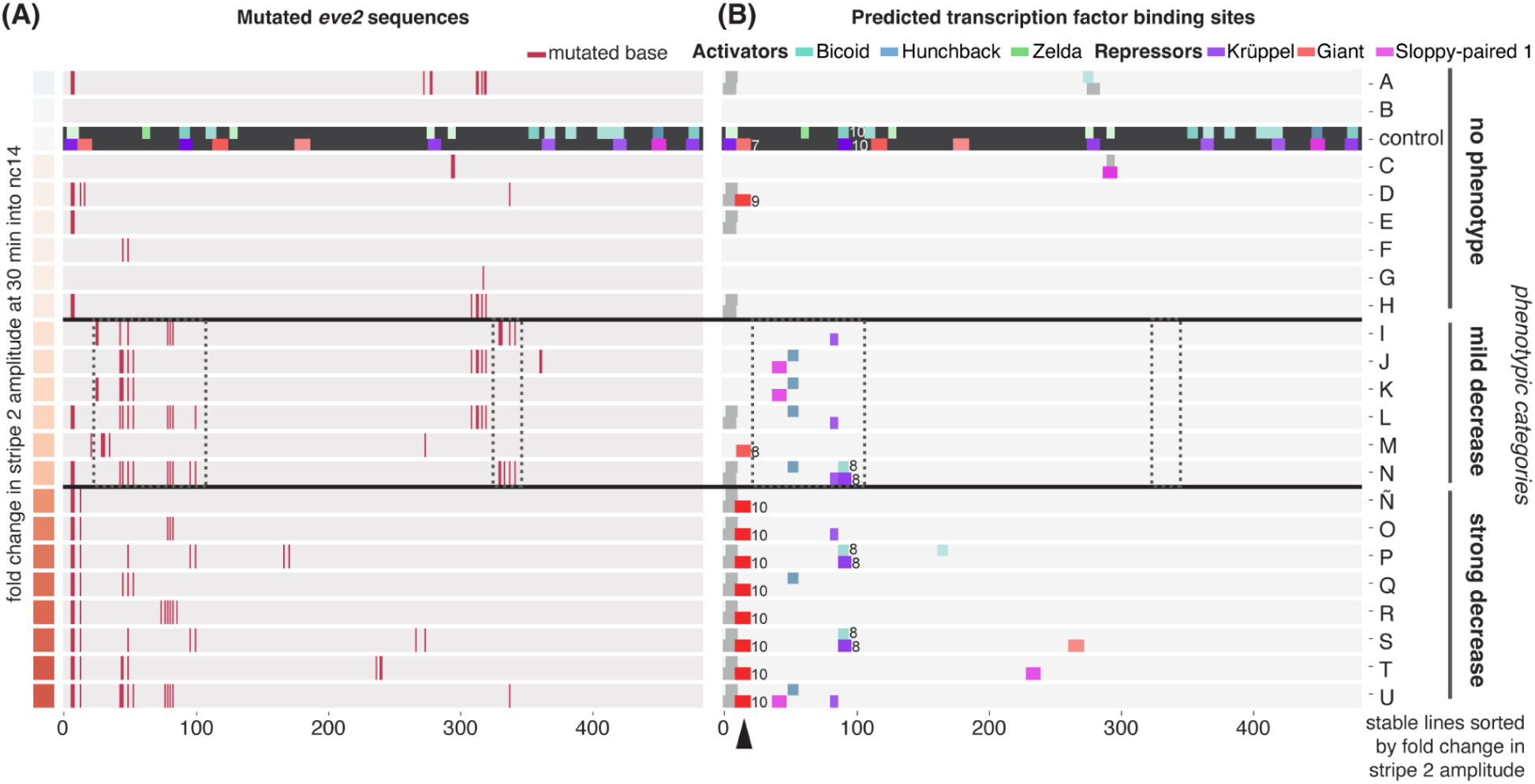
Predicted changes in transcription factor binding site affinity in each mutated line. **(A)** Results of sequencing the *eve2* enhancer in each of our lines overlaid with positions where mutations were found. Lines are sorted by change in *eve2* stripe amplitude at nuclear cycle 14 (left heatmap). **(B)** Predicted changes in binding site affinity as a result of sequenced mutations reported by their PATSER score. The BAC control line, shaded in black, shows the bioinformatically predicted sites in the wild-type *eve2* sequence. Only new sites or sites that change score are shown in the mutated lines, with numbers next to the site indicating the new score. Sites that disappear (i.e., their PATSER score falls below 6) are shown as dark grey boxes. Arrowhead indicates the higher scoring Giant site in lines with four mutations on the 5’ end of the enhancer.

## Materials and Methods

### Plasmids

Plasmids were generated using standard molecular biology techniques, either in house or by Genscript.

AID12* and AID123* or EvolvR (Addgene #83260) were kind gifts from Rahul Kohli (University of Pennsylvania) and John Dueber (UC Berkeley), respectively. These proteins were cloned downstream of a *bam* promoter and 5’UTR (Chen and McKearin, 2003) and upstream of nCas9 or enCas9 separated by a linker (see sequences below), and *bam 3’UTR* by Genscript, Inc (Piscataway, NJ) . These plasmids were further modified to replace the *bam* promoter by *act5C* (González et al., 2011) or *UASz* (Deluca and Spradling, 2018) from the pUASz1.0 plasmid (DRSC #1431) and *bam 3’UTR* by *tub 3’UTR*.

Plasmids *act5C-APOBEC1-nCas-UGI* and *act5C-evoCDA1-nCas-UGI* were kind gifts from Fillip Port (German Cancer ResearchCenter, Heidelberg University, Germany) (Doll et al., 2023).

pattB-Actin5C-BE2(pActin5c-CD-dCas9m4-UDI-attB) was obtained from Addgene #104879 (Marr and Potter, 2021).

pCDF5-U6-[4xgRNA-tRNA]-GFP expressing the 4 gRNAs targeting GFP was generated by cloning the gRNAs from (Ma et al., 2016) into pCDF5 (Addgene #73914) by PCR and HiFI assembly, following established protocols (Port and Bullock, 2016). pnos-nos5UTR-eGFP-tub3UTR was subsequently digested and cloned into pCDF5-U6-[4xgRNA-tRNA]-GFP to generate pCDF5-U6-[4xgRNA-tRNA]-GFP, nos-GFP, which contains both the GFP sequences and the 4 gRNAs targeting GFP separated by an insulator.

Cloning of the individual 20 or 23bp gRNAs in pCDF5 was performed by phosphorylating, annealing and ligating the product to *BbsI* digested pCDF5, following (Port et al., 2014).

Cloning of the GFP 24xgRNAs in arrays (2 arrays of 6 gRNAs and one of 12 gRNAs) was achieved following (Port et al., 2014) as well. Briefly, primers containing gRNA sequences overhangs and sequence to amplify the gRNA scaffold and tRNA cassette were designed to assemble first 4 arrays of 6 gRNAs each. In this manner, PCR products containing pairs of gRNAs flanking a gRNA scaffold and tRNA could be assembled in a HiFi reaction (New England Biolabs) into arrays and transformed into NEB Stable bacteria (New England Biolabs) to avoid RecA mediated recombination. Recombination events were still observed so that screening of multiple colonies was needed to find the 4 correctly assembled arrays. We then attempted to amplify these arrays by PCR to then fuse the 4 arrays of 6 gRNAs into 2 arrays of 12 gRNAs, with the objective of subsequently creating one array containing all 24 gRNAs. Since these arrays were highly repetitive we were only successful in creating one array of 12 gRNA, and therefore decided to transfect the 24 gRNAs in 3 plasmids (one array of 12 gRNAs and 2 arrays of 6 gRNAs).

To clone the *eve2* 20 bp 31x gRNA arrays, the final sequence containing all gRNAs flanked by gRNA scaffold and tRNA sequences was assembled computationally and then split into 500-600 bp chunks with Golden Gate overhangs. Each fragment was synthesized by Ansa Biotechnologies and cloned into plasmids ready to be assembled into the pCDF5 plasmid in a Golden Gate reaction, and transformed into NEBStable bacteria (New England Biolabs). Because of the difficulty in assembling highly repetitive fragments even in RecA- bacteria, we obtained plasmids missing long fragments of gRNA arrays. Nevertheless we obtained two long arrays that contained gRNAs [1-16, 29, 30, 31] and [1, 2, 3, 17-31] and were stable once isolated. Therefore gRNAs 1, 2, 3, 29, 30 and 31 were duplicated in both plasmids but we were able to obtain all the designed gRNAs.

To clone the *eve2* 23 bp 31xgRNA arrays, the final sequence containing all gRNAs flanked by gRNA scaffold and tRNA sequences was assembled computationally into two plasmids containing gRNAs 1-16 and 17-31. The 5-6kb insert for each gRNA set was synthesized by Ansa Biotechnologies and then cloned into the pCDF5 plasmid in a Golden Gate reaction using BbsI sites. The reaction was transformed into NEBStable bacteria (New England Biolabs) and all colonies screened contained the full repetitive array without recombination.

All used primers can be found in Supplementary Table 1.

All sequences for the plasmids developed or used during this study and sequences with annotated gRNAs can be found in this Benchling repository: benchling.com/juliafs/f_/NXjwqXma-optimizing-flybraries-paper/

### Cell culture

Kc-nls-GFP cells (DGRC #227, Kc167-PP-93E (nls-GFP), (Cherbas et al., 2015)) were maintained in cell media HyClone CCM3 and split twice a week to a density of 1.5 million cells/ml.

For the different experiments, Kc-nls-GFP cells were transfected with various nCas9 constructs driven by the *act5C* or *pMT* promoter together with plasmids expressing different gRNAs. In order to generate stable populations where mutating enzymes could be expressed long term, we also co-transfected an MTX-resistance plasmid (p8HCO - DGRC #1003), aiming to select cells that had integrated the co-transfected plasmids with the MTX resistance (Hannon and Eisen, 2024; Rebay et al., 1991).

Since integration rates are low and it can take up to 2 months to select stable populations, we aimed to see increased levels of mutations over time even if stable populations were not completely generated. If the mutagenic system was highly efficient, we could have also expected to see mutations saturating early on regardless of MTX selection. We thus decided to collect cells after 1, 2 and 4 weeks to quantify mutations introduced at each timepoint. However, very few cells survived MTX treatment, and mutation levels barely increased after MTX treatment or even decreased. We therefore assumed that MTX selection failed to work, but nevertheless sequenced samples at the different timepoints. Although the mutagenic enzymes were no longer active by week 4, we still expected to detect all the mutations generated in the genome while they were active.

In each condition, up to 750 ng of total DNA (containing the mutagenic enzyme and/or gRNA plasmid and the MTX resistance plasmid) was transfected in 0.75 million cells using the Effectene Transfection Reagent from Qiagen (cat. No 301425) in 24-well plates. The DNA was diluted in 75ul of EC buffer, mixed with 6ul of enhancer vortexed, and incubated at room temperature for 5 minutes. Afterwards, 8ul of Effectene reagent was added, vortexed, and incubated at room temperature for 15 minutes. The solution was then mixed and added directly to the cells. The cells were incubated for several weeks and harvested for genomic DNA extraction. Methotrexate (MTX, Sigma-Aldrich M8407) was added at 0.1 µg/mL from week 1 post-transfection and half of the cells were kept in conditioned media with MTX. At the indicated timepoints (1, 2, or 4 weeks post-transfection) cells were harvested and genomic DNA extracted. In cells transfected with pMT plasmids, a low concentration of copper (1mM CuSO4) was added one day after transfection and cells were kept until harvesting at day 8 post-transfection.

To test mutagenesis of *eve2* in cell culture, S2-attP40 cells (Mariyappa et al., 2021) cells were transfected with plasmids expressing act5C-evoCDA1-nCas9-UGI and U6-eve2-gRNAs[1-16, 29, 30, 31] and U6-eve2-gRNAs[1, 2, 3, 17-31], or with only the AID-nCas9 or gRNA plasmids for controls. Up to 1 µg of DNA was transfected using Effectene Transfection Reagent from Qiagen (cat. No 301425) per million cells in 12 well plates. DNA was harvested one week post transfection.

### Fly husbandry

*Drosophila melanogaster* flies were grown and maintained on Fly Food B from LabExpress (Ann Arbor, MI), of the following ingredients: Agar 0.56%, Cornmeal 6.71%, Inactivated Yeast 1.59%, Soy Flour 0.92%, Corn Syrup 7.0%, Propionic Acid 0.44%, Tegosept 0.15%.

Embryos were collected on apple juice agar plates (75% water, 25% apple juice, 22.5 g/Ll Bacto agar, 25 g/L sucrose, 2.5%b Nipagin M mold inhibitor 20% diluted in Ethanol, CHSL protocol doi:10.1101/pdb.rec065672) with yeast paste. Animals of both sexes were used for this study.

### Fly Strains and Genetics

The following transgenic lines were created by injection into lines containing an attP landing site using phiC31 recombination by BestGene (Chino Hills, CA) or RainbowTransgenic Flies, Inc. (Camarillo, CA):

yv ; pCFD5-GFP-4sgRNA-pNos-nos5UTR-eGFP-tub3UTR [VK22]

yw;+; bamP-bam5’UTR-enCas9-DNAPolI5M(EvolvR)-bam3’UTR [VK33]

yw;+; bamP-bam5’UTR-NLS-enCas9-DNAPolI5M(EvolvR)-bam3’UTR [VK33]

yw;+; bamP-bam5’UTR-enCas9-DNAPolI5M(EvolvR)-NLS-bam3’UTR [VK33]

yw;+; bamP-bam5’UTR-AID12*-nCas9-NLS-UGI-bam3’UTR [VK33]

yw;+; bamP-bam5’UTR-AID12*-nCas9-NLS-bam3’UTR [VK33]

yw;+; bamP-bam5’UTR-AID123*-nCas9-NLS-UGI-bam3’UTR [VK33]

yw;+; bamP-bam5’UTR-AID123*-nCas9-NLS-bam3’UTR [VK33]

yw;+; act5C-AID12*-nCas9-NLS-tub3’UTR [VK33]

yw;+; act5C-AID123*-nCas9-NLS-tub3’UTR [VK33]

yw;+; UASz-AID12*-nCas9-NLS-tub3’UTR [VK33]

yw;+; UASz-AID123*-nCas9-NLS-tub3’UTR [VK33]

yw;+; bamP-bam5’UTR-dCBEevoCDA1-nCas9-UGI-SV40 [VK33]

yw;+; bamP-bam5’UTR-dCBEevoCDA1-nCas9-UGI-SV40 [attP40]

yv ; pCFD5-U6-gRNAs-eve2-20bp (1-16, 29, 30, 31) [attP40]

yv ; pCFD5-U6-gRNAs-eve2-20bp (1, 2, 3, 17-31) [attP2]

yv ; pCFD5-U6-gRNAs-eve2-23bp (1-16) [attP40]

yv ; pCFD5-U6-gRNAs-eve2-23bp (17-31) [attP2]

The following lines were obtained from different sources:

*act5C-act5C5’UTR-dCBEevoCDA1-nCas9-UGI-SV40 [attP40]* from Phillip Boutros (Doll et al., 2023)

*act5C-act5C5’UTR-APOBEC1-nCas9-UGI-SV40 [attP40]* from Phillip Boutros (Doll et al., 2023)

*act5C-BE2(APOBEC1-dCas9)* (BDSC #92586)

*osk-Gal4 (w[1118]; P{w[+mC]=osk-GAL4::VP16}F/TM3, Sb[1]* (BDSC # 44242)

*bam-Ga4 (y[1] w[*] P{w[+mC]=bam-GAL4:VP16}1* (BDSC # 80579)

*vasa-Cas9 (w[*]; PBac{y[+mDint2] RFP[DsRed.OpIE2]=vas-Cas9.T2A.GFP}VK00033* (BDSC # 79006)

*BAC[eve]-MS2-y* was from (Berrocal et al., 2020)

In all experiments where mutagenesis occurred for one generation, male or female nos-GFP, U6-4xgRNAs flies were crossed with male or female EvolvR-nCas9 or AID-nCas9 flies. All F1 males and females nos-GFP, U6-4xgRNAs / + ; EvolvR-nCas9 / + or nos-GFP, U6-4xgRNAs / +; AID-nCas9 / + were placed in a cage and F2 embryos collected for DNA extraction.

In experiments where AID was left to mutate for multiple generations, stable stocks of nos-GFP, U6-4xgRNAs / (CyO) ; AID-nCas9 / (TM3) were created and maintained for multiple generations.

For the control with catalytically active Cas9, nos-GFP, U6-4xgRNAs was crossed with vasa-Cas9. F1 nos-GFP, U6-4xgRNAs / + ; vasa-Cas9 males and females were placed in a cage and F2 embryos collected for DNA extraction. Embryos could have received a GFP allele from either males or females, but only females expressed Cas9. Therefore obtained mutation rates of GFP are underestimated by half, and plots show measured mutation rates multiplied by 2 (Figure 3 - figure supplement 1A).

For the experiment to distinguish male and germline mutations, mutator parents were obtained as described above, by crossing males *yv ; nos-GFP, U6-4xgRNAs-v^+^* with females *yw ; act5C/bam-evoCDA1-nCas9-UGI-w^+^*. To obtain mutations only generated in the male germline, *yw / Y ; nos-GFP, U6-4xgRNAs-v^+^ / act5C/bam-evoCDA1-nCas9-UGI-w^+^* were crossed with *yw* females. White-eyed adult males and females (which could only be *yw ; nos-GFP, U6-4xgRNAs-v^+^ / +*) were selected and sequenced. To obtain mutations only generated in the female germline, *yw / yw ; nos-GFP, U6-4xgRNAs-v^+^ / act5C/bam-evoCDA1-nCas9-UGI-w^+^* females were crossed with *yw* males. White-eyed adults (which could be *yw ; nos-GFP,U6-4xgRNAs-v^+^ / +* or *yw ; + / +* if recombination had occurred) were selected and sequenced. Although these crosses contained a mix of flies harboring *nos-GFP,U6-4xgRNAs-v^+^* and not, GFP could only PCR-amplify from flies that harbored GFP, and which had been exposed to mutagenesis in the germline of its father.

To mutate endogenous *eve2* using different gRNA lengths and promoters, mutator fathers were obtained by crossing *act5C-evoCDA1-nCas9* or *bam-evoCDA1-nCas9* to *yv ; eve2-gRNAs-20bp [1-16, 29, 30, 31] ; eve2-gRNAs-20bp [1-16, 29, 30, 31]* or *yv ; eve2-gRNAs-23bp [1-16] ; eve2-gRNAs-23bp [17-31]*. F1 males containing the mutagenic enzyme and all gRNAs were crossed to yw females. At least 200 F2 white-eyed males (which could not contain the mutagenic enzyme) were selected and endogenous *eve2* sequenced. Since half of the *eve2* sequences in these F2 males come from *yw* flies, the obtained mutation rates would be underestimated by half. Plots comparing these mutation rates (Figure 6F Figure 6 - figure supplement 4) have been doubled to reflect this.

To mutate *eve2* contained in the BAC construct and isolate stable lines, mutator fathers were obtained by crossing *act5C-evoCDA1-nCas9 ; BAC[eve]-MS2-y* to *yv ; eve2-gRNAs-20bp [1-16, 29, 30, 31] ; eve2-gRNAs-20bp [1-16, 29, 30, 31]* or *yv ; eve2-gRNAs-23bp [1-16] ; eve2-gRNAs-23bp [17-31]*. F1 males containing the mutagenic enzyme and all gRNAs were crossed to balancer females. w^+^ males, containing *act5C-evoCDA1-nCas9* and/or mutated *BAC[eve]-MS2-y* were crossed again with balancers and stocks established in the third chromosome containing only mutated *BAC[eve]-MS2-y*.

### gDNA extraction, amplicon sequencing and library preparation

Genomic DNA from Kc or S2 cells was extracted using Qiagen’s DNeasy Blood & Tissue Kit (Cat. No. 69504) or the DNeasy Blood & Tissue QIAcube Kit (Cat. No. 69516).

Genomic DNA from embryos was extracted by blending embryos in PBS using a pestle and pestle mixer and subsequently using Qiagen’s DNeasy Blood & Tissue Kit (Cat. No. 69504).

Genomic DNA extraction from adult flies was performed by phenol:chloroform precipitation. Briefly flies were ground in 2.5ml of Tris HCl 0.1 M (pH 9.0) EDTA 0.1 M SDS 1% solution using Fisherbrand 15ml disposable tissue grinders (Cat no. 02-542-09) on ice. The mix was then incubated at 70C for 30 minutes. 10 ul of RNAse A were added and incubated at 37C for 15 minutes. 350 ul of KAc were added, inverted to mix and incubated on ice for 30 minutes. To remove debris, the mix was centrifuged for 15 minutes at 20G rpm in a table centrifuge at 4C. The supernatant was kept and then added 1:1 to phenol:chloroform (Sigma-Aldrich P1944). The mix was centrifuged for 5 minutes at 20G. The top layer was transferred to a new tube, 1:1 phenol:chloroform added again and centrifuged again. To precipitate DNA the top later was mixed with 0.7 volumes of isopropanol and centrifuged for 20 min at 20k G at 4C. The pellet was washed for 5 min in 70% Ethanol, left to dry and resuspended in 100 ul to 200 ul TE (10mM Tris-HCl 1mM EDTA).

Following DNA extraction, for Nanopore libraries, GFP was amplified using primers *CTCTTTCCCTACACGACGCTCTTCCGATCTtgcttcagccgctaccccgaccacatgaag* (lower case indicating the GFP sequence) and *ACTGGAGTTCAGACGTGTGCTCTTCCGATCttgatgccgttcttctgcttgtcggccatg* that add sequences equivalent to NEB adaptor and NEB Ultra II Q5 polymerase (Cat no. M0544). PCR products were purified during AMXPure beads (Beckman Coulter Cat no. A63880). A second round of PCR was performed using single index Illumina primers from New England Biolabs (Cat no. E7335S, E7500S, E7710S and E7730S). The Nanopore 9.4 ligation kit (Cat no. 110) or 10.4 ligation kit (Cat no. 114) was used to ligate the nanopore adaptors and libraries were loaded on 9.4 (Cat no. FLO-MIN106D) or 10.4 (Cat no. FLO-MIN114) flow cells using the standard protocol and sequenced on MinION Mk1B devices.

To sequence mutations in *eve2* using Nanopore technology, *eve2* was PCR amplified using primers *tgccaatgcaaattgcgggcgc* and *attgcgttgcctggacctgcgc* and NEB Ultra II Q5 polymerase (Cat no. M0544). PCR products were purified during AMXPure beads. Following the NEBNext workflow, A-tails were added to the amplified products using the NEBNext Ultra II End Prep Module (Cat no. E7546), NEB Illumina Adaptors ligated using the NEBNext Ultra II Ligation Module (Cat no. E7595), and purified with AMXPure beads. NEB single end Illumina indexes (Cat no. E7335S, E7500S, E7710S and E7730S) were added in a second round of PCR with single index primers for Illumina. The Nanopore 10.4 ligation kit (Cat no. 114) was used to ligate the nanopore adaptors and libraries were loaded on 10.4 (Cat no. FLO-MIN114) flow cells using the standard protocol and sequenced on MinION Mk1B devices.

### Analysis of sequenced Nanopore libraries

Basecalling was performed using guppy with GPU settings and ‘super-accurate’ basecalling, in a desktop computer with an Intel(R) Xeon(R) Gold 6134 CPU @ 3.20GHz, 64GB RAM and NVIDIA Quadro P4000 graphic card. For 9.4 Nanopore flowcells the base calling command was:

*“C:\Program Files\OxfordNanopore\ont-guppy\bin\guppy_basecaller.exe” --input_path [INPUT_PATH] --save_path [SAVE_PATH] -r --config dna_r9.4.1_450bps_sup.cfg--compress_fastq --verbose_logs --device cuda:0 --num_callers 8 --chunks_per_caller 1000*

and for 10.4 flowcells:

*“C:\Program Files\OxfordNanopore\ont-guppy\bin\guppy_basecaller.exe” --input_path [INPUT_PATH]\ --save_path [SAVE_PATH] -r --config dna_r10.4.1_e8.2_260bps_sup.cfg--compress_fastq --verbose_logs --device cuda:0 --num_callers 8 --chunks_per_caller 1000*

Nanopore reads were first filtered for an average quality of 10, aligned to *GFP* or *eve2* using BioPython (with alignment scores of match = 3, mismatch = 0, gap_opening = -3, gap_extension = 0) and filtered again for an average alignment score >600 for *GFP* or 1000 for *eve2*. All samples contained a high degree of mismatches, consistent with the average versions 9.4 or 10.4 sequencing quality of Nanopore sequencing and simplex basecalling. As a result of these mismatches, the alignment parameters used were adjusted from the BioPython default ones to recover most alignments.

Mismatch and gap_extension penalties were set to 0 to not penalize reads with introduced mutations or Nanopore sequencing and basecalling errors. However, the type and location of the mismatches seemed to be consistent across samples once enough reads were averaged, suggesting that this “sequencing/base calling error background” is sequence dependent and reproducible across samples, and consistent with known Nanopore sequencing biases (Delahaye and Nicolas, 2021). We therefore subtracted the average mismatch profiles calculated from control samples from experimental samples in order to obtain mismatch and indel profiles specific to the mutagenic enzymes used (Figure 2 - figure supplement 2B).

Average mutation rates were calculated by counting the number of “mutable bases” (Cs on FWD gRNAs and Gs on REV gRNAs) that were within the 20 or 23bp of the gRNA target and dividing the average C→T or G→A mismatch percentage by this number of “mutable bases”. Histograms showing the number of mutations per read were calculated by counting the number of C→T or G→A mutations on gRNA target sequences across all samples (Figure 3C, Figure 3 - figure supplement 4, Figure 6F, Figure 6 - figure supplements 1, 2, 3 and 4). The histograms in control samples showed the expected number of mismatches due to sequencing and base calling errors, and were highly skewed towards a low (1-3) number of mismatches. Control histograms were subtracted from experimental histograms to obtain an estimation of the number of mutations per read caused by the mutagenic enzymes . Mutation profiles along gRNAs were calculated by averaging C→T and G→A mutation rates over all gRNAs used in the experiment aligned by the PAM sequence (Figure 2 - figure supplement 3, Figure 5 - figure supplement 2).

Code used for sequence analysis can be found at github.com/juliafs93/CountMutations_Flybraries

### Analysis of sequenced Illumina libraries

Illumina libraries were sequenced in a MiSeq2 instrument with a MiSeq V2 Nano kit (Illumina, San Diego CA) by the Genomics Facility at the CZI Biohub (San Francisco, CA). Libraries were sequenced paired end (2x150 cycles) and de-multiplexed reads merged using BBMerge *(Bushnell et al., 2017)*. These were the inputs for similar scripts as for Nanopore data, where reads were filtered for an average quality of 30, filtered again by alignment score to GFP and each type of mismatch calculated. The code can be obtained here: <u>github.com/juliafs93/CountMutations_Flybraries</u>

### Quantification of Mutational Variability

To quantify the degree of variability in the mutations generated, in contrast to the analysis of Nanopore libraries, mutations were considered for each individual DNA molecule (i.e., on a read by read basis). First, we determined whether GFP gRNAs 1, 2, 3 or 4 contained at least one mutation in each sequenced DNA molecule. From these measurements we calculated the average mutation rate of each gRNA across all reads (defined as the fraction of DNA molecules with at least one mutation in that gRNA position) and the distribution of the number of mutated gRNAs in each read. We then used the mutation rate of each individual gRNA to predict the distribution of the number of mutated gRNAs assuming each mutation event to be independent. This prediction was compared to our experimental results in Figure 4C.

Specifically, as an illustrative example we consider a case with 4 gRNAs each mutating with independent probabilities of leading to at least one mutation given by *p_1_*, *p_2_*, *p_3_* and *p_4_*. The proportion of reads with exactly all four gRNAs with at least one mutation is

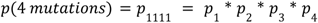

To calculate the probability of three gRNAs with at least one mutation, we need to account for all four combinations given by

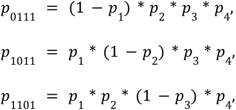

and

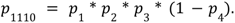

As a result, the probability of having three gRNAs with at least one mutation is

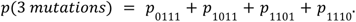

Similarly, the probability of having two or one gRNA with at least one mutation, as well as the probability of no mutations, can be calculated by accounting for all possible combinations. Each of these probabilities was used to calculate the expected percentage of reads with exactly 0, 1, 2, 3 or 4 gRNAs mutated, and then compared to the percentages obtained from simulations (Figure 4C).

As an additional quantification of the mutational variability between different DNA molecules, we also used Mutual Information theory to calculate the degree of correlation between mutated gRNAs or between mutable bases within a gRNA. This measure revealed whether two events (i.e., two gRNAs mutated on the same strand or two Cs mutated on the same gRNA) were correlated, uncorrelated or anticorrelated. For each pair of mutations we calculated the normalized pointwise mutual information (pointwise mutual information normalized by the joint self-information), which results in -1 for those two mutations never occurring together, 0 for mutations occurring independently, and +1 for complete co-occurrence of the mutations (Bouma, 2009). This mutual information is defined as

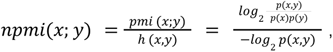

where x and y being pairs of mutated gRNA or mutable bases. Here, *p(x,y)* is the fraction of reads with the two events co-occurring and *p(x)* and *p(y)* the fraction of events with either event occurring.

The code used for the analysis of correlations can be obtained here: github.com/oliviarourke/Between-gRNA-Mutation-Correlation and github.com/oliviarourke/Within-gRNA-Mutation-Correlation

### Simulations

Simulations of AID-nCas9 mutagenesis were performed in Python by calculating probabilities of events occurring using a simple random number generator. We started from a wild type sequence and a list of gRNAs targeting this sequence. A probability of each gRNA binding to each target was calculated by generating a random number between 0 and 1. If the obtained number was lower than *p_bound_*, that gRNA would bind. If bound, AID would be allowed to mutate each C in that gRNA with a probability *p_mut_*.

In “uniform mutation profile” simulations, *p_mut_* was constant along the gRNA position. In “5’ biased mutation profile” simulations, *p_mut_*was scaled by a smoothed profile along the gRNA obtained from mutation profiles from experimental data (Figure 2 - figure supplement 3). This resulted in higher *p_mut_*at the 5’ end and lower *p_mut_* at the 3’ end, leading to a lower probability of mutating in the 3’ half of the gRNA. The same process was repeated for every “cell” (n = 200 or 1000) and for a set number of cell cyles (although the parameters are unitless and refer to probability of binding/mutation per generation). Average mutation rates and distribution of the number of mutations was calculated and plotted similarly to the experiments. The cell cycle number was set to 100 in most simulations to observe changes in mutation rates over time. A generation number of 10000 was used in some simulations to ensure mutation rates were saturated at the end of the simulation. When comparing to experiments with 4, 24 or 31 gRNAs, the cell cycle number was set to 10 to mimic the number of cell cycles occurring in cell culture or in the fly germline. We note that, in these situations, the distribution of number of mutations per DNA might not be the same after 10 generations compared to the distribution at saturation, especially for low *p_mut_*or *p_bound_* values. However this should reflect the small number of cell cycles that cells in culture or in vivo experience while mutations occur.

Code used for simulations can be found at: github.com/juliafs93/SimulatingFlybraries

### Fitting simulated data to experiments

To find which combination of *p_bound_* and *p_mut_*values better fit the experiment, we carried simulations of 10,000 reads with 10 *p_bound_* and *p_mut_* values between 0 and 1. For each of these simulations we calculated the number of mutations generated in each read, resulting in a grid of 100 histograms of the expected distribution of the number of mutations. We then compared these histograms to the ones obtained in the experiment. The best fit was then obtained by calculating the Euclidean distances between both histograms and selecting the one with the lowest value.

The code used for the fitting between simulations and experiments can be found here: github.com/oliviarourke/simulations_vs_data

### Correlation between mutation rates, gRNA score and position

We hypothesized that differences in mutation rates between gRNAs could be due to differences in binding affinity. Alternative gRNAs could be expressed at different levels depending on their distance from the promoter in our gRNA array. To shed light on this question, we calculated different biophysical parameters describing each gRNA and correlated these parameters with mutation rates in cell culture and *in vivo*. Quantified parameters were:

1. Position on the gRNA array
2. GC content
3. Binding score according to CRISPRscan
4. Binding score according to CRISPRater
5. Binding score according to Rule Set 1.

All binding scores were calculated using the package crisprDesign (github.com/crisprVerse/crisprDesign) (Hoberecht et al., 2022).

To account for overlapping gRNAs targeting the same bases, we aimed to limit which mutable bases were considered for each gRNA. Since there is a 5’ bias in mutation efficiency along each gRNA (Figure 2 - figure supplement 3), we only considered Cs in the first 10 positions of forward gRNAs and Gs in the first 10 positions of reverse gRNAs. The average % mismatch in those positions was considered the mutation rate for that gRNA.

The code used to correlate gRNA efficiency with position and binding parameters can be found at: github.com/juliafs93/be_efficiency

### smFISH experiments

Embryo fixation: Embryos were collected 2-4h post laying in apple juice plates at 25C and bleached to remove the chorion. Embryos were then transferred to heptane (Sigma #34873) and fixed for 45 min in a 1:1 heptane:formaldehyde 4% solution in PBS (Gibco #10010023) while rocking.. The bottom layer containing PBS formaldehyde was removed and replaced with methanol (Sigma #179957). The embryos were then shaken for 30” to devetillinize them. Embryos without a vitelline membrane that sunk to the bottom of the glass vial were then washed and stored at 20C in 100% methanol.

smFISH: following the smFISH protocol (Calvo et al., 2021), embryos were first rehydrated with increasing amounts of PBT (PBS 1x (Sigma) 0.05% Tween (Sigma)): 5 min per wash at 50% PBT, 75% PBT, 100% PBT. Samples were then washed 3 × 10 min in PBT, 10 min in 50% PBT 50% smiFISH wash buffer (2X SSC (Sigma #S6639), 10% deionized formamide (Sigma #F7503), in UltraPure water (Invitrogen #10977015), followed by 2 × 30 min pre-hybridization washes in smFISH wash buffer at 37 °C. All steps were performed in 2ml eppendorf tubes and solutions were removed with glass pipettes. Wash and hybridization buffers were made fresh before each experiment. 10ul of each 4μM probe mix were diluted to 80nM in 500 μl smFISH hybridization buffer (smFISH wash buffer + 10% w/v dextran sulphate (Sigma, molecular weight 6500–10,000)). Samples were left to hybridize for 14-16h in the dark at 37C. The following day samples were washed 4 × 15 min in smFISH wash buffer at 37 °C, then 3 × 10 min in PBT at room temperature. Embryos were then mounted between two coverslips with ProLong Antifade Diamond mounting medium (Invitrogen #P36961). Probes were designed to target the intronic regions of the *yellow* transcript contained in the BAC construct as described in (Calvo et al., 2021) and synthesized containing a FLAP sequence. Probes were annealed through the FLAP to another primer bound to Alexa546 (Calvo et al., 2021). Probe sequences, including target and FLAP sequences, can be found in Supplementary Table 2.

Imaging was performed in a Zeiss LSM800 confocal microscope, using a 63x oil objective. Briefly, 4096 x 2024 px images at 0.396μm/pixel resolution were collected in 8x4 tiles. The confocal scanning was bidirectional and the pinhole set to 44μm. 20-30 z stacks of 1μm each were collected.

### Image analysis and gene expression quantification

To obtain expression profiles and other transcription parameters from smFISH images, we focused on the bright smFISH puncta corresponding to nascent sites of transcription. First, images are rotated to set the anterior posterior axis horizontally and the anterior side to the left and the maximum intensity projection obtained. A Gaussian filter of 10 pixels is then applied (Figure 7 - figure supplement 8A) and Otsu threshold calculated to mask the shape of the embryo (Figure 7 - figure supplement 8B). A difference of Gaussians filter (sigmas 1 and 2 pixels) is then applied to the maximum projected image to identify the spots (Figure 7 - figure supplement 8C). The position of all spots is then used to manually draw the 7 stripes, and assign spots to each of them (Figure 7 - figure supplement 8D, E, F). Similarly to (Berrocal et al., 2020), the stripes were then straightened to account for the shape of the embryo. Briefly, the position of all spots assigned to each stripe was fitted to a quadratic function (Figure 7 - figure supplement 8G). The average x-position of all spots for one stripe was set as the center of the stripe. Then, the y-position was kept for each spot and the x-position shifted from the new center of the stripe by the difference between the original x-position and the parabola. In this manner, each spot kept the same distance from the center of the stripe as it did from the original stripe (the fitted parabola) while being straightened vertically (Figure 7 - figure supplement 8H).

The intensity at the centroid of each spot at the new spot location was then used to calculate the expression profile. Briefly, fluorescent intensities were added along the x-axis, resulting in a noisy expression profile. Positions with no spots were replaced by 0s and then a one dimensional gaussian filter of 20 pixels was applied to obtain the expression profiles. The function *find_peaks* from the *SciPy* package was then applied to this profile to obtain the maxima of 7 peaks (Virtanen et al., 2020) . The whole profile was rescaled in x by fixing the positions of the maxima corresponding to peaks 1, 3 and 7, and in y by normalizing to the average of all peaks excluding the one corresponding to stripe 2 (Figure 7 - figure supplement 8I). In this manner we accounted for differences in x scaling related to the orientation of the embryo and normalized internally to account for differences in signal intensity across embryos. We note that, as a result, figures showing expression profiles do not quantitatively report on x-axis position.

Stripe two amplitude was defined as the height of the second peak after normalization. In addition to amplitude, we also quantified the stripe 2 width (defined as mean + 2 SD of the X positions of stripe 2 spots), average spot intensity (stripe 2 normalized to stripe 3) and fraction of active nuclei (number of spots in stripe 2 normalized to stripe 3).

Since these parameters changed over time, to calculate changes in expression patterns we first had to classify control and mutant embryos in buckets corresponding to different timepoints. While differences were clear enough to perform manual classification into timepoints, based on the width and relative intensities of the stripes, we automated this process by fitting each embryo to a temporal bucket based on the width for all stripes. Briefly, in control embryos stripes narrow over time and stripe 7 is the widest at early timepoints (Figure 7 - figure supplement 8J). For each embryo, we found the temporal bucket where the distribution of width vs stripe number was a better fit, i.e. the least Euclidean distance. Obvious misclassifications were corrected manually.

The same process was performed for all mutant and control embryos to classify into timepoints. To quantify changes in phenotype, for each mutant embryo we calculated the fold change in a given parameter (stripe 2 amplitude, width, spot intensity or fraction of active nuclei) by dividing by the average of controls at the same timepoint.

The code used for all image analysis is available at: github.com/JialingFang/eve_BAC_analysis

### Genotyping stable lines

DNA was extracted from individual adult males using the standard fly squish protocol (Gloor and Engles, 1992). Briefly, flies were squished with a tip in an eppendorf tube containing 50ul of squishing buffer (10 mM Tris-HCl pH8, 1 mM EDTA, 25 mM NaCl, 200 g/ml Proteinase K). The mix was then incubated at 37C for 30 min and 95C for 3 min to inactivate Proteinase K. 1ul of this mix was used to PCR amplify and genotype *eve2* sequences present in each stable line.

To genotype mutations in endogenous *eve2,* PCRs were performed using 1ul of squished fly extract, primers *tgccaatgcaaattgcgggcgc* and *attgcgttgcctggacctgcgc*, and JumpStart REDTaq ReadyMix Reaction Mix (Sigma Cat no. P0982). Amplified products were purified using AMXPure beads and sequenced by Sanger Sequencing at the Berkeley DNA sequencing facility using primer *ttgcgggcgcaatataaccc*.

To genotype mutations in BAC eve2, a first PCR was performed using the same FWD primer as for endogenous *eve2* (*tgccaatgcaaattgcgggcgc*) and a primer that would only amplify the *yellow* coding sequence present in the BAC (*cggcgtcaccaaggtgatcaggg*) using JumpStart REDTaq ReadyMix Reaction Mix. 1ul of this reaction was used for a second PCR using primers for *eve2 ttgcgggcgcaatataaccc* and *attgcgttgcctggacctgcgc.* Amplified products were purified using AMXPure beads and sequenced by Sanger Sequencing at the Berkeley DNA sequencing facility using primer *ttgcgggcgcaatataaccc*.

### Finding putative binding sites with PATSER

We used a modified version of SiteOut (Vincent et al., 2016), to run PATSER (Hertz and Stormo, 1999) on *eve2* and predict binding sites based on the position weight matrices of selected transcription factors. Used PWMs, selected from SiteOut, are included in the GitHub repository below. Bicoid, Hunchback and Krüppel PMWs were from (Bergman et al., 2005), and Sloppy paired 1 PWM from (Noyes et al., 2008). Following parameters from (Hare et al., 2008), GC% parameter was set to 0.43 as that is the average in *Drosophila* and sites with a PATSER score above 6 and *p value* lower than *e^-6^*(0.01 for Zelda) were selected.

The code used to run PATSER on wild type and mutated *eve2* sequences is available here: github.com/juliafs93/PATSER-eve2

## Acknowledgements

We would like to thank Steve Quake for the initial discussion that launched this project as well as the Phillips lab for their intellectual and technical support as we develop this technology. We also acknowledge John Dueber and Fillip Port, as well as their lab members, for extensive discussions and reagents. We thank members of the Garcia and Eisen labs for helpful discussion and feedback. J. F-S. was supported by EMBO (ALTF 1267-2020) and HFSP (LT000310/2021-L) Long Term Postdoctoral Fellowships. Y. D-T was supported by the PREP UC Berkeley/CND Post Baccalaureate Program (funded by the US National Institutes of Health via R25GM140276). M.A.T. was supported by the NSF Graduate Research Fellowship. J.D and O.R. were supported by the Cal-Bridge Summer Program. C. M. was supported by a Chancellor’s Fellowship from UC Berkeley’s Office for Graduate Diversity. H. G. G. was supported by NIH Awards R01GM139913, R01GM152815 and R35GM158200, by the Koret-UC Berkeley-Tel Aviv University Initiative in Computational Biology and Bioinformatics, by a Winkler Scholar Faculty Award, by the Chan Zuckerberg Initiative Grant CZIF2024-010479, and by the Miller Institute for Basic Research in Science, University of California Berkeley. H.G.G. is also a Chan Zuckerberg Biohub Investigator (Biohub – San Francisco). M.B.E. was supported by the Howard Hughes Medical Institute. J. F-S. and H. G. G were also supported by a Biohub Imaging Residency.

